# Vγ9Vδ2 T cells recognize butyrophilin 2A1 and 3A1 heteromers

**DOI:** 10.1101/2023.08.30.555639

**Authors:** Thomas S. Fulford, Caroline Soliman, Rebecca G. Castle, Marc Rigau, Zheng Ruan, Olan Dolezal, Rebecca Seneviratna, Hamish G. Brown, Eric Hanssen, Andrew Hammet, Shihan Li, Samuel J. Redmond, Amy Chung, Michael A. Gorman, Michael W. Parker, Onisha Patel, Thomas S. Peat, Janet Newman, Andreas Behren, Nicholas A. Gherardin, Dale I. Godfrey, Adam P. Uldrich

**Affiliations:** Department of Microbiology & Immunology at the Peter Doherty Institute for Infection and Immunity, University of Melbourne, Parkville, Victoria, 3010, Australia; Institute of Molecular Medicine and Experimental Immunology, Rheinische Friedrichs-Wilhelms University of Bonn, D-53127 Bonn, Germany; Biomedical Manufacturing, Commonwealth Scientific and Industrial Research Organisation, 343 Royal Parade, Parkville, Victoria, 3052, Australia; Ian Holmes Imaging Centre, Bio21 Molecular Science and Biotechnology Institute, University of Melbourne, Parkville, Victoria, 3052, Australia; Department of Biochemistry and Pharmacology, Bio21 Molecular Science and Biotechnology Institute, University of Melbourne, Parkville, Victoria, 3052, Australia; ARC Industrial Transformation Training Centre for Cryo-electron Microscopy of Membrane Proteins, University of Melbourne, Parkville, Victoria, 3052, Australia; CSL Limited, Bio21 Molecular Science and Biotechnology Institute, Parkville, Victoria 3052, Australia; St. Vincent’s Institute of Medical Research, Fitzroy, Victoria, 3065, Australia; The Walter and Eliza Hall Institute of Medical Research, Parkville, Victoria, 3052, Australia; Department of Medical Biology, University of Melbourne, Parkville, Victoria, 3052, Australia; Olivia Newton-John Cancer Research Institute, and School of Cancer Medicine, La Trobe University, Heidelberg, Victoria, 3084, Australia; Department of Medicine, University of Melbourne, Parkville, Victoria, 3010, Australia; Cancer Immunology Program, Peter MacCallum Cancer Centre, Melbourne, Australia

## Abstract

Butyrophilin (BTN) molecules are emerging as key regulators of T cell immunity, however, how they trigger cell-mediated responses is poorly understood. Here, the crystal structure of a gamma-delta T cell receptor (γδTCR) in complex with BTN member 2A1 (BTN2A1) revealed that BTN2A1 engages the side of the γδTCR, leaving the apical TCR surface bioavailable. We reveal that BTN3A1 is a second γδTCR ligand, that co-engages γδTCR via binding to this accessible apical surface. BTN2A1 and BTN3A1 also directly interact with each other in cis, and structural analysis revealed formation of W-shaped heteromeric multimers. This BTN2A1–BTN3A1 interaction involved the same epitopes that BTN2A1 and BTN3A1 each use to engage γδTCR; indeed, either forced separation or locking together of BTN2A1 and BTN3A1 resulted in enhanced or abrogated γδTCR interaction, respectively. Our findings reveal a new paradigm in immune activation, whereby γδTCRs recognize dual epitopes on BTN2A1 and BTN3A1 complexes.

## Main text

T cell receptor (TCR) recognition of antigen is a central event in immunity. Alpha-beta (αβ) T cells become activated following recognition of peptide fragments in complex with major histocompatibility complex molecules (pMHC), which are sensed by somatically rearranged T cell receptors (αβTCRs) derived from the TCRα (*TRA*) and TCRβ (*TRB*) gene loci. By contrast, gamma-delta (γδ) T cells represent a separate lineage of MHC-unrestricted T cells that express rearranged antigen (Ag) receptors derived from the TCRγ (*TRG*) and TCRδ (*TRD*) gene loci. γδ T cells play a key role in the priming and effector phases of immunity to infectious diseases as well as in tissue surveillance ^1^. In humans, the majority of circulating γδ T cells express a semi-invariant Vγ9Vδ2^+^ (TRGV9–TRGV2) γδTCR that confers reactivity to a distinct class of non-peptide Ag, termed phosphoantigens (pAg), which are metabolic intermediates in the biosynthesis of isoprenoids ^2,3^. There are two classes of pAg; those derived from the non-mevalonate pathway such as (E)-4-hydroxy-3-methylbut-2-enyl pyrophosphate (HMBPP), found in bacteria and apicomplexan parasites, and those derived from either the mevalonate or non-mevalonate pathways, such as isopentenyl pyrophosphate (IPP), found in all classes of life, including vertebrates. Both ‘foreign’ HBMPP and ubiquitous IPP pAgs are stimulatory for γδ T cells to differing degrees, and facilitate potent anti-microbial and anti-tumor immunity, respectively ^4,5^.

Butyrophilin (BTN) and butyrophilin-like (BTNL) molecules are a family of surface-expressed transmembrane proteins that are typically comprised of extracellular immunoglobulin-superfamily variable (IgV)- and constant (IgC)-like domains, as well as an intracellular B30.2 domain. In certain combinatorial pairs, BTN and BTNL molecules support the activation of discrete γδ T cell subsets. For instance, BTNL3 and BTNL8 are expressed by gut epithelia and cooperate to facilitate the activation of Vγ4^+^ γδ T cells ^6^. Likewise, in mice, Btnl1 and Btnl6 facilitate the activation of gut-resident Vγ7^+^ γδ T cells ^6,7^, and the Btnl family members Skint1 and Skint2 are important for the development and function of skin-resident Vγ5Vδ1^+^ dendritic epidermal T cells (DETCs) ^8,9^. Unlike all other modes of T cell antigen recognition in which antigens are presented on the cell surface for recognition via the TCR, BTN member 3A1 (BTN3A1) sequesters pAg via a positively charged pocket within its intracellular B30.2 domain, which is an essential step in the initiation of Vγ9Vδ2^+^ T cell activation ^10^. Together, BTN member 2A1 (BTN2A1) and BTN3A1 mediate γδ T cell responses to pAg ^11,12^. Thus, BTN molecules have emerged as important regulators of γδ T cell-mediated immunity and do so as heteromeric pairs.

Recently, reports have shown that BTN molecules can directly bind the γδTCR, including BTNL3, which interacts with human Vγ4, and mouse Btnl1 which interacts with Vγ7 ^7,13^. Likewise, BTN2A1 is a binding partner for human Vγ9 domain ^11,12^. However, whilst there appears to be an evolutionarily conserved mode of recognition between these systems based on overlapping regions of importance within Vγ domains ^7,9,11–13^, the molecular mechanism that governs BTN ligand recognition by γδTCR heterodimers is poorly understood, and structural information is lacking. Furthermore, in the case of BTN2A1, the mode of ligand recognition is unclear since one study found a dependence on the side of the Vγ9 domain ^11^, whereas other studies have additionally implicated the hypervariable-4 (HV4) loop ^12,13^ or the complementarity-determining region (CDR) 3δ loop ^14^, both located on the apical surface of the γδTCR.

In addition to Vγ9Vδ2^+^ TCR binding to BTN2A1, mutational analysis revealed the presence of a second ligand-binding region that was distinct from the BTN2A1 binding site, which is also essential for immune reactivity to pAg ^11,15^. In one model, BTN3A1 could represent the second ligand, however, a direct BTN3A1–γδTCR interaction has not been established. An alternative model proposes that an undefined ligand binds γδTCR, and BTN3A1 engages another undefined receptor expressed by Vγ9Vδ2^+^ T cells, rather than the γδTCR itself ^14^. Finally, although BTN2A1 and BTN3A1 are constitutively expressed by immune cells, γδ T cell activation only occurs following pAg challenge, or cross-linking with agonist anti-BTN3A antibodies such as clone 20.1 ^16^. Thus, a molecular switch that involves either a conformational change and/or remodelling of BTN complexes on the cell surface of APCs appears to initiate γδ T cell responses. However, the nature of these regulatory mechanisms is unclear.

Here, we report the crystal structure of the BTN2A1–γδTCR complex, which represents the first structural example of a TCR engaging a ligand outside of an MHC/MHC-like family member, wherein BTN2A1 binds to the side of Vγ9. Furthermore, we reveal the identity of the second ligand for the Vγ9Vδ2^+^ TCR as BTN3A1, which co-binds γδTCR along with BTN2A1. We also report the structure of a heteromeric BTN complex, namely BTN2A1–BTN3A1, in which the interface between the disparate BTN molecules involves the same epitopes that both BTN2A1 and BTN3A1 utilize to co-bind the γδTCR. Indeed, forced separation of BTN2A1 and BTN3A1 enhanced binding to γδTCR, while conversely, locking them together abrogated this interaction. Thus, the association between BTN2A1 and BTN3A1 in cis might serve as a regulatory mechanism by burying the TCR-binding determinants via sequestration with each other. Accordingly, we provide evidence for a two-ligand sensing system that, upon triggering, acquires affinity for γδTCR through the exposure of cryptic epitopes, and coupled with an apparent conformational change to BTN3A1, can then co-bind the Vγ9Vδ2^+^ TCR.

## Results

### BTN2A1 engages the side of γδTCR

To understand the molecular mode of BTN engagement by γδTCR, we solved the crystal structures of BTN2A1 ectodomain either alone (‘apo’), or in complex with Vγ9Vδ2^+^ γδTCR, which diffracted to 3.6 Å (anisotropic range 3.6 Å – 4.3 Å) and 2.1 Å resolution, respectively (**Extended Data Table 1**). The unbiased electron density was clear in both structures (**Extended Data Fig. 1A**). BTN2A1 is reported to exist on the cell surface predominantly as a homodimer, which is stabilized by a membrane-proximal interchain disulfide bond ^12^. Our BTN2A1 construct lacked the terminal Cys residue responsible for this disulfide bond and consequently behaved as a monomer in solution (**Extended Data Fig. 1B**). Nonetheless, the apo form of BTN2A1 contained five copies in the asymmetric unit, arranged as two V-shaped homodimers (‘V-dimers’) (**Fig. 1A**) that align head-to-tail to one-another, with the fifth copy also forming a separate V-dimer via crystallographic symmetry. This V-dimer is broadly reminiscent of the BTN3A1 V-dimer ^17^, although the BTN2A1 V-dimers formed at an angle of 59°, which is wider than BTN3A1 V-dimers (49°), and the BTN2A1 V-dimers were also twisted by 25° relative to BTN3A1 V-dimers ^17^ (**Fig. 1B** and **Extended Data 1C**). The V-dimer was characterized by a small interface dominated by a limited array of primarily non-polar interactions including three π-mediated interactions, with a buried surface area (BSA) of ∼430 Å^2^ per molecule (**Extended Data Fig. 1D** and **Extended Data Table 2**). A head-to-tail dimer of BTN2A1 was also observed in both the apo structure (**Fig. 1B**) and BTN2A1– γδTCR complex (**Extended Data Fig. 1E**), although the latter only involved the unliganded BTN2A1, via crystallographic symmetry, because the head-to-tail footprint overlapped with the γδTCR binding site. The head-to-tail dimer had a larger BSA of ∼1180 Å^2^ per molecule compared to the V-dimer (**Extended Data Fig. 1D** and **Extended Data Table 3**), and could potentially form via a cis or a trans interaction (**Fig. 1B**), akin to the purported BTN3A1 head-to-tail homodimer ^17^.

**Figure 1.**
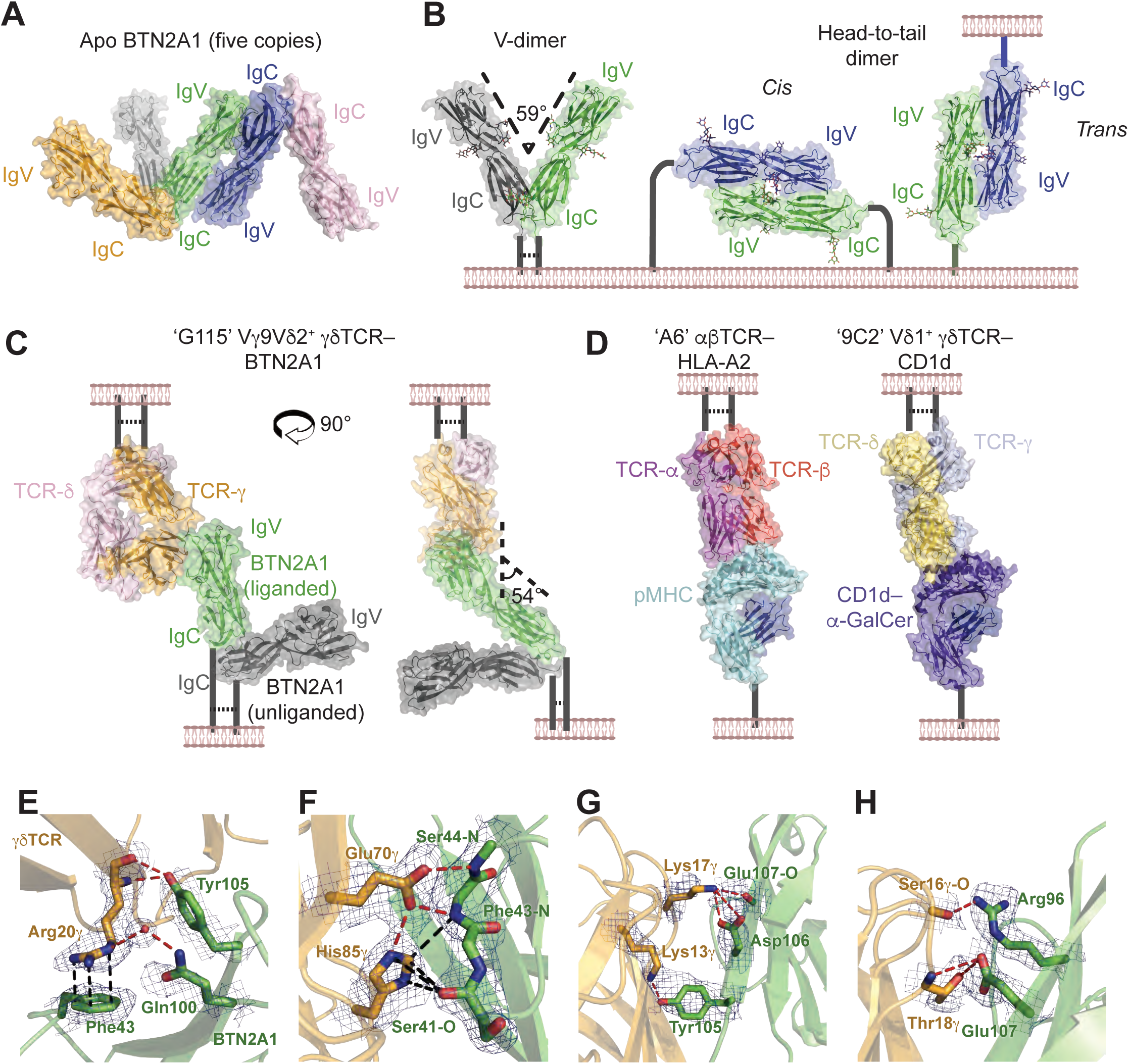
BTN2A1 engages the side of Vγ9. (**A**) Surface and cartoon representation of the apo-BTN2A1 crystal structure. BTN2A1 copies, orange, gray, green, blue and light pink. (**B**) The BTN2A1 V-dimer (left) and cis (middle) or trans (right) interpretation of the head-to-tail homodimer. Color scheme as in (A). (**C**) Surface and cartoon representation of the BTN2A1– Vγ9Vδ2^+^ TCR clone G115 crystal structure. G115 TCRδ, pink; G115 TCRγ, orange; liganded BTN2A1, green; unliganded BTN2A1, gray. (**D**) Comparison with a representative pHLA Class I–αβTCR (left, PDB code 1QSE ^29^) and CD1d–α-GalCer–clone 9C2 Vγ5Vδ1^+^ TCR (right, PDB code 4LHU ^30^). TCRα, dark pink; TCRβ, red; HLA-I, light blue; β2-microglobulin, dark blue; 9C2 TCRδ, yellow; 9C2 TCRγ, pale blue, CD1d, purple. Molecular contacts between the Vγ9Vδ2^+^ clone G115 TCR (orange) and BTN2A1 (green) ectodomains showing the (**E**) Arg20γ, (**F**) Glu70γ and His85γ, (**G**) Lys13γ and Lys17γ, (**H**) Ser16γ and Thr18γ side and/or main chains and their BTN2A1 contacts as sticks. H-bonds and salt-bridges, red; VDW and cation-π, black; 2mFo-DFc electron density composite omit map contoured at 1σ, blue mesh.

The asymmetric unit of the complex contained two copies of BTN2A1, also arranged as a V-dimer, which was similar to the apo BTN2A1 V-dimers (**Extended Data Fig. 1F**). One of the BTN2A1 copies liganded to the γδTCR, and the other one was unliganded (**Fig. 1C**). BTN2A1 engaged the side of the γ-chain, binding to the Vγ9-encoded IgV domain, jutting out at an angle of ∼54°, which starkly contrasted αβTCR engagement of pMHC, or γδTCR recognition of CD1d (**Fig. 1D**). Previous studies implicated the hypervariable region 4 (HV4) loop, also known as the DE loop of Vγ9, as well as the CDR3δ loop, in binding BTN molecules ^12–14^. The crystal structure revealed that the BTN2A1 binding site on Vγ9 was distal to both the CDR and HV4 loops (>7 Å and >9 Å separation, respectively), and instead left the entire apical surface of the γδTCR solvent exposed (**Fig. 1C**). The outer face of the Vγ9 germline-encoded β-sheet formed by the A, B, D and E β-strands (ABED face) mediated binding to the β-sheet encoded by the C, F and G β-strands (CFG face) of the BTN2A1 IgV domain (**Extended Data Fig. 1G**), with interactions by all these strands (**Extended Data Fig. 1H**). The γδTCR buried 470 Å^2^ upon ligation, and BTN2A1 buried 480 Å^2^ (BSA of total interface = 950 Å^2^) (**Extended Data Fig. 1H)**, which is approximately half of a typical αβTCR-pMHC complex, with the molecules anchored together by fourteen H-bonds or salt bridges **(Extended Data Table 4**). On the γδTCR, the B-, D- and E-strands of Vγ9 contributed 57%, 17% and 11% of the BSA, respectively, whereas the CC′-loop, F- and G-strands of BTN2A1 contributed 35%, 15%, and 44%, respectively. Within the BTN2A1 interface, the aromatic residues Phe43, Tyr98 and Tyr105 made molecular interactions, with minimal conformational change between the apo and liganded states, indicative of a lock- and-key mode of binding (**Extended Data Fig. 1I**). Ser41, Gln42, Phe43 and Ser44 formed the CC′ loop (**Extended Data Fig. 1G**), and their involvement is consistent with the over-representation of this loop in other IgV-mediated interfaces ^18^. Of note, the aromatic side chain of Phe43 sat planar to the guanidinium moiety of the Arg20γ side chain (**Fig. 1E**), facilitating a cation-π interaction with a predicted electrostatic binding energy of -4.6 kcal/mol.

Arg20γ also formed a water-mediated H-bond with Gln100 of BTN2A1, along with main chain-mediated H-bonds to the Tyr105 side chain hydroxyl group (**Fig. 1E**), providing a structural basis for the importance of Arg20γ in BTN2A1-binding ^11^. Likewise, we have shown previously that mutations to Glu70γ and His85γ abrogate BTN2A1 reactivity ^11^, and these residues were interconnected by a H-bond, and also bound BTN2A1. Here, Glu70γ H-bonded to the Phe43 and Ser44 main chains, and His85γ made Van der Waal (VDW) contacts with Ser41, Gln42 and Phe43 on BTN2A1 (**Fig. 1F**). Further contacts were made by Lys13γ within the A-strand of Vγ9, which H-bonded to Tyr105, and Lys17γ within the B-strand of Vγ9, forming a salt bridge with Asp106 (**Fig. 1G**). The adjacent Thr18γ H-bonded with Glu107, and Ser16γ H-bonded to the Arg96 side chain (**Fig. 1H**). Accordingly, BTN engagement by γδTCR represents a fundamentally unique mode of ligand recognition by the immune system.

### BTN3A1 modulates Vγ9Vδ2^+^ TCR tetramer interaction

Given the high bioavailability of the apical surface of the Vγ9Vδ2 TCR when liganded to BTN2A1, we hypothesized that Vγ9Vδ2 TCR co-binds a second ligand. Since BTN3A1 intracellular domain binds pAg ^10^, we first examined whether soluble BTN3A1 ectodomain could directly bind γδTCR by probing Vγ9Vδ2 TCR-transfected *BTN2A^KO^*.*BTN3A*^KO^ HEK293T cells, which lack endogenous *BTN2A1*, *2A2*, *2A3p*, *3A1*, *3A2* and *3A3,* with BTN3A1 ectodomain tetramers. Consistent with an earlier report ^10^, we did not detect any staining (**Extended Data Fig. 2A**). We next tested whether mouse NIH-3T3 fibroblasts, which lack human BTN or BTNL molecules and are inherently incapable of mediating Vγ9Vδ2^+^ T cell activation by pAg, could bind Vγ9Vδ2^+^ TCR tetramer. These cells were transfected with either full-length human BTN2A1 or BTN3A1. Compared to BTN2A1^+^ NIH-3T3 cells, which bound all Vγ9Vδ2^+^ TCR tetramers (clones TCR3, TCR6, TCR7 and G115; **Fig. 2A**), BTN3A1^+^ cells showed little, if any, staining (**Fig. 2B**, **empty bars; Extended Date Fig. 2B, black dots**). Previous studies showed that crosslinking of BTN3A1 on the surface of APCs with anti-BTN3A antibody (mAb clone 20.1) converts BTN3A1 into a stimulatory form that can activate Vγ9Vδ2^+^ T cells in a way that mimics pAg challenge ^16,17^. Conversely, a separate anti-BTN3A mAb (clone 103.2) is a potent antagonist of Vγ9Vδ2^+^ T cell reactivity to pAg ^16,17^. Strikingly, cross-linking of BTN3A1^+^ cells with agonistic mAb clone 20.1 induced clear staining with Vγ9Vδ2^+^ TCR tetramers, particularly clones TCR3, TCR7 and G115 (**Fig. 2B, half-filled bars; Extended Data Fig. 2B, red plots**). This contrasted with the antagonist anti-BTN3A mAb clone 103.2, which did not induce any Vγ9Vδ2^+^ TCR tetramer staining, nor did mAb 20.1 treatment of un-transfected or BTNL3-transfected cells (**Fig. 2B, 20%-filled bars; Extended Data Fig. 2B, blue plots**). We obtained a similar pattern of mAb 20.1-induced BTN3A1-dependent Vγ9Vδ2^+^ TCR staining using BTN3A1-transfected human *BTN2A*^KO^.*BTN3A*^KO^ HEK293T cells (**Extended Data Fig. 2C**). Furthermore, chimeric TCR tetramers comprised of a pAg-reactive Vγ9^+^ γ-chain paired with an irrelevant Vδ1^+^ δ-chain (**Extended Data Fig. 2D**) retained binding to BTN2A1^+^ cells, but not to mAb 20.1-cross-linked BTN3A1^+^ cells, indicating that unlike BTN2A1 protein interactions, the BTN3A1 association depends on Vδ2 and/or the CDR3δ loops (**Fig. 2C** and **D; Extended Data Fig. 2E**). Thus, mAb 20.1 pre-treatment of BTN3A1-transfected cells induces an interaction with Vγ9Vδ2^+^ TCR via recognition of a second ligand, herein termed ‘ligand-two’. Ligand-two association could be induced upon mAb 20.1 cross-linking of BTN3A1 in both human and mouse cell lines, and unlike the BTN2A1 interaction, this binding was dependent on the Vδ2 domain and/or the CDR3 loops, hereafter referred to as ‘epitope two’ (**Fig. 2F, cartoon inset**).

**Figure 2.**
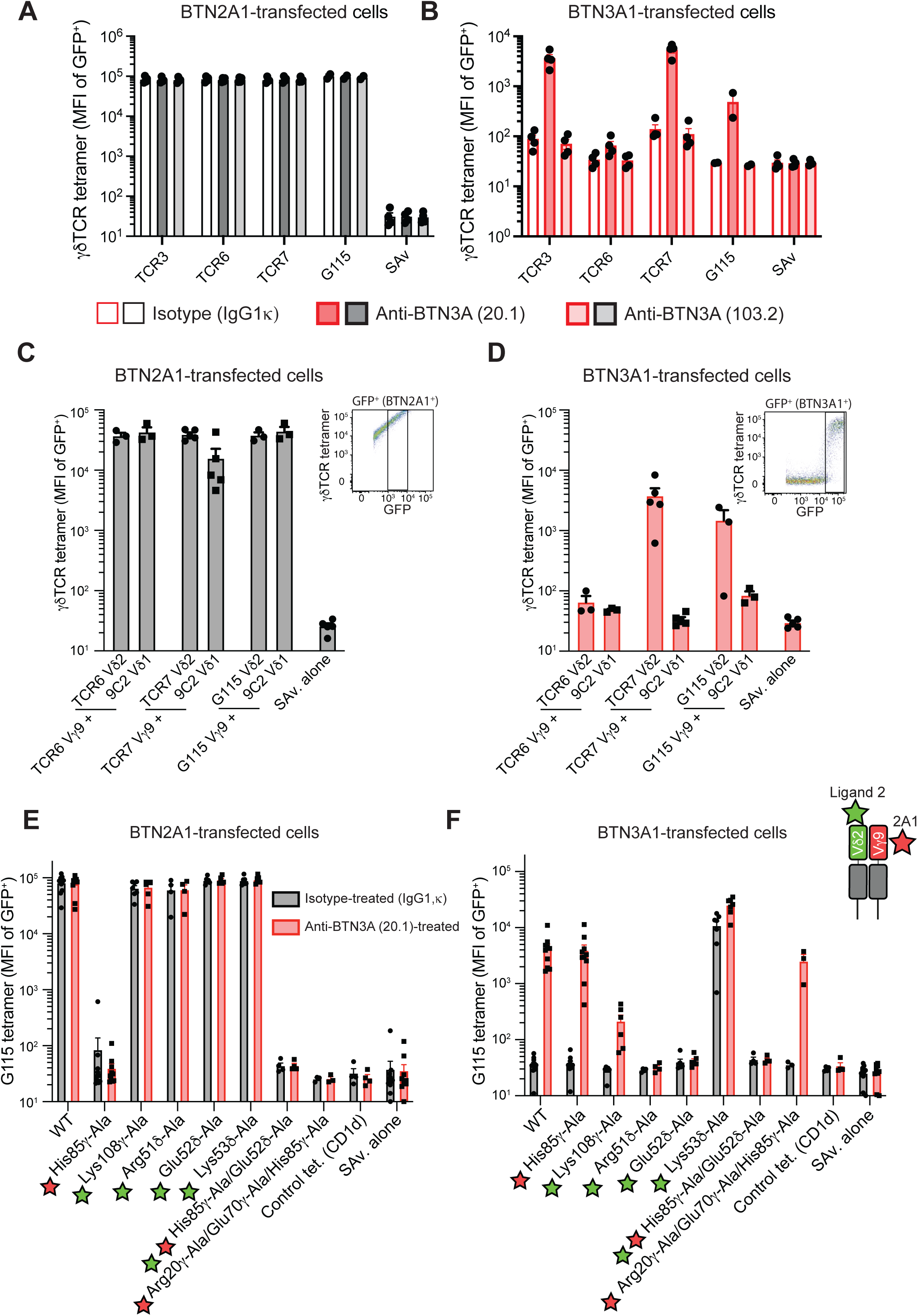
BTN3A1 supports binding to the apical surface of the Vγ9Vδ2^+^ γδTCR. MFI of chimeric γδTCR tetramer staining of (**A**) gated GFP^+^ BTN2A1-transfected or (**B**) GFP^+^ BTN3A1-transfected NIH-3T3 cells treated with isotype control, anti-BTN3A (clone 20.1) or anti-BTN3A (clone 103.2) antibodies. Graphs are presented as mean ± SEM. N ³ 3, where each point represents an independent experiment. MFI of chimeric γδTCR tetramer staining of (**C**) gated GFP^+^ BTN2A1-transfected or (**D**) GFP^+^ BTN3A1-transfected NIH-3T3 cells treated with anti-BTN3A (clone 20.1). Inset plots depict parent cell gating. Graphs are presented as mean ± SEM. N ³ 3, where each point represents an independent experiment. (**E**) GFP^+^ BTN2A1-transfected or (**F**) GFP^+^ BTN3A1-transfected NIH-3T3 cells. Cartoon inset depicts the locations of BTN2A1-epitope (red star) and the ligand-two epitope (green star). Graphs depict mean ± SEM. N ≥ 3, where each point represents an independent experiment.

### Lys53δ regulates the interaction with ligand-two

Since the ABED β-sheet of Vγ9 mediates binding to BTN2A1 (**Fig. 1C**), we tested whether the symmetrically equivalent ABED β-sheet of Vδ2 is also important in sensing pAg. Jurkat cells expressing Ala mutants within the Vδ2-encoded ABED β-sheet did not impair reactivity to zoledronate (Zol; an aminobisphosphonate that increases intracellular IPP pAg), suggesting there is no Vδ2-encoded equivalent ABED binding interface to the BTN2A1-binding domain on Vγ9 (**Extended Data Fig. 3A–C**). Since pAg-mediated γδ T cell responses depend on Arg51δ and Glu52δ, both located within the CDR2δ loop [^11,15^ and **Extended Data Fig. 3C, red residues**], we screened two additional mutations within this loop: Lys53δ-Ala and Asp54δ-Ala. Whilst Asp54δ-Ala did not appreciably affect reactivity to zoledronate, surprisingly, Jurkat cells expressing G115 Vγ9Vδ2^+^ TCR with a Lys53δ-Ala mutation (G115^Lys53δ-Ala^) exhibited spontaneous activation, indicating that this residue may have a role in dampening γδ T cell reactivity to TCR stimuli (**Extended Data Fig. 3A–C**).

To reconcile these observations with mAb 20.1-induced Vγ9Vδ2^+^ TCR tetramer staining of BTN3A1^+^ cells, we produced G115 Vγ9Vδ2^+^ TCR tetramer (hereafter referred to as G115 tetramer) with these corresponding Ala substitutions. As expected, wild-type G115 tetramer interacted with BTN2A1^+^ NIH-3T3 fibroblasts, and also with mAb 20.1-pretreated BTN3A1^+^ cells (**Fig. 2E** and **F; Extended Data Fig. 4A**). G115 tetramers with mutations at the BTN2A1 binding site (‘epitope one’), notably G115^His85γ-Ala^ and an G115^Arg20γ-Ala/Glu70γ-Ala/His85γ-Ala^ ‘triple-mutant’, were unable to stain BTN2A1^+^ cells, but still retained the ability to interact with mAb 20.1-pretreated BTN3A1^+^ cells (**Fig. 2E** and **F; Extended Data Fig. 4A**). Conversely, TCR tetramers with ‘epitope two’ mutations, G115^Arg51δ-Ala^ or G115^Glu52δ-Ala^, readily stained BTN2A1^+^ cells, but lost their ability to interact with mAb 20.1-pretreated BTN3A1^+^ cells (**Fig. 2E** and **F; Extended Data Fig. 4A**). G115^Lys108γ-Ala^, located within the CDR3γ and near the CDR2δ (5–8 Å away), also exhibited a reduced association with mAb 20.1-pretreated BTN3A1^+^ cells, but not to BTN2A1 (**Fig. 2E** and **F; Extended Data Fig. 4A**). Strikingly, G115^Lys53δ-Ala^ tetramers, which was the mutant that resulted in autoactivation in functional assays (**Extended Data Fig. 3A** and B), did not affect the interaction with BTN2A1^+^ cells, but stained BTN3A1^+^ cells even without any mAb 20.1 cross-linking (**Fig. 2E** and **F; Extended Data Fig. 4A**). Indeed, mAb 20.1 pre-treatment only marginally enhanced G115^Lys53δ-Ala^ tetramer staining of BTN3A1^+^ cells above this spontaneous level of interaction (**Fig. 2F** and **Extended Data Fig. 4A**). The strong interaction of G115^Lys53δ-Ala^ γδTCR tetramers with BTN3A1^+^ cells also held true for other Vγ9Vδ2^+^ TCR clones tested with the same substitution (**Extended Data Fig. 4B**), indicating that the Lys53δ-Ala mutation enhances Vγ9Vδ2^+^ TCR binding potential irrespective of CDR3 sequence heterogeneity. Tetramers with combined His85γ-Ala (in epitope one) and Glu52δ-Ala (in epitope two) mutations, G115^His85γ-Ala/Glu52δ-Ala^, lost the ability to interact with both BTN2A1^+^ and mAb 20.1-pretreated BTN3A1^+^ cells (**Fig. 2E** and **F; Extended Data Fig. 4A**). What is more, the antagonistic anti-BTN3A mAb (clone 103.2) completely prevented the binding of G115^Lys53δ-Ala^ γδTCR tetramers to BTN3A1^+^ cells implying, a specific interaction between these molecules, since it is unlikely that anti-BTN3A1 mAb could completely impair binding of Vγ9Vδ2 TCR to another unrelated ligand (**Extended Data Fig. 4C**). In further support of the observation that Vγ9Vδ2^+^ TCR closely associates with BTN3A1 following anti-BTN3A mAb 20.1-pretreatment, we co-stained BTN3A1- or BTN2A1-expressing cells with control SAv-PE or Vγ9Vδ2 TCR-PE tetramer along with isotype control-AF647 (MOPC21) or anti-BTN3A-AF647 (20.1) mΑb (**Extended Data Fig. 4D**). Förster resonance energy transfer (FRET) was observed when BTN3A1^+^ cells were co-stained with Vγ9Vδ2 TCR-PE tetramer and anti-BTN3A-AF647 Αb, suggesting close proximity (<10 nm) when co-bound to BTN3A1-transfected cells. Collectively, these data suggest ‘ligand-two’, being either BTN3A1 itself or a closely associated molecule, binds to Vγ9Vδ2^+^ TCR via ‘epitope two’, located on the apical surface of the Vγ9Vδ2^+^ TCR and incorporating residues within the CDR2δ and CDR3γ loops. Within epitope two, Lys53δ appears to act as a gatekeeper residue for ligand-two accessibility, suggesting that upon cross-linking of BTN3A1 with agonist mAb 20.1, a conformational change to ligand-two occurs that partly circumvents this steric barrier.

### BTN3A1 is a co-ligand of the Vγ9Vδ2^+^ TCR

We next explored the hypothesis that ligand-two is BTN3A1, and that BTN2A1 stabilizes BTN3A1 binding to the γδTCR. To test this, we produced soluble BTN3A1–BTN2A1 ectodomain heteromeric complexes (**Extended Data Fig. 5A–C**), which were tethered together with C-terminal leucine zippers, and measured whether they could bind to epitope two, being the ligand-two binding site on Vγ9Vδ2^+^ TCR. The BTN2A1–BTN3A1 heteromer complex retained staining with anti-BTN2A1 and anti-BTN3A1 mAb by ELISA (**Extended Data Fig. 5D**) and was comprised of two chains after purification (BTN2A1 and BTN3A1; **Extended Data Fig. 5B** and C) and following crystallisation (**Extended Data Fig. 5E**), suggestive of a correct conformation.

Consistent with the BTN2A1–γδTCR docking mode (**Fig. 1C**), BTN2A1 tetramers readily stained *BTN2A*^KO^.*BTN3A*^KO^ HEK293T cells expressing G115^WT^ TCR, as well as cells expressing G115 TCR mutants located in epitope two, namely G115^Glu52δ-Ala^ and G115^Lys53δ-Ala^, but not cells expressing the epitope one mutant G115^His85γ-Ala^ TCR (**Fig. 3A** and **Extended Data Fig. 5F**). Soluble BTN3A1 ectodomain tetramers failed to interact with G115 TCR^+^ HEK-293T cells [**Fig. 3A** and ^10^]. BTN2A1–BTN3A1 complex tetramers bound G115^WT^ TCR^+^ cells (**Fig. 3A** and **Extended Data Fig. 5F**). Akin to staining with BTN2A1 tetramers, a G115^His85γ-Ala^ mutation abrogated the interaction with BTN2A1–BTN3A1 tetramers, indicating a dependence on BTN2A1. However, unlike BTN2A1 tetramers, BTN2A1–BTN3A1 tetramer binding was modulated by mutations to epitope two. Here, G115^Glu52δ-Ala^, which was essential for TCR tetramer staining of BTN3A1-transfected cells (**Fig. 2F**), marginally reduced staining with BTN2A1–BTN3A1, compared to G115^WT^ TCR tetramer, whereas the gatekeeper residue mutant G115^Lys53δ-Ala^ resulted in a clear increase in BTN2A1–BTN3A1 staining intensity (**Fig. 3A** and **Extended Data Fig. 5F**). In agreement with the notion that epitope two on Vγ9Vδ2 TCR modulates the binding of the BTN2A1– BTN3A1 tetramer, a monovalent anti-BTN3A Fab fragment (clone 103.2) also interfered with BTN2A1–BTN3A1 tetramer interaction with Vγ9Vδ2 TCR, reinforcing the idea that ligand two is BTN3A1 (**Extended Data Fig. 5G**). These data indicate that soluble BTN2A1– BTN3A1 ectodomain complex can bind γδTCR, but, unlike BTN2A1 alone, BTN2A1– BTN3A1 complex binding is co-dependent on epitopes one and two.

**Figure 3.**
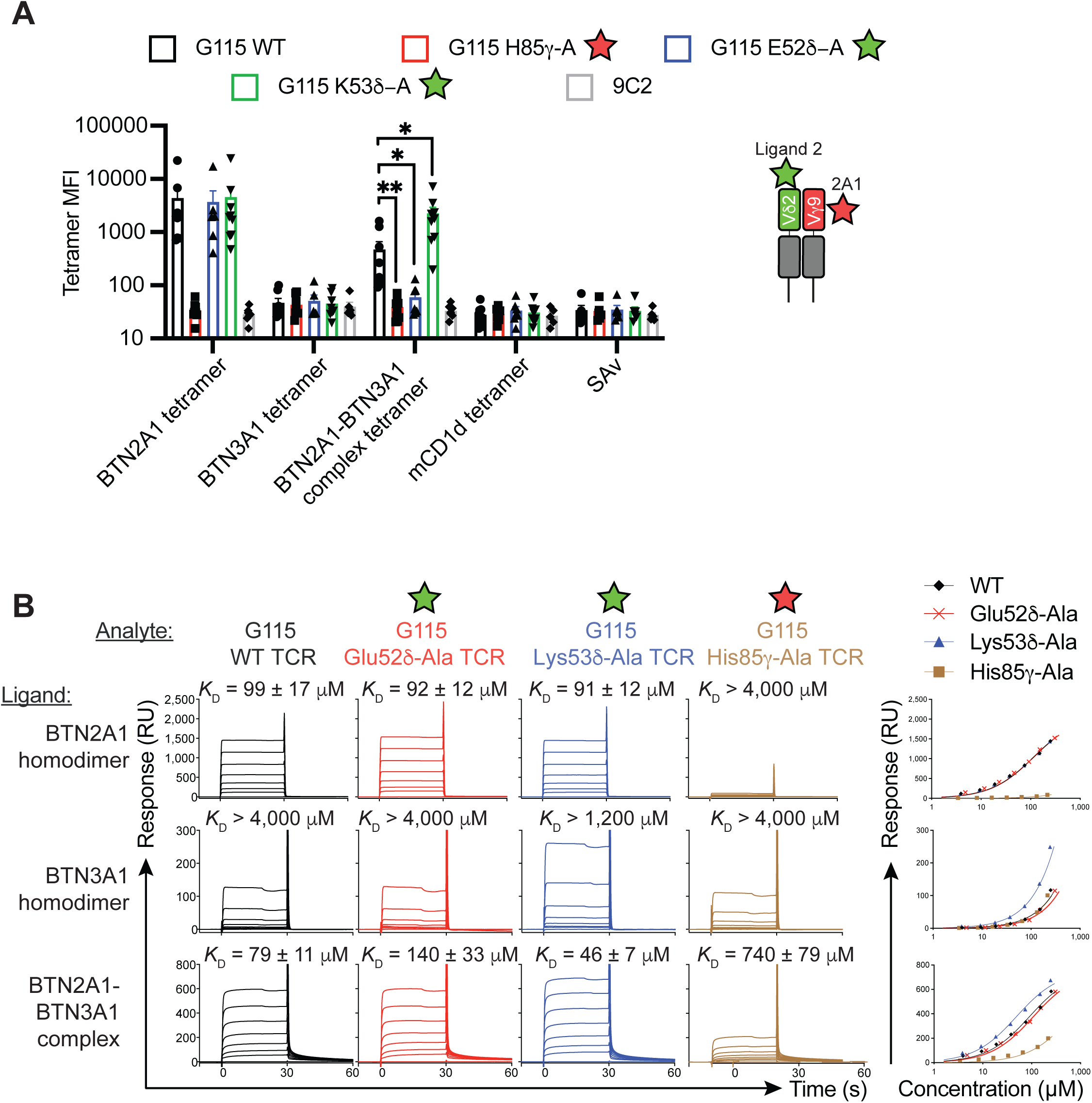
BTN3A1 is a ligand for the γδTCR. (**A**) BTN2A1-, BTN3A1-, BTN2A1– BTN3A1 complex- or control mouse CD1d-ectodomain tetramers, or control streptavidin-PE (SAv) staining of HEK293T cells co-transfected with CD3 plus G115 Vγ9Vδ2^+^ TCR wild-type, His85γ-Ala, Glu52δ-Ala, Lys53δ-Ala or control 9C2 Vγ5Vδ1^+^ TCR. Graphs depict mean ± SEM. N ≥ 5 independent experiments. * P < 0.05, ** P < 0.01; BTN2A1–BTN3A1 complex tetramer binding to Vγ9Vδ2^+^ TCRs tested by Kruskal-Wallis test with two-stage step-up multiple correction method of Benjamini, Krieger and Yekutieli. (**B**) Sensorgrams (left) and saturation plots (right) depicting binding of soluble G115 Vγ9Vδ2^+^ TCR wild-type (black, 302–4.2 µM), His85γ-Ala (brown, 410–1.6 µM), Glu52δ-Ala (red, 370–5.8 µM) and Lys53δ-Ala (blue, 295–4.6 µM) to immobilised BTN2A1 ectodomain homodimer, BTN3A1 ectodomain homodimer and BTN2A1–BTN3A1 ectodomain complex, as measured by surface plasmon resonance from the same experiment. *K*_D_, dissociation constant calculated at equilibrium ± SEM, derived from the mean of n=2 (WT and His85γ) or n=3 (Glu52δ and Lys53δ) independent experiments.

We next tested whether BTN2A1–BTN3A1 complexes can co-bind epitopes one and two of Vγ9Vδ2^+^ TCR in a cell-free assay, by using surface plasmon resonance (**Fig. 3B**). Soluble G115^WT^

γδTCR bound immobilized BTN2A1 homodimer with an affinity of *K*_D_ = 99 μM, which is similar to previous studies (*K*_D_ = 40–50 μM in ^11,12^), but did not bind immobilized BTN3A1 (*K*_D_ > 4,000 μM). Consistent with the role of epitope one, but not epitope two, in binding BTN2A1, soluble G115^His85γ-Ala^ TCR did not interact with BTN2A1, whereas G115^Glu52δ-Ala^ and G115^Lys53δ-Ala^ bound BTN2A1 equivalently to G115^WT^. Interestingly, TCR with the gatekeeper mutation G115^Lys53δ-Ala^ may have exhibited some low-level binding to BTN3A1 at the highest concentrations, but the predicted affinity was very weak (*K*_D_ > 1,200 μM). G115^WT^ γδTCR bound immobilized BTN2A1–BTN3A1 complex with a similar affinity to BTN2A1 (*K*_D_ = 79 μM and 99 μM, respectively). However, in contrast to G115 TCR binding to BTN2A1, binding of G115 TCR to BTN2A1–BTN3A1 was modulated by mutations within epitope two of the γδTCR, since G115^Lys53δ-Ala^ resulted in an increase in affinity (*K*_D_ = 46 μM) whereas G115^Glu52δ-Ala^ resulted in a slight decrease in affinity (*K*_D_ = 140 μM). Furthermore, unlike BTN2A1, the BTN2A1–BTN3A1 complex also appeared to interact weakly with G115^His85γ-Ala^ TCR (*K*_D_ = 740 μM; **Fig. 3B**). Therefore, the pattern of ligand-two interaction with Vγ9Vδ2^+^ TCR can be recapitulated with soluble BTN2A1–BTN3A1 complex, in both cell-surface staining-based and cell-free biophysical-based assays. Together, these data reveal that BTN3A1 directly interacts with epitope two on the δ-chain of the Vγ9Vδ2^+^ TCR, and along with BTN2A1, is necessary and sufficient to co-engage γδTCR.

### BTN3A1 IgV domain interacts with both BTN2A1 and Vγ9Vδ2^+^ TCR

BTN2A1 and BTN3A1 are located within 10 nm of each other in cis on the cell surface ^11^, however, whether they directly interact is unclear. Using surface plasmon resonance, full-length BTN3A1 ectodomain (IgV–IgC) bound immobilized disulfide-linked BTN2A1 homodimer with an affinity of *K*_D_ = 500 μM, but not immobilized BTN3A1 homodimer. Conversely, full-length BTN2A1 ectodomain weakly bound immobilised BTN3A1 homodimer (*K*_D_ ∼1,800 μM), but not immobilized BTN2A1 homodimer (**Fig. 4A**), indicating that BTN2A1 and BTN3A1 ectodomains are capable of directly interacting, albeit with a low affinity. Since BTN3A1 ectodomain exists as a homodimer and may therefore exhibit enhanced binding in SPR assays due to increased avidity, we also tested monomeric BTN3A1 IgV domain, which retained specific binding to BTN2A1 (*K*_D_ = 1,100 µM; **Fig. 4A**). To understand the molecular nature of this interaction, we crystallized the BTN2A1– BTN3A1–zipper complex ectodomains. The crystals diffracted weakly (anisotropic diffraction range 5.6 Å – 8.9 Å resolution), which nonetheless yielded a single solution in space group *F*222, wherein the asymmetric unit contained a single copy of BTN2A1 and BTN3A1, that interfaced via their IgV domains at a docking angle of ∼29° (**Fig. 4B** and **Extended Data Table 1**). The C-terminal zipper domains were not modelled, although there was space for them within the asymmetric unit underneath the IgC domains. An unbiased electron density map enabled unambiguous identification of the IgV and IgC domains (**Extended Data Fig. 6A**). V-shaped homodimers of both BTN2A1 and BTN3A1 were also present within the unit cell (**Fig. 4B**), which were structurally similar to those found in the apo crystal structures, although they were twisted slightly (∼10° and ∼16°, respectively; **Extended Data Fig. 6B**). The BTN2A1 and BTN3A1 V-dimers buried 620 Å^2^ and 640 Å^2^, respectively, for a combined BSA of ∼1,300 Å^2^ (**Extended Data Fig. 6C**). The BTN2A1 and BTN3A1 V-dimers came together at a planar angle of ∼80° to form a distorted W-shaped heterotetramer (**Fig. 4B**), which could be even further expanded through crystallographic symmetry to yield a linear polymer of the composition [BTN2A1_homodimer_–BTN3A1_homodimer_]_n_ (**Fig. 4C**).

**Figure 4.**
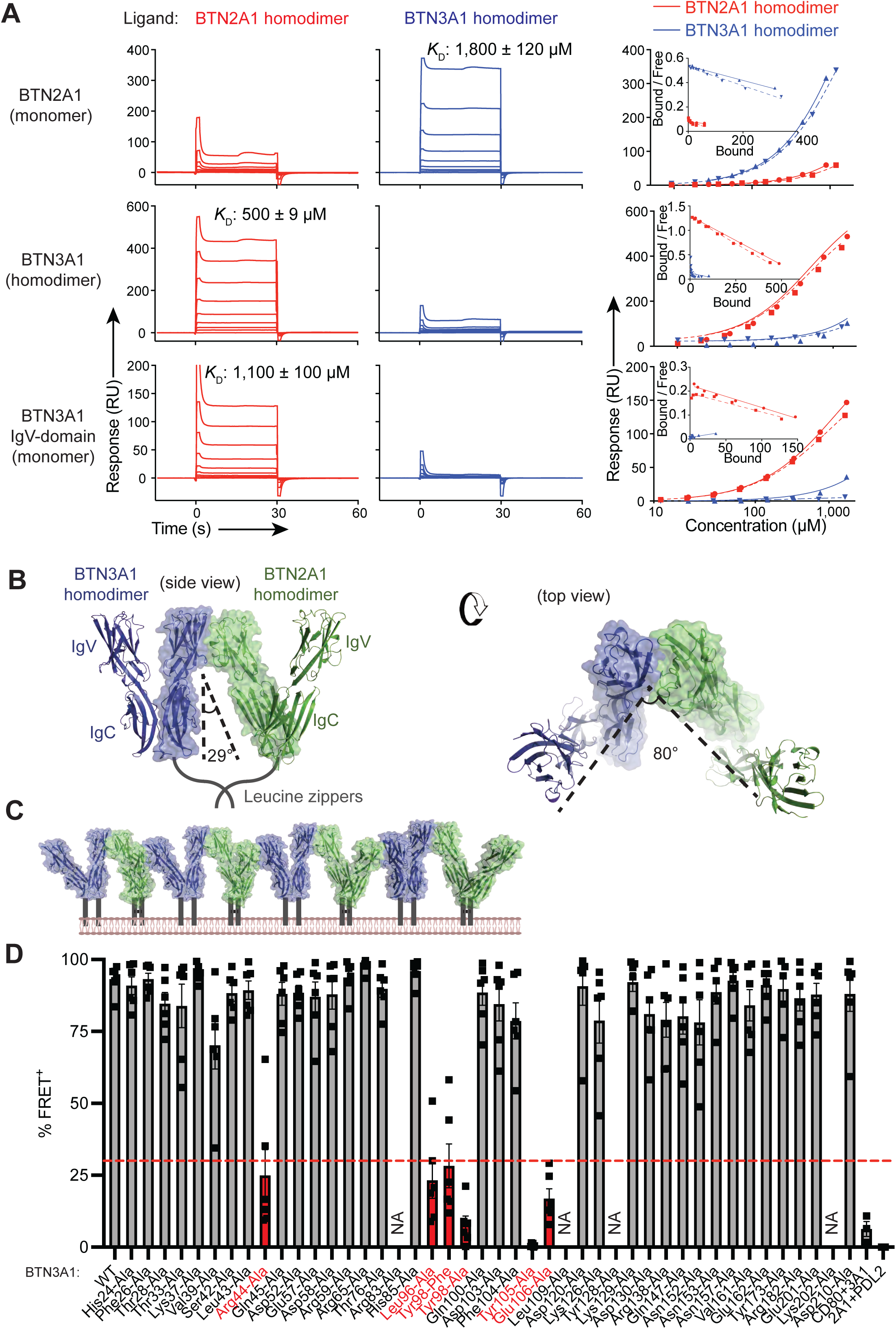
BTN2A1 and BTN3A1 directly associate and form heteromers. (**A**) Sensorgrams (left) and saturation plots (right) depicting binding of soluble monomeric BTN2A1 ectodomain (top row, 890–28 µM), homodimeric BTN3A1 ectodomain (middle row, 1,520–24 µM), or monomeric BTN3A1 IgV domain (bottom row, 1,590–25 µM) to immobilised BTN2A1 ectodomain homodimer (red) or BTN3A1 ectodomain homodimer (blue), as measured by surface plasmon resonance. Insert graphs depict Scatchard plots. *K*_D_, dissociation constant calculated at equilibrium ± SEM, derived from the mean of n=2 independent experiments each shown separately as dotted and close lines on the saturation plots. (**B**) The BTN2A1–BTN3A1 ectodomain complex crystal structure, showing the asymmetric unit as a surface and the V-dimers as a cartoon. BTN3A1, blue; BTN2A1, green. (**C**) Surface and cartoon representation of the BTN2A1 V-dimer–BTN3A1 V-dimer repeating unit within the crystal structure. Color scheme as in (B). (**D**) Association between BTN2A1 and BTN3A1 ectodomains on the cell surface of mouse NIH-3T3 fibroblasts co-expressing wild-type BTN2A1 and individual BTN3A1 mutants, as determined by mean percentage ± SEM of FRET^+^ cells between anti-BTN2A1-AF647 (clone 259) and anti-BTN3A-PE (clone 103.2). Controls (right) depict FRET between transiently expressed CD80 and BTN3A1, or BTN2A1 and PD-L2. n = 6 where each point represents an individual experiment, except for controls where n = 3. NA, data were excluded if BTN3A1 mutant protein levels were > 2-fold lower than BTN3A1 WT as determined by anti-BTN3A (clone 103.2) mAb staining.

The BTN2A1–BTN3A1 intermolecular contacts were determined based on a model where the higher resolution apo BTN2A1 and BTN3A1 structures were overlaid on to the low-resolution BTN2A1–BTN3A1 complex (**Extended Data Fig. 6D–G** and **Extended Data Table 5**). Using this model, and assuming no significant changes in side-chain positions, we have identified a number of contacts between BTN2A1 and BTN3A1. These include BTN2A1^Phe43^ which interacts with BTN3A1^Arg44^ and a mainchain interaction with BTN3A1^Ser41^, and BTN2A1^Ser44^ that also interacts with BTN3A1^Ser41^. We have also identified that BTN2A1^Glu107^ interacts with BTN3A1^Arg44^. BTN2A1 residues ^Glu35^, ^Lys51^ and ^Gln100^ all appear to interact with BTN3A1^Tyr105^. Of these residues that are identified as interacting with BTN3A1 based on the crystal structure, BTN2A1^Phe43^, ^Ser44^, ^Gln100^ and ^Glu107^ were also implicated in the interaction with Vγ9. BTN2A1^Glu107^ was also previously shown by Thomas Herrmann and colleagues to be important for interacting with Vγ9 ^12^.

We next tested which BTN3A1 residues are responsible for engaging BTN2A1 in cis on the cell surface ^11^. Of a panel of forty-five BTN3A1 Ala ectodomain mutants, including residues within both the IgV and IgC domains, forty retained expression on the cell surface and retained staining by anti-BTN3A mAb clone 103.2 (**Extended Data Fig. 7A** and B). Mutations to five residues: BTN3A1^Arg44-Ala^, BTN3A1^Leu96-Ala^, BTN3A1^Tyr98-Ala^ (and additionally BTN3A1^Tyr98-Phe^), BTN3A1^Tyr105-Ala^ and BTN3A1^Glu106-Ala^ abrogated FRET between anti-BTN2A and anti-BTN3A mAbs (**Fig. 4D** and **Extended Data Fig. 7C**). Thus, mutations to these residues disrupt the association between BTN2A1 and BTN3A1 on the cell surface. These residues mapped to the CFG face of BTN3A1 and correlated closely with the crystal structure interface (**Extended Data Fig. 7D**), thereby affirming this mode of binding. Accordingly, BTN2A1 and BTN3A1 interact via the CFG faces of their IgV domains and form W-shaped heterodimers and/or hetero-oligomers.

Using this same panel of BTN3A1 Ala mutants, we investigated which residues were involved in the Vγ9Vδ2^+^ TCR interaction. Thirty-four of the panel of forty-five mutants retained binding to anti-BTN3A mAb clone 20.1 **(Extended Data** Figs. 7A–B and **8A)**. Of these, six completely abrogated G115 tetramer staining of mAb 20.1-pretreated BTN3A1^+^ cells: BTN3A1^Val39^, BTN3A1^Arg44^, BTN3A1^His85^, BTN3A1^Tyr98^, BTN3A1^Phe104^ and BTN3A1^Tyr105^, plus a further four residues that reduced G115 tetramer staining by >90%: Phe26, Lys37, Ser42 and Leu96 (**Fig. 5A** and **Extended Data Fig. 8A**). The panel of BTN3A1 Ala mutants were next co-expressed with BTN2A1 (WT) in NIH-3T3 cells and used to activate Vδ2^+^ T cells in the presence of zoledronate. All six BTN3A1 Ala mutants that abrogated G115 tetramer interaction – BTN3A1^Val39^, BTN3A1^Arg44^, BTN3A1^His85^, BTN3A1^Tyr98^, BTN3A1^Phe104^ and BTN3A1^Tyr105^ – also abrogated Vδ2^+^ T cell activation, as did BTN3A1^Leu96^ (**Fig. 5B** and **Extended Data Fig. 8B**). Except for BTN3A1^His85^, which mapped to the ABED face, all other residues mapped to the CFG face. These data extend upon an earlier report that the CFG face of BTN3A1 IgV domain is functionally important ^13^, and moreover, attribute a role for these residues in binding to Vγ9Vδ2^+^ TCR.

**Figure 5.**
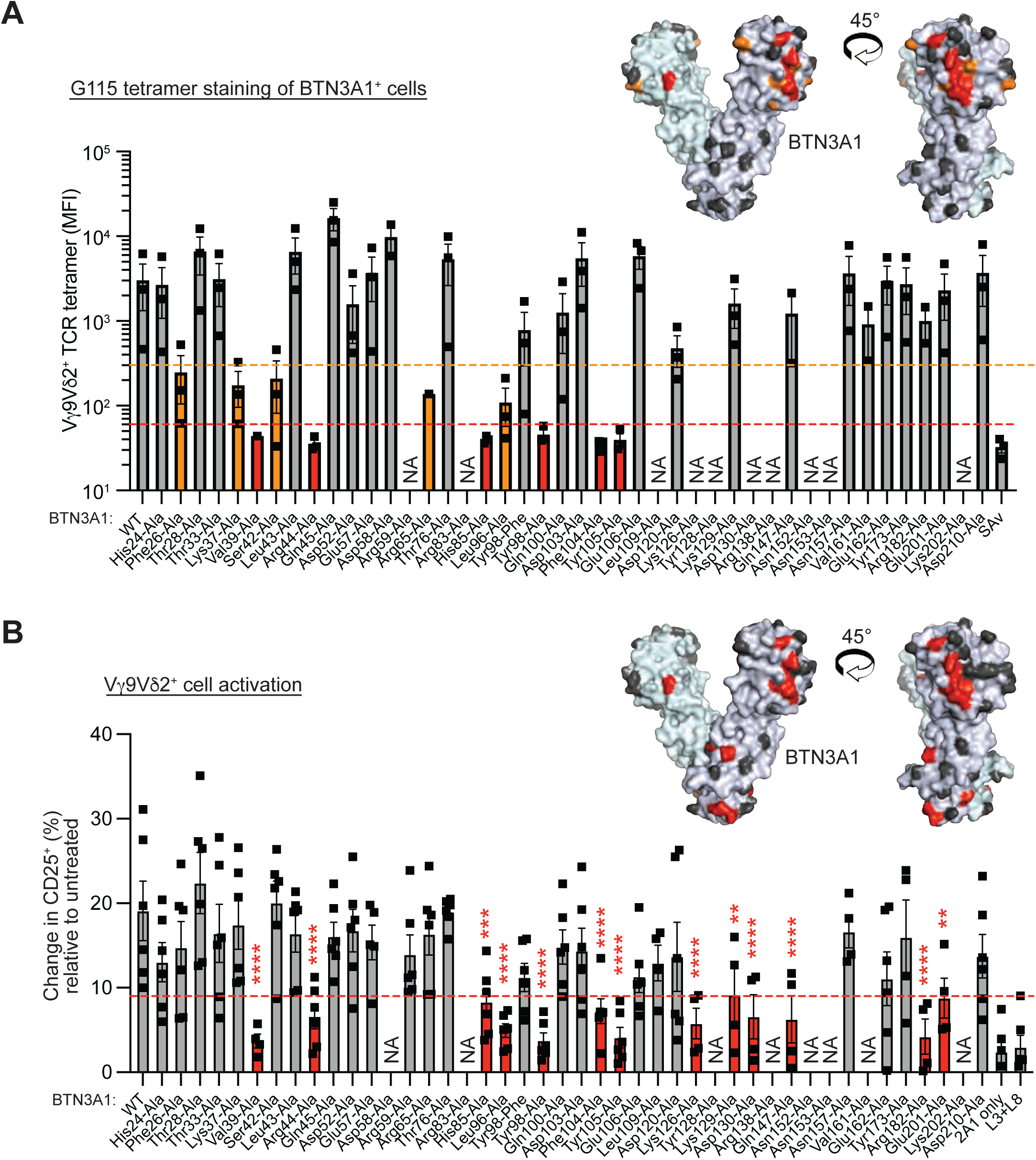
BTN3A1 IgV domain interacts with Vγ9Vδ2^+^ TCR. (**A**) G115 Vγ9Vδ2^+^ TCR tetramer-PE staining of mouse NIH-3T3 fibroblasts transfected with either wild-type BTN3A1 or the indicated mutants, following pre-treatment with anti-BTN3A1-AF647 (clone 20.1) antibody. SAv, streptavidin-PE control staining of wild-type BTN3A1^+^ cells. Bar graphs depict median fluorescence intensity (MFI) ± SEM. Dotted lines represents 90–98% reduction (orange) and >98% reduction (red) in MFI. Inset: surface representation of BTN3A1 V-dimer with mutations to residues that led to an abrogation of the anti-BTN3A antibody (20.1)-dependent G115 tetramer interaction coloured in red (> 98% reduction), orange (90–98% reduction), or gray (< 90% reduction). n = 3, where each point represents an independent experiment. NA, not applicable since BTN3A surface expression was too low to measure G115 tetramer staining. (**B**) Change in CD25 expression (normalized to unstimulated control for each sample) on purified in vitro-expanded Vδ2^+^ γδ T cells co-cultured for 24 h with 5 µM zoledronate and mouse NIH-3T3 fibroblast APCs transfected with wild-type *BTN2A1* and individual *BTN3A1* mutants. Bar graphs depict mean ± SEM. Red dotted line represents >50% reduction in activation compared to BTN2A1–BTN3A1 WT. NA = data not available since BTN3A1 levels were too low to induce zoledronate-dependent activation of γδ T cells. Data are from 2-3 independent experiments each with n=1-2 different donors each. ** P < 0.01, *** P < 0.001, **** P < 0.0001, by two-way ANOVA with Šidák multiple comparison correction. Inset: surface representation of BTN3A1 V-dimer with mutations to residues that led to an abrogation of zoledronate-dependent Vδ2^+^ γδ T cell activation shown in red, or did not impact Vδ2^+^ γδ T cell activation shown in gray.

We also note that mutation of several residues in BTN3A1 appeared to have a subtle increase in binding of Vγ9Vδ2 TCR tetramers, especially Gln45-Ala (**Fig. 5A** and **Extended Data Fig. 8A**). Soluble BTN3A1 ectodomain protein with the Gln45-Ala mutation was next produced to test whether soluble BTN3A1 could be engineered to improve its interaction with Vγ9Vδ2 TCR, and measure direct binding to Vγ9Vδ2 TCR in the absence of BTN2A1. Indeed, Gln45-Ala BTN3A1 tetramers specifically bound to cells expressing the Lys53-Ala variant of the G115 Vγ9Vδ2 TCR (**Extended Data Fig. 8C**), albeit weakly. Since this interaction is Lys53-Ala-augmented, it indicates that binding is occurring via epitope two on the Vγ9Vδ2 TCR, which is consistent with the binding mode we have already demonstrated (**Fig. 2F**). Thus, despite the challenges in working with soluble BTN3A1, binding can indeed be observed between soluble BTN3A1 and Vγ9Vδ2 TCR in the absence of BTN2A1, when the constructs are optimized to enhance the binding affinity.

### BTN2A1 and BTN3A1 utilize the same epitopes to bind each other and Vγ9Vδ2^+^ TCR

Paradoxically, four of the seven BTN3A1 Ala mutants (BTN3A1^Arg44^, BTN3A1^Leu96^, BTN3A1^Tyr98^ and BTN3A1^Tyr105^) that were important for binding to Vγ9Vδ2^+^ TCR were also critical for binding to BTN2A1 (**Figs. 4D, 5A,** and **Extended Data Fig. 7D**). Likewise, many of the residues within the BTN2A1 IgV domain that contacted γδTCR also mediated binding to BTN3A1, including BTN2A1^Phe43^ and BTN2A1^Ser44^ (**Extended Data Tables 4** and **5; Extended Data Fig. 8D**). Since the regions of BTN3A1 that engage BTN2A1 and γδTCR are overlapping, and conversely, the regions of BTN2A1 that engage BTN3A1 and γδTCR are also overlapping, it was challenging to reconcile how BTN2A1 and BTN3A1 can co-bind to each other along with Vγ9Vδ2^+^ TCR. Indeed, a superimposition of the BTN2A1–γδTCR and BTN2A1–BTN3A1 crystal structures identified major steric clashes between BTN3A1 and the γδTCR, suggesting that co-binding as a ternary complex in this manner is unlikely (**Extended Data Fig. 8E** and **Extended Data Movie 1**). This implies that whilst both BTN2A1 and BTN3A1 are ligands for each other, they must disengage or undergo major conformational changes prior to co-binding Vγ9Vδ2^+^ TCR.

We tested this hypothesis using BTN2A1–BTN3A1–zipper ectodomain complex tetramers that contained the BTN3A1^Glu106-Ala^ mutant, which, based on FRET measurements, largely disrupts the natural IgV–IgV domain interaction between BTN2A1 and BTN3A1, without having any effect on binding to Vγ9Vδ2^+^ TCR (**Figs. 4D** and **5A**, respectively). Compared to BTN2A1^WT^–BTN3A1^WT^ tetramers, BTN2A1^WT^–BTN3A1^Glu106-Ala^ tetramers stained G115^WT^ γδTCR-transfected HEK-293T cells at a higher intensity, suggesting that the affinity may be increased (**Fig. 6A**). G115^Glu52δ-Ala^ γδTCR, which abrogates binding to BTN3A1, was also stained more brightly by the BTN2A1^WT^–BTN3A1^Glu106-Ala^ tetramers, further suggesting that despite disrupting the capacity of BTN3A1 to engage γδTCR, binding of BTN2A1 within the BTN2A1^WT^–BTN3A1^Glu106-Ala^ complex to γδTCR is actually enhanced. The BTN2A1^WT^–BTN3A1^Glu106-Ala^ tetramers also stained G115^WT^-expressing cells more brightly than BTN2A1-only tetramers, implying that BTN2A1^WT^–BTN3A1^Glu106-Ala^ has a higher affinity for Vγ9Vδ2^+^ TCR compared to apo BTN2A1. These observations supported the hypothesis that disrupting the IgV–IgV interaction within a BTN2A1–BTN3A1 complex enhances overall binding to Vγ9Vδ2^+^ TCR, but to directly measure the binding affinity, we performed SPR (**Fig. 6B**). Consistent with the tetramer staining of cell lines, G115^WT^ γδTCR bound immobilized BTN2A1^WT^–BTN3A1^Glu106-Ala^ complexes with a higher affinity than BTN2A1^WT^–BTN3A1^WT^ complexes (*K*_D_ = 37 μM compared to 85 μM). The affinity of G115^WT^ γδTCR for BTN2A1^WT^–BTN3A1^Glu106-Ala^ complexes was also higher than it was for BTN2A1-only (*K*_D_ = 37 μM compared to 99 μM; **Fig. 6B**), supporting a model wherein disruption of the natural IgV–IgV-mediated interaction between BTN2A1 and BTN3A1 facilitates enhanced co-binding of BTN2A1 and BTN3A1 to Vγ9Vδ2^+^ TCR.

**Figure 6.**
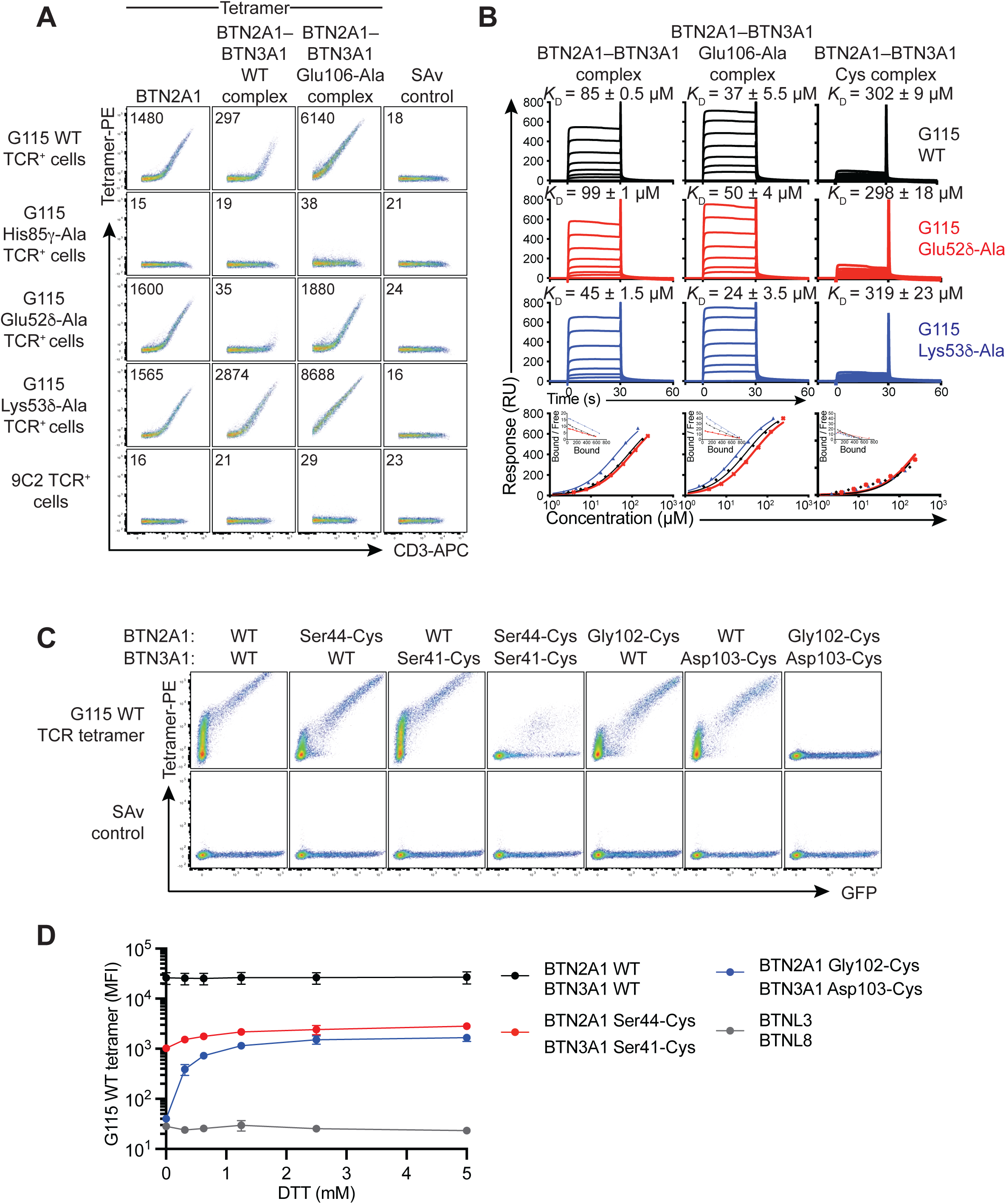
BTN2A1 and BTN3A1 must disengage in order to bind Vγ9Vδ2^+^ TCR. (**A**) BTN2A1 tetramer, BTN2A1–BTN3A1 WT complex tetramer, BTN2A1–BTN3A1 Glu135-Ala complex tetramer, or control streptavidin alone (SAv), versus anti-CD3 staining on HEK293T cells co-transfected with CD3 plus G115 Vγ9Vδ2^+^ TCR wild-type, Glu52δ-Ala, or control 9C2 Vγ5Vδ1^+^ TCR. Inset – median fluorescence intensity (MFI) of PE parameter. Representative of one of six independent experiments. (**B**) Sensorgrams (top) and saturation plots (bottom) depicting binding of soluble G115 Vγ9Vδ2^+^ TCR wild-type (black, 181–2.8 µM), Glu52δ-Ala (red, 243–3.8 µM) and Lys53δ-Ala (blue, 139–2.2 µM) to immobilised BTN2A1–BTN3A1 wild-type (left) or BTN2A1–BTN3A1 Glu135-Ala (right) complexes. *K*_D_, dissociation constant calculated at equilibrium ± SEM, derived from the mean of two independent experiments. (**C**) G115 Vγ9Vδ2^+^ TCR tetramer-PE, or control streptavidin-PE (SAv) staining of mouse NIH-3T3 fibroblasts co-transfected with BTN2A1 and BTN3A1 wild-type or cysteine mutants in the depicted combinations. Representative of one of two independent experiments. (**D**) G115 tetramer-PE staining of NIH-3T3 fibroblasts co-transfected with either WT or Cys-mutant BTN2A1 plus BTN3A1, or control BTNL3 plus BTNL8, following pre-incubation of the cells with DTT at indicated concentrations. Graphs depict mean ± SEM. Data pooled from 3-4 separate experiments.

The collective data thus far suggests that BTN2A1 and BTN3A1 epitopes are tethered to each other, which impairs the Vγ9Vδ2^+^ TCR from efficiently engaging. To further test this hypothesis, we reasoned that locking BTN2A1 and BTN3A1 IgV domains together would abrogate their interaction with Vγ9Vδ2^+^ TCR. For this, we introduced cysteine (Cys) residues in the BTN2A1 and BTN3A1 IgV domain CFG faces that, based on the BTN2A1–BTN3A1 crystal structure, were optimally positioned for formation of an interchain disulfide bond. We identified two separate Cys pairs: BTN2A1^Gly102-Cys^ plus BTN3A1^Asp103-Cys^, and BTN2A1^Ser44-Cys^ plus BTN3A1^Ser41-Cys^ (**Extended Data Fig. 9A**). Cells co-transfected with BTN2A1^Ser44-Cys^ plus BTN3A1^Ser41-Cys^, or BTN2A1^Gly102-Cys^ plus BTN3A1^Asp103-Cys^, exhibited a major reduction or total loss of G115 tetramer binding, respectively (**Fig. 6C**). However, when the Cys mutants were co-expressed with the corresponding WT molecule (for example, BTN2A1^Gly102-Cys^ plus BTN3A1^WT^, or vice versa), their binding to Vγ9Vδ2^+^ TCR was retained (**Fig. 6C**). Treatment of BTN2A1^Cys+^ BTN3A1^Cys+^ cells with graded doses of the reducing agent dithiothreitol (DTT) partly restored the ability of G115 tetramer to stain these cells, indicating that an interchain disulfide bond was responsible for the loss of interaction with Vγ9Vδ2^+^ TCR (**Fig. 6D** and **Extended Data Fig. 9B**).

Based on the BTN2A1–BTN3A1 crystal structure, we predicted that soluble BTN2A1^Gly102-Cys^–BTN3A1^Asp103-Cys^ ectodomain complexes would adopt an M-shaped tetramer comprised of a core BTN3A1 V-dimer and two outer copies of BTN2A1, each linked to BTN3A1 via a disulfide bond (**Extended Data Fig. 9C**). In support of this, 2D class averages of negatively stained electron micrographs of soluble BTN2A1^Gly102-Cys^– BTN3A1^Asp103-Cys^ complex revealed the presence of M-shaped particles (**Extended Data Fig. 9D**). A tetrameric form of the BTN2A1^Gly102-Cys^–BTN3A1^Asp103-Cys^ heteromers was completely unable to stain G115^WT^ Vγ9Vδ2^+^ TCR-transfected HEK-293T cells; however, pre-treatment of these soluble tetrameric complexes with DTT immediately prior to staining in order to break the interchain disulfide bonds, restored their ability to interact with G115 γδTCR^+^ cells (**Extended Data Fig. 9E** and F). Thus, locking BTN2A1 and BTN3A1 together with a covalent bond prevents them from interacting with Vγ9Vδ2^+^ TCR, and disruption of this bond restores the Vγ9Vδ2^+^ TCR interaction, suggesting that the epitopes that tether BTN2A1 and BTN3A1 to one-another are also important for mediating the interactions with Vγ9Vδ2^+^ TCR.

### Sequestration of pAg by BTN3A1 enhances Vγ9Vδ2^+^ TCR binding

The intracellular domains of BTN2A1 and BTN3A1 are both required for pAg-induced activation of Vγ9Vδ2^+^ T cells ^11,19^, and it has recently been found that pAg induces an association between the BTN3A1 and BTN2A1 intracellular domains ^20,21^. In further support of these later studies, we identified three residues within the BTN2A1 intracellular domain – two in the C-terminal cytoplasmic tail (BTN2A1^Thr482^ and BTN2A1^Leu488^) and one in the B30.2 domain (BTN2A1^Arg449^) – that are critical for the activation of Vγ9Vδ2^+^ T cells (**Extended Data Fig. 10A-C**). Using biolayer interferometry, we found that the B30.2 domain of BTN3A1 and the BTN2A1 full-length intracellular domain indeed interacted following pAg treatment (*K*_D_ ∼200 – 800 nM) and that mutations of BTN2A1^Arg449-Ala^ and BTN2A1^Thr482-Ala^ abrogate this pAg-induced association (**Extended Data Fig. 10D**). Thus, we propose that pAg sequestration of pAg by the intracellular domains facilitates allosteric changes to the ectodomains of BTN2A1–BTN3A1, converting them from an inactive state into an active state.

### Zoledronate challenge enhances binding of BTN3A1 to Vγ9Vδ2^+^ TCR

Our findings reveal that BTN3A1 can bind Vγ9Vδ2^+^ TCR following treatment with agonist anti-BTN3A mAb clone 20.1. Whilst this mAb is considered a mimic of natural pAg-driven activation, we sought to determine whether pAg themselves could also enhance the BTN3A1– Vγ9Vδ2^+^ TCR interaction. This was technically challenging, since pAg activity requires co-expression of BTN2A1 and BTN3A1 on APCs, yet Vγ9Vδ2^+^ TCR tetramers bind strongly to BTN2A1, irrespective of whether BTN3A1 is co-expressed. To overcome this challenge, we used G115^His85γ-Ala^ mutant Vγ9Vδ2^+^ TCR tetramers, which greatly reduces the TCR interaction with BTN2A1 [^11^ and **Fig. 2C**]. Despite the inability of G115^His85γ-Ala^ mutant tetramer to bind BTN2A1-only or BTN3A1-only transfected cells, co-expression of BTN2A1 and BTN3A1 supported binding of G115^His85γ-Ala^ tetramers (**Extended Data Fig. 11A**). This appeared to be dependent on BTN3A1, since treatment with the anti-BTN3A antagonist mAb 103.2 prevented G115^His85γ-Ala^ tetramer binding to BTN2A1 and BTN3A1 co-expressing cells (**Extended Data Fig. 11B**). Treatment with zoledronate significantly enhanced binding of the G115^His85γ-Ala^ tetramer to BTN2A1 plus BTN3A1 co-expressing cells in a dose-dependent manner (**Figs. 7A** and **Extended Data Fig. 11A**). This effect was also abrogated by pre-treatment with the HMG-CoA inhibitor, simvastatin (**Extended Data Fig. 11C**). Simvastatin also blocked zoledronate-dependent activation of Vγ9Vδ2^+^ T cells, supporting its ability to prevent zoledronate-induced accumulation of endogenous pAg (**Extended Data Fig. 11D**). Since zoledronate might also have off-target effects on cells due to generalized disruption of isoprenoid synthesis, we next tested whether the zoledronate-mediated enhancement of G115^His85γ-Ala^ tetramer binding was acting specifically via pAg binding to intracellular BTN3A1 and BTN2A1 B30.2 domains. We generated an extensive panel of modified BTN2A1 and BTN3A1 constructs that contained putative loss-of-function alterations to their intracellular domains, including truncated variants, domain-swapped variants, and variants with Ala substitutions at residues directly involved in pAg sequestration (**Fig. 7B, schematic**). As expected, zoledronate treatment induced a ∼6-fold increase in G115^His85γ-Ala^ tetramer mean fluorescence intensity of BTN2A1^WT^ plus BTN3A1^WT^ transfected cells, predominately by increasing the affinity of the TCR tetramers for cells with lower BTN expression, compared to untreated cells (**Fig. 7B**). Notably however, with the possible exception of BTN2A1^Leu488-Ala^, all other BTN2A1 and BTN3A1 variants with altered intracellular domains failed to facilitate a zoledronate-dependent enhancement in G115^His85γ-Ala^ tetramer staining (**Figs. 7B** and **Extended Data Fig. 11E**). This strongly indicates that the zoledronate-mediated enhancement of G115^His85γ-Ala^ tetramer staining is due to accumulation of endogenous pAg following zoledronate treatment.

**Figure 7.**
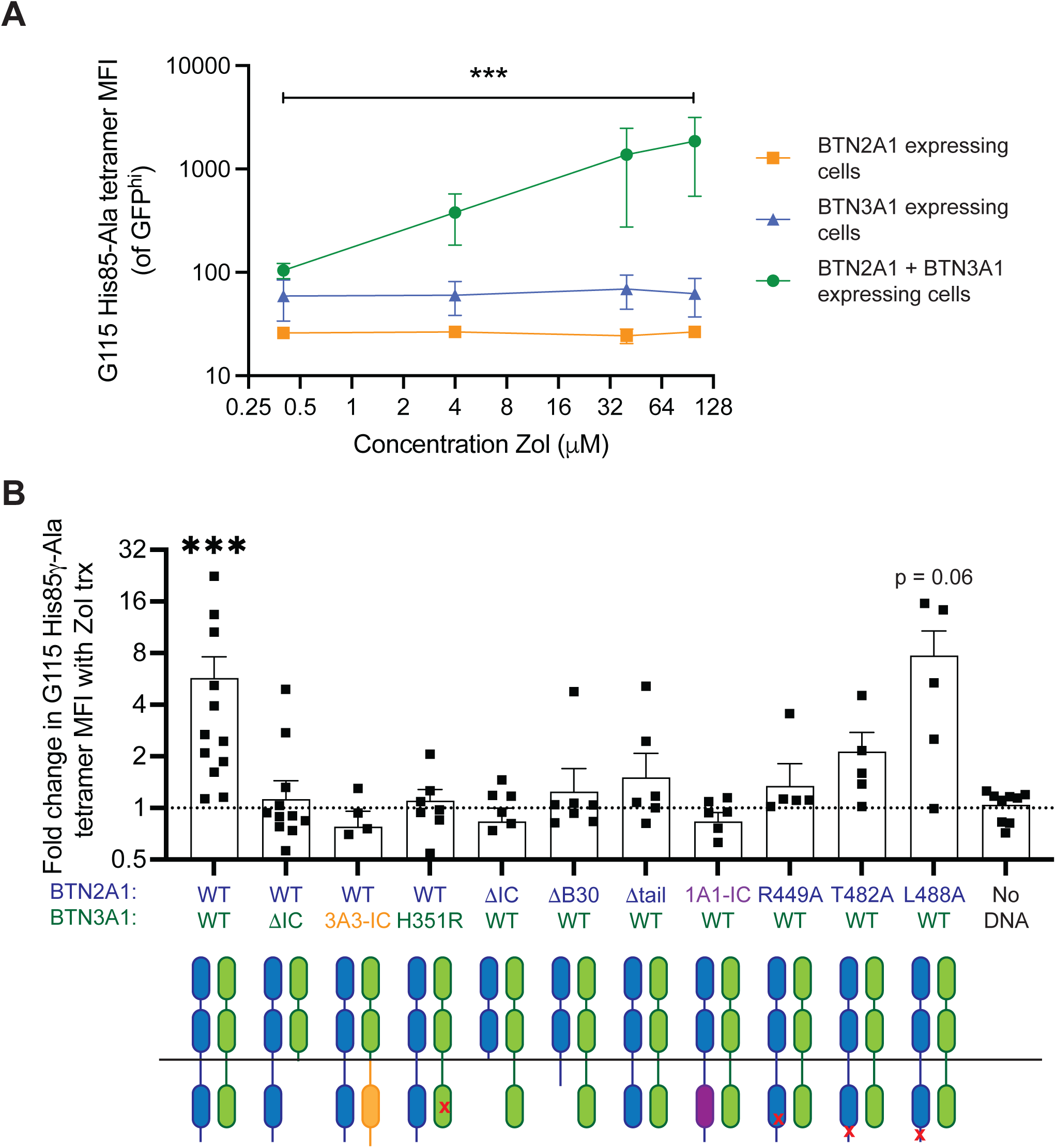
Zoledronate augments the BTN3A1-dependent interaction with Vγ9Vδ2 TCR. (**A**) Median fluorescence intensity (MFI) of Vγ9Vδ2^+^ ‘G115 His85γ-Ala’ TCR tetramer staining GFP^hi^ mouse NIH-3T3 cells transfected with BTN2A1, BTN3A1 or both BTN2A1 and BTN3A1, pre-treated with various concentrations of Zol for 16 hr. Graphs depict mean ± SEM. n = 4 (BTN3A1 expressing cells and BTN2A1 plus BTN3A1 co-expressing cells) or 2 (BTN2A1 expressing cells) separate experiments. *** P < 0.001; effect of Zol on BTN2A1 and BTN3A1 expressing cells tested by Kruskal-Wallis test with two-stage step-up multiple correction method of Benjamini, Krieger and Yekutieli. (**B**) NIH-3T3 cells expressing intracellular mutants of BTN2A1 (blue schematics) and BTN3A1 (green schematics) were treated with 100 µM zoledronate (Zol) for 16 h, stained with the Vγ9Vδ2 ‘G115 His85γ-Ala’ TCR tetramer and examined by flow cytometry. Graph depicts change in His85γ-Ala tetramer MFI with zoledronate treatment compared to untreated. *** P < 0.001 by Wilcoxon matched-pairs signed rank test; n ≥ 6. ΔIC, entire intracellular (IC) domain deleted; ΔB30, B30.2 and tail IC domains removed; Δtail, IC tail domain after B30.2 domain deleted; 3A3-IC, BTN3A1 IC domain replaced with BTN3A3 IC domain; 1A1-IC, BTN2A1 B30.2 domain replaced with B30.2 domain of BNT1A1. Schematics representative of mutations depicted below graph.

Collectively, these data indicate that the intracellular domains of BTN2A1 and BTN3A1 facilitate a conformational change to their ectodomains, which in turn enhances binding to Vγ9Vδ2^+^ TCR. Vγ9Vδ2^+^ TCR co-binds BTN2A1 and BTN3A1 via two spatially distinct epitopes, with BTN2A1 engaging the side of Vγ9, and BTN3A1 binding to the apical surface. The activated BTN2A1–BTN3A1 complexes can interact with Vγ9Vδ2^+^ TCR, facilitating γδ T cell-mediated immunity (**Extended Data** Fig. 12).

## Discussion

Akin to pMHC recognition by the αβTCR, BTN molecules have emerged as critical γδTCR ligands, however, their molecular mode of recognition is poorly defined. Furthermore, the precise mechanism by which Vγ9Vδ2^+^ T cells recognise pAg remains unclear. Here we report the first structure of a TCR engaging an endogenous non-MHC or MHC-like ligand, namely BTN2A1, revealing that BTN2A1 engages the side of Vγ9, leaving the apical face of the Vγ9Vδ2^+^ TCR exposed. We also demonstrate that a second ligand, BTN3A1, binds the apical Vγ9Vδ2^+^ TCR surface with low affinity, resolving a decade long mystery of the mechanism behind pAg detection by Vγ9Vδ2^+^ T cells that is dependent on BTN3A1.

The ability of Vγ9Vδ2^+^ TCR to co-bind two ligands contrasts the recognition of MHC and MHC-like molecules by αβ T cells, which bind with one-to-one stoichiometry. Thus, the γδTCR appears to be capable of discriminating between a dual and a single ligand-binding event. Since Vγ9 is often incorporated into non-pAg-reactive Vγ9Vδ1^+^ TCRs, other non-BTN γδ T cell ligands such as MICA, CD1 or MR1 might also co-bind in conjunction with BTN2A1. Likewise, there are numerous parallels between the BTN2A1–BTN3A1 axis and BTN-like proteins, e.g. BTNL3/BTNL8 and Btnl1/Btnl6. It is intriguing that the BTN heterodimers appear to be associating via the same general molecular mechanism. Moreover, the similarity between the BTNL3-Vγ4 and Btnl1-Vγ7 interfaces is compelling, and it will also be interesting to see if the mouse Skint1-Skint2-DETC axis hold similarities ^7,9,13^. However, no role for Ag has been described in those systems, nor have two-ligand binding sites been mapped onto the TCRs. Thus, whilst there is clearly a level of similarity, future studies will need to inform just how homologous those other BTN recognition systems are to that which we define here.

We previously identified three residues in the AEBD face of Vγ9 that are important for mediating the interaction with BTN2A1 ^11^ which we confirm are important in our crystal structure. By contrast, the HV4 and CDR2γ loops, which were proposed by Karunakaran et al. (2020) ^12^ to be involved, do not form part of the interface with BTN2A1. Nonetheless, the binding mode between BTN2A1 and Vγ9Vδ2 TCR is consistent with the notion that germline-encoded sequence variability within the ABED sheet (also termed HV4γ region in some studies) is associated with a spatially distinct epitope that engages BTN molecules ^7^.

Unlike αβTCRs and BCRs, which directly sense foreign Ag, pAg-reactive γδ TCRs are activated by inside-out signalling via BTN conformational changes. As such, additional regulatory mechanisms are likely required to maintain γδ T cell self-tolerance. To this end we identified two important molecular checkpoints, namely Lys53 in the CDR2δ loop of Vγ9Vδ2^+^ TCR, which suppresses BTN3A1 binding, and a second mechanism whereby the Vγ9Vδ2^+^ TCR-binding epitopes of BTN2A1 and BTN3A1 are partnered to each other in cis on the cell surface of APCs, but in a non-activating conformation. The ability of the Lys53δ- Ala mutant TCR to induce Vγ9Vδ2^+^ T cell autoactivation, and stronger BTN3A1 binding, suggests that this residue may have a natural role in regulating Vγ9Vδ2^+^ T cell activation. Upon pAg-induced conformational change of BTN3A1, the Lys53δ side chain steric barrier may be circumvented by the Vγ9Vδ2^+^ TCR, enabling it to engage BTN3A1 via an epitope encompassing Arg51δ, Glu52δ and/or Lys108γ. These residues have been implicated in previous screens as being important for pAg-responsiveness of Vγ9Vδ2 T cells ^11,15^, and moreover, are conserved amongst pAg-reactive human Vγ9Vδ2 TCRs and orthologues of *TRDV2* identified in higher order primates (ensembl.org; **Extended Data Fig. 13A**), as well as pAg-reactive T cells isolated from Rhesus macaque ^22^. We do not exclude a role for other residues identified as being important for pAg-responses based on conservation across pAg-reactive TCRs, such as a strong enrichment of isoleucine, leucine or valine at residue 97 in the adjacent CDR3δ (^23,24^ and **Extended Data Fig. 13B**), although we do note that modification of residue 97 of G115 δ-chain did not heavily impact upon pAg-mediated activation in our previous study ^11^.

Since BTN2A1 and BTN3A1 are both ligands of the Vγ9Vδ2^+^ TCR, yet also directly interact with each other, this may ensure that whilst both ligands remain in an off-state they are proximal to one another such that upon pAg triggering the conversion of the complex into a stimulatory form is rapid and efficient. While the significance of the BTN2A1 V- and head-to-tail dimers remains to be tested, they are reminiscent of the reported BTN3A1 V- and head-to-tail dimers ^17,25^. The stoichiometry of the stimulatory BTN2A1 and BTN3A1 complex needs to be addressed in future studies.

Our data provide three lines of evidence that BTN3A1 is a direct ligand of the Vγ9Vδ2^+^ TCR. Firstly, treatment of BTN3A1-transfected (but not BTN2A/BTN3A-deficient) human or mouse APCs with agonist BTN3A mAb clone 20.1 bind Vγ9Vδ2^+^ TCR tetramers, and do so via a separate Vγ9Vδ2^+^ TCR epitope compared to where BTN2A1 binds. Secondly, recombinant BTN2A1–BTN3A1 complexes bind Vγ9Vδ2^+^ TCR-transfected cells in a way that co-depends on these same dual epitopes. Lastly, the co-binding by BTN2A1–BTN3A1 complexes was recapitulated in cell-free biophysical assays, thus excluding the role of any alternative ligands that bind Vγ9Vδ2^+^ TCR. It is nonetheless important to emphasize that our observations do not preclude the possible involvement of other proteins in driving pAg-dependent activation of Vγ9Vδ2^+^ T cells, nor do they exclude the possibility that BTN3A2 or BTN3A3 can substitute as the TCR ligand in place of BTN3A1. Of note, a recent preprint by Thomas Herrmann and colleagues has established a broader role for the BTN3A isoforms, BTN3A2 and BTN3A3, in the pAg-reactive BTN3A heterodimer (Karunakaran et al., 2023, research square). Previous studies have suggested that the BTN3A isoforms stabilise BTN3A1 expression ^25,26^, however, Herrmann et al. demonstrate a role for the IgV domain of BTN3A2 and BTN3A3 following pAg binding the B30.2 domain of BTN3A1 (Karunakaran et al., 2023, research square). In agreement the notion that other BTN3A isoforms can engage Vγ9Vδ2 TCR, we were able to demonstrate a CDR2δ-mediated interaction of Vγ9Vδ2 TCR tetramers with cells expressing BTN3A2 and BTN3A3, suggesting that TCR sensing by BTN3A isoforms is not limited to BTN3A1 (**Extended Data Fig. 13C**). But our data do demonstrate that BTN2A1 and BTN3A1 are necessary and sufficient for binding Vγ9Vδ2^+^ TCR and pAg responses in gene-transfer assays.

Interestingly, we found that whilst membrane-bound full-length BTN3A1 can bind Vγ9Vδ2^+^ TCR, a soluble form of the BTN3A1^WT^ ectodomain cannot do so unless BTN2A1 is also present, perhaps due to the requirement for a conformational change. However, soluble BTN3A1^Gln45-Ala^ could bind surface-expressed G115^Lys53δ-Ala^, suggesting that the affinity of soluble BTN3A1 for TCR is weak. Whether BTN2A1 induces a conformational change in BTN3A1, or vice versa, is unclear. Our findings suggest that engagement of BTN3A1 by mAb 20.1, or exposure of BTN2A1–BTN3A1-expressing cells to pAg enhances the binding of Vγ9Vδ2 TCR. This is consistent with a model of pAg-induced conformational change in BTN2A1 and/or BTN3A1, which has been proposed as a possible mechanism of inside-out signalling ^21,25,27,28^. However, it is also possible that other pAg-induced effects on the BTN complex may contribute to inside-out signalling, such as localization of BTN molecules into discrete clusters or aggregates. Recombinant BTN2A1–BTN3A1 ectodomain complexes bound Vγ9Vδ2^+^ TCR with a similar affinity to BTN2A1 alone, suggesting that the energetic penalty of having BTN2A1 and BTN3A1 co-liganded to each other is offset by the gain in affinity achieved by having two complementary ligands. Indeed, the enhanced binding affinity of a BTN2A1^WT^–BTN3A1^Glu106Ala^ complex supports this conclusion, and further, also suggests that a single Vγ9Vδ2^+^ TCR molecule can simultaneously co-bind both ligands.

Yuan et al. demonstrate that pAg brings the intracellular domains of BTN2A1 and BTN3A1 together ^21^. Association of the intracellular domains may result in torsional forces that propagate through the rigid coiled-coil domains towards the ectodomains of the BTN complex. Alternatively, another as-yet unidentified molecule might facilitate a conformational change in the BTN complex. This might then enable the Vγ9Vδ2 TCR to first engage BTN2A1 with high affinity, and subsequently bind to BTN3A1 to convey the presence of pAg. Collectively, our findings reveal a fundamentally different molecular mode of immune activation underpins γδ T cell immunity compared to αβ T cells.

## Methods

### Human samples

Healthy donor blood derived human peripheral blood cells (PBMCs) from male and female donors were obtained from the Australian Red Cross Blood Service under ethics approval 17-08VIC-16 or 16-12VIC-03, with ethics approval from University of Melbourne Human Ethics Sub-Committee (1035100) and isolated via density gradient centrifugation (Ficoll-Paque PLUS GE Health care) and red blood cell lysis (ACK buffer, produced in-house).

### Cell lines

Jurkat (JR3-T3.5), LM-MEL-75, HEK293T and NIH-3T3 cells were existing tools in the lab and were maintained in RPMI-1640 (Invitrogen) supplemented with 10% (v/v) FCS (JRH Biosciences), penicillin (100 U/ml), streptomycin (100 μg/ml), Glutamax (2 mM), sodium pyruvate (1 mM), nonessential amino acids (0.1 mM) and HEPES buffer (15 mM), pH 7.2–7.5 (all from Invitrogen Life Technologies), plus 50 μM 2-mercaptoethanol (Sigma-Aldrich) (complete RMPI). Expi293F cells were purchased from ThermoFisher (Cat. No. A14527) and maintained in Expi293 Expression Medium (ThermoFisher, A1435101).

### **γδ** T cell isolation and expansion

In some experiments γδ T cells were enriched by MACS using anti-γδTCR-PECy7 followed by anti-phycoerythrin–mediated magnetic bead purification. After enrichment CD3^+^ Vδ2^+^ γδ T cells were further purified by sorting using an Aria III (BD). Enriched γδ T cells were stimulated *in vitro* for 48 h with plate-bound anti-CD3ε (OKT3, 10 μg/ml, Bio-X-Cell), soluble anti-CD28 (CD28.2, 1 μg/ml, BD Pharmingen), phytohemagglutinin (0.5 μg/ml, Sigma) and recombinant human IL-2 (100 U/ml, PeproTech), followed by maintenance with IL-2 for 14–21 d. Cells were cultured in complete medium consisting of a 50:50 (v/v) mixture of AIM-V (Thermo Fisher) and RPMI-1640 supplemented with 10% (v/v) FCS, penicillin (100 U/ml), streptomycin (100 μg/ml), Glutamax (2 mM), sodium pyruvate (1 mM), nonessential amino acids (0.1 mM) and HEPES buffer (15 mM), pH 7.2–7.5, plus 50 μM 2-mercaptoethanol.

### Flow cytometry

To examine the capacity of γδTCR tetramers to bind to BTN molecules, NIH-3T3 cells were transfected with BTN2A1, BTN3A1 or control BTNL3 in pMIG (a gift from D. Vignali (Addgene plasmid # 52107) ^31^ using ViaFect® (Promega) in OptiMEM™ (Gibco, Thermo-Fisher). 48 h following transfection, cells were harvested with trypsin, filtered through a 30 or 70 µm cell strainer, and incubated with anti-BTN3A antibody (clone 20.1) or IgG1,κ isotype control (clone MOPC-21, BioLegend; or BM4-1, a gift from CSL Limited) at 5 µg/mL for 15 min at room temperature. Cells were then stained with PE-labelled γδTCR tetramers (produced in house, see below), or control PE-conjugated streptavidin, at 5 µg/mL for 30 min at room temperature. The median fluorescence intensity (MFI) of γδTCR tetramer interacting with BTN proteins was examined on gated GFP^+^ cells by flow cytometry. For γδTCR tetramer staining, data were excluded if BTN3A1 mutant protein levels were > 2-fold lower than wild-type BTN3A1, as determined by anti-BTN3A mAb staining. To examine the capacity of BTN tetramers to bind to γδTCRs, HEK293T cells were co-transfected with γδTCR genes in pMIG using FuGENE® HD (Promega) in OptiMEM™ plus 2A-linked CD3εδγζ in pMIG ^32^. 48 h following transfection, cells were collected by pipetting, filtered through a 30 or 70 µm cell strainer, and stained with anti-CD3ε antibody for 15 min at 4°C. Cells were then stained with anti-γδTCR, anti-TCR Vδ2 as well as PE-labelled BTN tetramers (produced in house, see below), PE-labelled control mouse CD1d ectodomain tetramers (loaded with α-GalCer and produced in house, see below), or control PE-conjugated streptavidin (BD), for 30 min at 4°C. The MFI of BTN tetramer on gated CD3^+^GFP^+^ cells was measured by flow cytometry. In other assays, human peripheral blood-derived cells were stained with 7-aminoactinomycin D (7-AAD, Sigma) or LIVE/DEAD® viability markers (ThermoFisher) plus antibodies against: CD3ε, γδTCR, TCR Vδ2, CD45, CD25, CD69, and/or isotype controls (IgG1,κ clone MOPC-2) in various combinations (**Extended Data Table 6**). All data were acquired on an LSRFortessa™ II (BD) and analyzed with FACSDiva and FlowJo (BD) software. All samples were gated to exclude unsTable events, doublets and dead cells using time, forward scatter area versus height, and viability dye parameters, respectively (**Extended Data** Fig. 14).

### Generation of BTN2A.BTN3A-knockout cells

HEK293T cells were nucleofected with Cas9/RNP complexes and two guide RNAs, one targeting the intronic region directly upstream of BTN3A2 (5′- AACTTTCACCTACAAACCGC) and one downstream of BTN2A1 (5′- GAACCCTGACTGAAACGATC). Guides were designed using the Broad Institute CRISPick web tool ^33^. After seven days in culture, BTN2A^-^ BTN3A^-^ cells were bulk-sorted (FACS Aria III) and after another round of culture were single cell-sorted based upon the same criteria. To verify excision of the BTN locus, genotyping of the expanded clones was performed using PCR primers targeting BTN3A2, BTN2A1 and the excised locus (**Extended Data Table 7**).

### Jurkat assays

2.5×10^4^ APCs (LM-MEL-75) cells were plated per well of a 96-well plate and incubated overnight, before 2×10^4^ G115 mutant-expressing J.RT3-T3.5 (Jurkat) cells ± zoledronate (40 μM) were added for 20 h. CD69 expression was measured by flow cytometry on GFP^+^ Jurkat cells. A panel of 15 single-residue alanine (Ala) mutants, each within in the Vδ2 domains of the Vγ9Vδ2^+^ G115 TCR were generated by either site-directed mutagenesis using the primers listed in **Extended Data Table 7**, or by cloning of gene fragments (IDT). Primers (IDT) were phosphorylated (PNK, NEB) followed by 25 cycles of PCR using KAPA HiFi master mix (KAPA Biosystems) using G115 WT TCR in pMIG as template, and PCR product was digested with DpnI (NEB) and in some cases ligated with T4 DNA ligase (NEB). Construct sequences were verified by Sanger sequencing prior to use.

### **γδ** T cell functional assays

For co-culture assays, NIH-3T3 cells were transfected with BTN2A1 in combination with wild-type or mutant BTN3A1, or separately with control BTNL3 and BTNL8 in pMIG with ViaFect® in OptiMEM™. 48 h following transfection, NIH-3T3 cells (3×10^4^) were harvested, transferred to 96-well plates and incubated with purified in vitro-expanded Vδ2^+^ γδ T cells (2×10^4^) for 24 h ± zoledronate (5 μM). γδ T cell activation was determined by CD25 upregulation using flow cytometry. For γδ T cell functional assays, samples were excluded if transfection efficiency was less than 10%.

### Detection of Förster Resonance Energy Transfer

Thirty-thousand NIH-3T3 cells were transfected with BTN2A1 in combination with wild-type or mutant BTN3A1, or control BTN2A1 transfected with PDL2 / BTN3A1 transfected with CD80, in pMIG with ViaFect® in OptiMEM™. 48 h following transfection, NIH-3T3 cells were harvested with trypsin, filtered through 30–70 µm cell strainers, and stained with anti-BTN2A1-AlexaFluor647 (clone 259) and BTN3A-PE (clone 103.2) or isotype controls (clones BM4-2a and MOPC-21, respectively) for 30 min at 4°C. The frequency of cells identified as FRET^+^ was examined on gated GFP^+^AlexaFluor647^+^PE^+^ NIH-3T3 cells. For FRET experiments, data were excluded if BTN3A1 mutant protein levels were > 2-fold lower than BTN3A1 WT as determined by anti-BTN3A (clone 103.2) mAb staining.

### Production of soluble proteins and tetramers

Soluble human BTN2A1–BTN3A1 ectodomains, or alternatively BTN2A1 ectodomains containing a C-terminal Cys (Cys247) and an acidic or basic leucine zipper ^34^, along with soluble γδTCRs, BTN1A1, BTN2A1 lacking Cys247, BTN3A1, BTN3A1 IgV domain, and mouse CD1d ectodomains were expressed by transient transfection of mammalian Expi293F or *MGAT1*^null^ (GNTI) HEK-293S cells using ExpiFectamine or PEI, respectively, with pHL-sec vector DNA encoding constructs with C-terminal biotin ligase (AviTag™) and His_6_ tags ^35^. Protein was purified from culture supernatant using immobilized metal affinity chromatography (IMAC) and gel filtration, and enzymatically biotinylated using BirA (produced in-house). Proteins were re-purified by size exclusion chromatography and stored at -80°C. Biotinylated proteins were tetramerized with streptavidin-PE (BD) at a 4:1 molar ratio.

### Structure determination

BTN2A1 and G115 γδTCR were mixed at a 1:1 molar ratio (15 mg/ml in Tris-buffered saline pH 8) and crystallized at 20°C in 20% polyethylene glycol (PEG) 3350/0.2 M sodium malonate/malonic acid pH 7.0; apo BTN2A1 (10 mg/ml in Tris-buffered saline pH 8) was crystallized at 20°C in 1.65 M ammonium sulfate/2% (v/v) PEG 400/0.1 M HEPES pH 8; and BTN2A1–BTN3A1–zippered complex (1 mg/ml in Tris-buffered saline pH 8) was crystallized at 20°C in 6% (w/v) PEG 6000/0.1 M magnesium sulfate/0.1 M HEPES pH 6 by sitting drop vapour diffusion (C3 facility, CSIRO, Australia). Crystals of BTN2A1–G115 γδTCR, apo BTN2A1 and BTN2A1–BTN3A1-zippered complex were flash frozen in mother liquor plus 27.5% (w/v) PEG/0.2 M sodium malonate, 1.8 M ammonium sulfate/2% (v/v) PEG 400/15% (v/v) glycerol, or in well solution plus 20% (v/v) glycerol, respectively. Data were collected at 100 K using the MX2 (3ID1) beamline at the Australian Synchrotron ^36^ with an Eiger detector operating at 100 Hz. Data were integrated using iMosflm version 7.3.0 ^37^ and, in the case of BTN2A1–G115 γδTCR, processed using the Aimless package in CCP4, or in the case of apo BTN2A1 and BTN2A1–BTN3A1-zippered complex, subjected to the STARANISO Server (Global Phasing Ltd.) (staraniso.globalphasing.org/cgi-bin/staraniso.cgi) to perform an anisotropic cut-off and to apply an anisotropic correction to the data. Apo BTN2A1 was solved by molecular replacement using the IgV and IgC domains of bovine BTN1A1 as separate search ensembles (PDB code 4HH8 ^38^); BTN2A1–G115 γδTCR was solved by molecular replacement using G115 TCR (PDB code 1HXM ^39^) and monomeric BTN2A1; BTN2A1–BTN3A1–zippered complex was solved by molecular replacement using monomeric BTN2A1, and BTN3A1 (from PDB code 4F80 ^17^), with Phaser ^40^. Refinement of BTN2A1–G115 γδTCR was performed by iterative rounds of model building into experimental maps in Coot and refinement with Buster version 2.10.4 (Global Phasing), using non-crystallographic symmetry (NCS) restraints applied to BTN2A1, excluding residues at the TCR-binding interface ^41^. Refinement of apo BTN2A1 and BTN2A1–BTN3A1-zippered complex were similarly restrained against the unliganded copy of BTN2A1 from BTN2A1– G115 γδTCR, or BTN3A1 from 4F80, excluding residues at the interfaces. The BTN2A1–G115 γδTCR, apo BTN2A1 ectodomain and BTN2A1–BTN3A1 ectodomain complex structural models were deposited in the Protein Data Bank under the accession codes 8DFW, 8DFY and 8DFX, respectively, and were analyzed with the CCP4 suite version 7.1 ^42^. Molecular figures were generated with PyMOL (Schrödinger). Cation-π interactions were determined as described ^43^. Angles were calculated between the center of masses of the Ig domains, or in some cases by the intersection of two planes, each defined by three points. Modelling in Extended Data Fig. 11 was performed using AlphaFold 2.0 ^44^. In order to remove model bias, electron density 2mFo-DFc omit maps were generated as a composite of partial models that were refined with simulated annealing using Phenix.

### Enzyme-linked immunosorbent assay (ELISA)

ELISA plates (Microtitre) were coated with 50 µl/well of 5 µg/ml NeutrAvidin (Thermo Fisher Scientific, #31000) for 1 h at room temperature, washed with 0.05% v/v Tween-20 in PBS (PBST), blocked with 1% w/v BSA in PBS for 1 h at room temperature, and biotinylated BTN proteins captured. Alternatively, BTN proteins were coated directly on to an ELISA plate, washed with PBST and blocked with 1% BSA in PBS for 1 h at room temperature. BTN proteins were then probed with 2 µg/ml anti-BTN2A1 (mAb clone 259), anti-BTN3A (mAb clones 103.2 or 20.1 as indicated) or appropriate isotype control mAb BM4-2a (IgG2a,κ; a gift from CSL Limited) or MOPC-21 (IgG1,κ) and detected with goat anti-mouse Ig-HRP (Thermo Fisher Scientific, #63-65-20). 50 µl/well of TMB (Thermo Fisher Scientific; 1-Step Ultra TMB-ELISA) was added and incubated for ∼10 min, and subsequently quenched with 2.25 M HCl, followed by measuring absorbance at 450 nm on a ClariostarPlus (BMG LabTech). Data were analysed in GraphPad Prism.

### Surface plasmon resonance

SPR experiments were conducted at 25°C on a Biacore T200 instrument (GE Healthcare) using 10 mM HEPES-HCl (pH 7.4), 150 mM NaCl, 3 mM EDTA, and 0.05% Tween 20 buffer. Biotinylated BTN ectodomains were immobilized to 1,500-2,000 resonance units (RU) on a Biacore sensor chip SA pre-immobilized with streptavidin. Soluble BTN molecules or G115 γδTCR were two-fold serially diluted and simultaneously injected over test and control surfaces at a rate of 30 μl/min. After subtraction of data from the control flow cell (BTN1A1) and blank injections, interactions were analyzed using Biacore T200 evaluation software (GE Healthcare), Scrubber (Biologic) and Prism version 9 (GraphPad), and equilibrium dissociation constants were derived at equilibrium.

### Biolayer Interferometry (BLI)

Affinity measurements for the BTN2A1-BTN3A1 interaction were determined by BLI on an Octet Red 96e instrument (Sartorius). Assays were performed in black 96 well flat-bottom plates at 25°C with agitation (1000 rpm), using sample buffer 1 (150 mM NaCl, 10 mM HEPES, 0.05% Tween-20, 3 mM EDTA, pH 7.4), sample buffer 2 (150 mM NaCl, 10 mM HEPES, 0.05% Tween-20, 3 mM EDTA, 1 mM DTT, pH 7.4) and regeneration buffer (1 M NaCl, 10 mM HEPES pH 7.4). First, Streptavidin Biosensors (Sartorius) were hydrated for 20 minutes at 100 rpm in sample buffer 1. Biotinylated BTN3A1-B30.2 was then loaded at 5 µg/ml in sample buffer 1 to a ligand coating density of 1.2 nm, and a baseline signal was established for 180 seconds in sample buffer 1. For series in the absence of phosphoantigen, a second baseline signal was established in sample buffer 2 before dipping sensors into 2-fold serial dilutions of BTN2A1-BFI (wildtype or mutant) in sample buffer 2 starting at 10 µM. A 360 second association phase was recorded, followed by 2 separate 600 second dissociation phases in different wells of buffer 2. Sensors were then regenerated by three cycles of immersion in regeneration buffer followed by buffer 2 for 90 seconds per step, and assay steps were repeated for remaining sample wells. For series in the presence of phosphoantigen (20 µM HMBPP), a baseline signal was established in wells containing sample buffer 2 with HMBPP ahead of ligand association in BTN2A1-BFI (wildtype or mutant) along with HMBPP. This was followed by a dissociation phase in buffer 2 with HMBPP, and then a further dissociation phase in buffer 2 without HMBPP. Reference sensor data from the BTN3A1-B30-coated sensor dipped in buffer 2 alone was subtracted to account for baseline drift, and the data were aligned to the y-axis at baseline. Kinetic constants were determined using the Octet Data Analysis HT software v12.0.2.59 (ForteBio). Curve fitting was performed using a 1:1 global fit biding model.

### Electron microscopy

Soluble BTN2A1 Gly130-Cys–BTN3A1 Asp132-Cys complex was enzymatically digested with thrombin to remove C-terminal leucine zippers, repurified by size exclusion and anion exchange chromatography, and spotted onto glow-discharged 400 mesh thin carbon-coated copper grids at 380 μg/ml in TBS for 30 seconds, followed by negative staining with 2% w/v uranyl acetate. Grids were observed on a FEI Tecnai F30 (Eindhoven, NL) 300 kV transmission electron microscope at a nominal magnification of ×52,000. Seventeen micrographs were acquired on a CETA (Thermofisher, USA) camera with a 3.7 Å pixel size. Particles were picked using blob picking followed by 2D class averaging in cryoSPARC ^45^, with 10,238 particles contributing to the final set of 2D class averages.

### Statistical analysis

γδ T cell functional assays were analysed by 2-way ANOVA with Šidák’s correction when comparing γδ T cell activation (CD25^+^) with and without treatment across various BTN mutants. Change in His85γ-Ala tetramer MFI staining with Zol treatment was analysed by Wilcoxon matched-pairs signed rank test. All independent datapoints are biological replicates.

### Data and materials availability

Reagents used in this study are available upon request. The crystal structures are deposited in the Protein Data Bank under accession codes 8DFW, 8DFY and 8DFX.

## Acknowledgements

We thank Con Panousis, Geoffrey Matthews and staff at CSL Limited for providing anti-BTN2A1 mAb. We thank Meghan Hattarki from CSIRO for support with SPR. We thank staff at the C3 protein crystallisation facility (CSIRO), the Ian Holmes Imaging Centre (Bio21 Institute, University of Melbourne), the MX beamlines at the Australian Synchrotron, the Melbourne Cytometry Platform (University of Melbourne), and AGRF for DNA sequencing support. This research was undertaken in part using the MX2 beamline at the Australian Synchrotron, part of ANSTO, and made use of the ACRF detector. Infrastructure support from the NHMRC Independent Research Institutes Infrastructure Support Scheme and the Victorian State Government Operational Infrastructure Support Program are gratefully acknowledged. This work was supported by a Miller Foundation Research Accelerator fund (to APU), Cancer Council of Victoria (1126866), the National Health and Medical Research Council of Australia (NHMRC; 1184906, 1165467); the Australian Research Council (ARC; CE140100011). MR is supported by a Cancer Council Victoria Postdoctoral Research Fellowship. AB is supported by a fellowship from the Department of Health and Human Services acting through the Victorian Cancer Agency. MWP is supported by an NHMRC Investigator Fellowship (APP1194263). NAG is supported by an ARC Discovery Early Career Researcher Award (DE210100705). DIG was supported by NHMRC Senior Principal Research Fellowship (1117766) and subsequently by an NHMRC investigator grant (2008913).

## Author Contributions

Conceptualization, APU, DIG, TSF, NAG, AB; methodology, TSF, CS, MR, OD, SL, AC, TSP, NAG, APU; investigation, TSF, CS, RGC, MR, OD, HGB, EH, APU; resources, ZR, RS, AH, SL, SJR, MAG, MWP, OP, JN, AB, NAG, DIG, APU; writing – original draft, APU, TSF; writing – reviewing & editing, APU, DIG, TSF, NAG, CS, TSP, MR, MWP, OP, AH; supervision, APU, DIG, TSF; funding acquisition, APU, DIG, AB.

## Additional Information

Correspondence and requests for materials should be addressed to Dr Adam Uldrich or Prof Dale Godfrey. Reprints and permissions information is available at www.nature.com/reprints.

## Extended Data for

**Extended Data Table 1.** Data collection and refinement statistics.

|  | **γδTCR–BTN2A1** | **Apo BTN2A1** | **BTN2A1–BTN3A1** |
| --- | --- | --- | --- |
| PDB ID code | 8DFW | 8DFY | 8DFX |
| **Data collection** |  |  |  |
| Radiation source | Australian Synch. MX2 | Australian Synch. MX2 | Australian Synch. MX2 |
| Wavelength (Å) | 0.95372 | 0.95372 | 0.95372 |
| Resolution range (Å) | 49.7–2.10 (2.14–2.10) | 49.8–3.55 (3.89–3.55) | 47.4–5.55 (6.62–5.55) |
| Ellipsoid resolution (Å) (direction)<sup>a</sup> | - | 4.30 (0.95 a*+0.33 c*) | 5.93 (a*) |
|  | - | 3.92 (b*) | 5.28 (b*) |
|  | - | 3.55 (-0.80 a*+0.60 c*) | 8.24 (c*) |
| Best/Worst diffraction limits (Å)<sup>a</sup> | - | 3.55 / 5.62 | 5.55 / 8.85 |
| Space group | C 2 2 2<sub>1</sub> | C 2 | F 2 2 2 |
| Unit cell parameters: a, b, c (Å) | 112.0, 218.5, 107.9 | 237.0, 94.2, 134.7 | 114.6 138.9 336.2 |
| α, β, γ (°) | 90, 90, 90 | 90, 106.35, 90 | 90, 90, 90 |
| **Data processing** |  |  |  |
| Total observations (N<sup>o</sup>) | 1,038,721 (53,094) | 101,471 (5,622)<sup>a</sup> | 18,589 (2,124)<sup>a</sup> |
| Unique observations (N<sup>o</sup>) | 77,369 (3,834) | 22,300 (1,115)<sup>a</sup> | 2,500 (278)<sup>a</sup> |
| Multiplicity | 13.4 (13.8) | 4.6 (5.0)<sup>a</sup> | 7.4 (7.6)<sup>a</sup> |
| Data completeness (isotropic, %) | 99.97 (100.00) | 64.3 (13.5)<sup>a</sup> | 56.5 (15.6)<sup>a</sup> |
| Data completeness (ellipsoidal, %)<sup>a</sup> | - | 85.2 (70.0)<sup>a</sup> | 85.9 (49.2)<sup>a</sup> |
| Mean I/σ<sub>I</sub> | 13.8 (2.8) | 5.8 (2.0)<sup>a</sup> | 5.3 (1.6)<sup>a</sup> |
| Wilson B-factor (Å<sup>2</sup>) | 36 | 111 | 186 |
| R<sub>merge</sub> (%) | 12.0 (108) | 15.3 (78.6)<sup>a</sup> | 20.2 (145.6)<sup>a</sup> |
| R<sub>meas</sub> (%) | 12.4 (112.1) | 17.3 (87.5)<sup>a</sup> | 22.1 (156.2)<sup>a</sup> |
| R<sub>pim</sub> (%) | 3.37 (29.9) | 10.8 (51.3)<sup>a</sup> | 8.6 (55.8)<sup>a</sup> |
| CC<sub>1/2</sub> (%) | 99.8 (72.5) | 98.0 (64.0)<sup>a</sup> | 91.9 (37.3)<sup>a</sup> |
| **Refinement statistics** |  |  |  |
| Resolution range (Å)<sup>b</sup> | 49.7–2.10 (2.14–2.10) | 49.8–3.55 (3.76–3.55) | 47.4–5.55 (6.88–5.55) |
| Reflections used in refinement<sup>b</sup> | 77,362 (3,834) | 22,300 | 2,500 |
| Reflections used for R-free<sup>b</sup> | 3,723 (189) | 1,144 | 226 |
| R<sub>work</sub> (%)<sup>b</sup> | 26.1 (33.6) | 25.5 | 29.0 |
| R<sub>free</sub> (%)<sup>b</sup> | 28.7 (35.9) | 26.8 | 33.6 |
| Number of non-hydrogen atoms | 7,186 | 8993 | 3305 |
| - macromolecules | 6,848 | 8524 | 3277 |
| - ligands | 173 | 469 | 28 |
| - solvent | 165 | 0 | 0 |
| Protein residues | 868 | 1087 | 425 |
| RMSD (bond length)<sup>b</sup> | 0.008 | 0.008 | 0.007 |
| RMSD (bond angle)<sup>b</sup> | 1.02 | 0.99 | 0.91 |
| Ramachandran - favored (%) | 96.4 | 96.8 | 97.6 |
| - allowed (%) | 3.5 | 3.2 | 2.4 |
| - outliers (%) | 0.1 | 0 | 0 |
| Rotamer outliers (%) | 3.7 | 4.4 | 3.1 |
| Clashscore | 5.2 | 6.9 | 8.7 |
| Average B-factor | 58 | 269 | 278 |
| - macromolecules | 58 | 268 | 278 |
| - [per chain] | [57,54,54,68] | [248,246,280,283,287] | [277,279] |
| - ligands | 70 | 281 | 266 |
| - solvent | 42 | - | - |

<sup>a</sup> Calculated by the Staraniso server (Global Phasing). The cut-off surface is unlikely to be perfectly ellipsoidal, so this is only an estimate.
<sup>b</sup> Calculated by Buster (Global Phasing).

**Extended Data Table 2.** BTN2A1 V-dimer contacts.

| BTN2A1 copy #1 region | BTN2A1 copy #1 residues | BTN2A1 copy #2 region | BTN2A1 copy #2 residues | Bond type |
| --- | --- | --- | --- | --- |
| IgC A-strand | Met125 | Juxta-membrane | Pro209, Phe212 | VDW |
| IgC A-strand | Arg126 | IgC G-strand | Phe207 | Cation-π |
| IgC A-strand | Arg126 | IgC G-strand, juxta-membrane | Phe207, Pro209 | VDW |
| IgC A-strand | Gly127 | Juxta-membrane | Pro209, Phe212 | VDW |
| IgC A-strand | His128-Nδ1 | Juxta-membrane | Ser211 Oγ | HB |
| IgC A-strand | His128-N | Juxta-membrane | Ser211-Oγ | Water-mediated HB |
| IgC A-strand | His128 | Juxta-membrane | Ser211, Phe212 | VDW |
| IgC G-strand | Phe207 | IgC A-strand | Arg126 | Cation-π |
| IgC G-strand | Phe207 | IgC A-strand | Arg126 | VDW |
| Juxta-membrane | Pro209 | IgC A-strand | Met125, Arg126, Gly127 |  |
| Juxta-membrane | Ser211-Oγ | IgC A-strand | His128-Nδ1 | HB |
| Juxta-membrane | Ser211-Oγ | IgC A-strand | His128-N | Water-mediated HB |
| Juxta-membrane | Ser211 | IgC A-strand | His128 | VDW |
| Juxta-membrane | Phe212 | Juxta-membrane | Phe212 | π-π stacking |
| Juxta-membrane | Phe212 | IgC A-strand | Met125, Gly127, His128 | VDW |

Van der Waals (VDW) defined as non-hydrogen bond contact distances of 4 Å or less, hydrogen bonds (HB) 3.5 Å or less, cation-π and $\pi$ - $\pi$ as 4.5 Å or less and water-mediated HB 3.3 Å or less.

**Extended Data Table 3.** BTN2A1 head-to-tail dimer contacts.

| BTN2A1 copy #1 region | BTN2A1 copy #1 residues | BTN2A1 copy #2 region | BTN2A1 copy #2 residues | Bond type |
| --- | --- | --- | --- | --- |
| N-terminus | Gln1-Nε2 | IgC D-strand | Lys162-Nζ | HB |
| N-terminus | Gln1 | IgC A/D-strands | Glu129, Lys162 | VDW |
| IgV C-strand | Phe39 | IgC D-strand | Met166, Pro167 | VDW |
| IgV CC'-loop | Ser41 | IgC D/E-strands | Pro167, Gly171 | VDW |
| IgV CC'-loop | Gln42-Nε2 | IgC DE-loop | Ala169-O | HB |
| IgV CC'-loop | Gln42 | IgV EF-loop, IgC D-strand, IgC DE-loop, IgC E-strand | Gln90, Asp168, Ala169, Gly171 | VDW |
| IgV CC'-loop | Phe43-N | IgC D-strand | Asp168-O | HB |
| IgV CC'-loop | Phe43 | IgC D-strand, IgC DE-loop | Met166, Pro167, Asp168, Ala169 | VDW |
| IgV CC'-loop | Ser44 | IgC DE-loop | Ala169 | VDW |
| IgV EF-loop | Gln90 | IgV CC'-loop | Gln42 | VDW |
| IgV glycans | (Asn92) NAG221-O6 | IgV glycans | (Asn92) NAG221-O6 | HB |
| IgV glycans | (Asn92) NAG221 | IgV glycans | (Asn92) NAG221 | VDW |
| IgV F-strand | Arg96 | IgC D/E-strands | Pro167, Phe173 | VDW |
| IgV F-strand | Tyr98 | IgC D-strand | Met166 | VDW |
| IgV FG-loop | Arg103-Nε | IgC B-strand | Glu136-Oε1, Glu136-Oε2 | SB |
| IgV FG-loop | Arg103-Nη1 | IgC B-strand | Glu136-Oε1 | SB |
| IgV FG-loop | Arg103 | IgC A/B-strands | Arg126, Glu136 | VDW |
| IgV G-strand | Tyr105 | IgC D/E-strands | Val164, Met174, Thr176 | VDW |
| IgV G-strand | Asp106 | IgC D-strand | Val164 | VDW |
| IgV G-strand | Glu107 | IgC D-strand | Ser165 | VDW |
| IgC A-strand | Arg126 | IgV FG-loop | Arg103 | VDW |
| IgC A-strand | Glu129 | N-terminus | Gln1 | VDW |
| IgC B-strand | Glu136-Oε1 | IgV FG-loop | Arg103-Nε, Arg103-Nη1 | SB |
| IgC B-strand | Glu136-Oε2 | IgV FG-loop | Arg103-Nε | SB |
| IgC B-strand | Glu136 | IgV FG-loop | Arg103 | VDW |
| IgC D-strand | Lys162 | N-terminus | Gln1 | VDW |
| IgC D-strand | Val164 | IgV G-strand | Tyr105, Asp106 | VDW |
| IgC D-strand | Ser165-N | IgV G-strand | Glu107-Oε1 | Water-mediated HB |
| IgC D-strand | Ser165 | IgV G-strand | Glu107 | VDW |
| IgC D-strand | Met166 | IgV C-strand, CC'-loop, F-strand | Phe39, Phe43, Tyr98 | VDW |
| IgC D-strand | Pro167 | IgV C-strand, CC'-loop, F-strand | Phe39, Ser41, Phe43, Arg96 | VDW |
| IgC D-strand | Asp168-O | IgV CC'-loop | Phe43-N | HB |
| IgC D-strand | Asp168 | IgV CC'-loop | Gln42, Phe43 | VDW |
| IgC DE-loop | Ala169-O | IgV CC'-loop | Gln42-Nε2 | HB |
| IgC DE-loop | Ala169 | IgV CC'-loop | Gln42, Phe43,<br>Ser44 | VDW |
| IgC E-strand | Gly171 | IgV CC'-loop | Ser41, Gln42 | VDW |
| IgC E-strand | Phe173 | IgV F-strand | Arg96 | VDW |
| IgC E-strand | Met174 | IgV G-strand | Tyr105 | VDW |
| IgC E-strand | Thr176 | IgV G-strand | Tyr105 | VDW |

Van der Waals (VDW) defined as non-hydrogen bond contact distances of 4 Å or less, hydrogen bonds (HB) as 3.5 Å or less, salt bridge (SB) as 4.5 Å or less and water-mediated HB 3.3 Å or less.

**Extended Data Table 4.** Vγ9 TCR contacts with BTN2A1.

| Vγ9 TCR region | Vγ9 TCR residue | BTN2A1 IgV region | BTN2A1 residue | Bond type |
| --- | --- | --- | --- | --- |
| A-strand | Lys13γ-Nζ | G-strand | Tyr105-OH | HB |
| A-strand | Lys13γ | G-strand | Tyr105 | VDW |
| B-strand | Ser16γ-O | F-strand | Arg96-Nη2 | HB |
| B-strand | Ser16γ-O | G-strand | Glu107-Oε2 | HB |
| B-strand | Ser16γ | F/G strands | Arg96, Glu107, Ile109 | VDW |
| B-strand | Lys17γ-Nζ | G-strand | Asp106-Oδ1, Asp106-Oδ2 | SB |
| B-strand | Lys17γ-Nζ | G-strand | Glu107-O | HB |
| B-strand | Lys17γ | G-strand | Asp106, Glu107 | VDW |
| B-strand | Thr18γ-N | G-strand | Glu107-Oε1 / Glu107-Oε2 | HB |
| B-strand | Thr18γ-Oγ1 | G-strand | Glu107-Oε1 | HB |
| B-strand | Thr18γ | C/F/G-strands | Phe39, Arg96, Tyr105, Glu107 | VDW |
| B-strand | Ala19γ | G-strand | Tyr105 | VDW |
| B-strand | Arg20γ-N | G-strand | Tyr105-OH | HB |
| B-strand | Arg20γ-O | G-strand | Tyr105-OH | HB |
| B-strand | Arg20γ-Nε | F-strand | Gln100-Nε2 | Water-mediated HB |
| B-strand | Arg20γ | CC'-loop, F/G-strands | Phe43, Tyr98, Tyr105 | VDW |
| B-strand | Arg20γ | CC'-loop | Phe43 | Cation-π |
| C''-strand | Gly67γ-O | CC'-loop | Ser41-O | Water-mediated HB |
| C''-strand | Gly67γ | CC'-loop | Gln42 | VDW |
| D-strand | Glu70γ-Oε1 | CC'-loop | Phe43-N | HB |
| D-strand | Glu70γ-Oε2 | CC'-loop | Ser44-N | HB |
| D-strand | Glu70γ | CC'-loop | Gln42, Phe43, Ser44 | VDW |
| E-strand | Thr83γ | CC'-loop | Phe43 | VDW |
| E-strand | His85γ | CC'-loop | Ser41, Gln42, Phe43 | VDW |
| E-strand | Asn86 | F-strand | Arg96 | VDW |

Van der Waals (VDW) defined as non-hydrogen bond contact distances of 4 Å or less, hydrogen bonds (HB) 3.5 Å or less, cation-π and salt bridge (SB) as 4.5 Å or less and water-mediated HB 3.3 Å or less.

**Extended Data Table 5.**
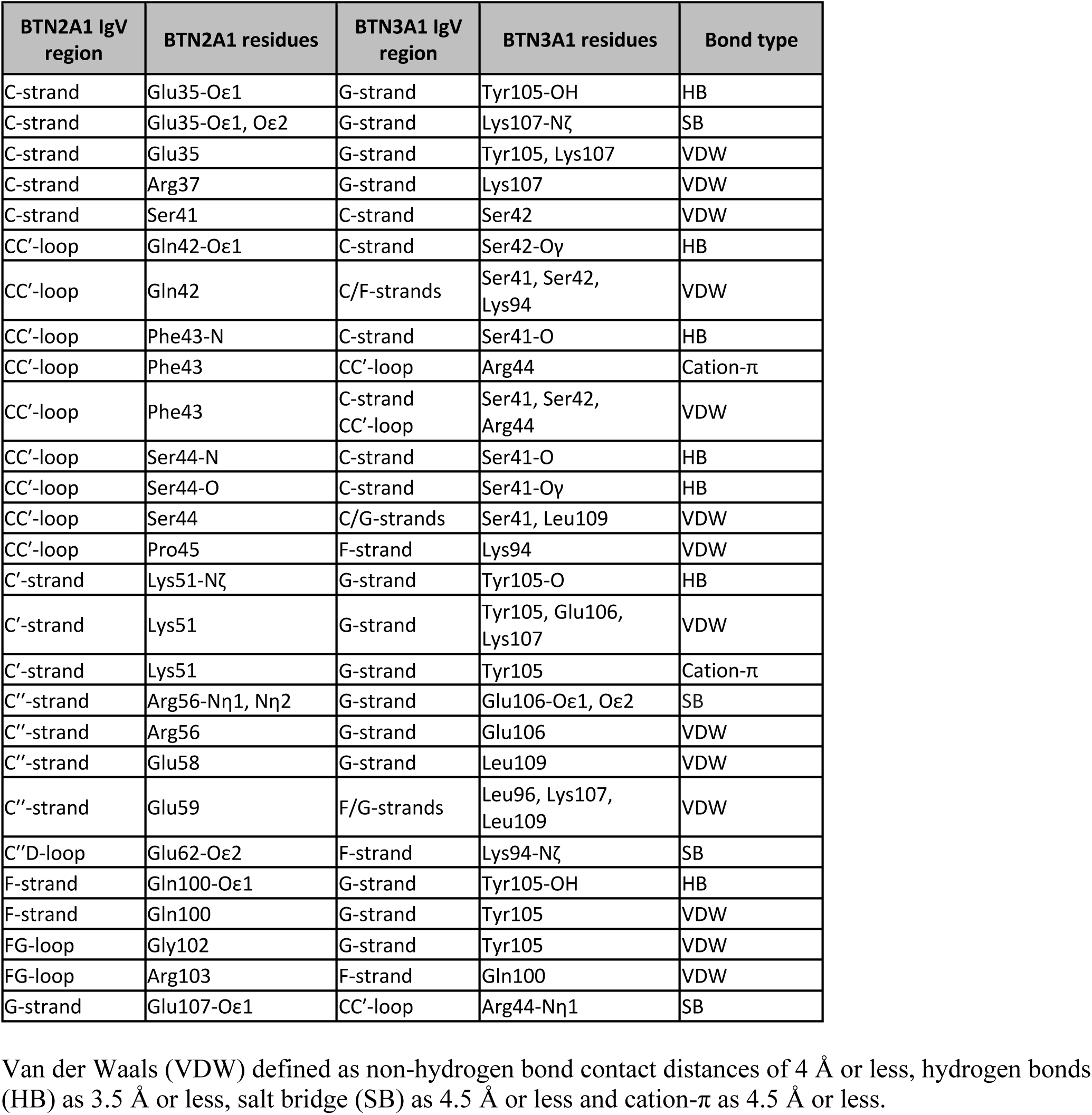
Putative BTN2A1 ectodomain contacts with BTN3A1 ectodomain.

**Extended Data Table 6.**
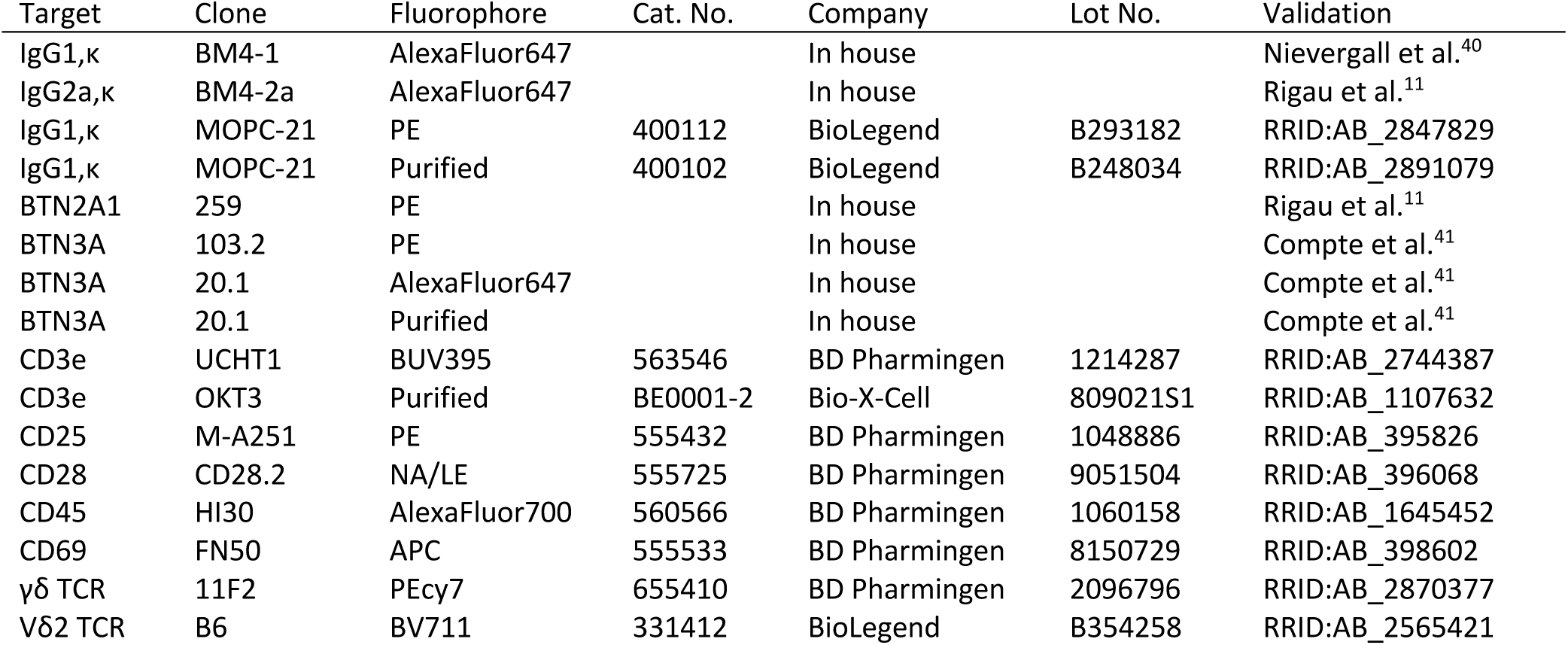
Antibodies used for Flow Cytometry and T cell expansion.

**Extended Data Table 7.**
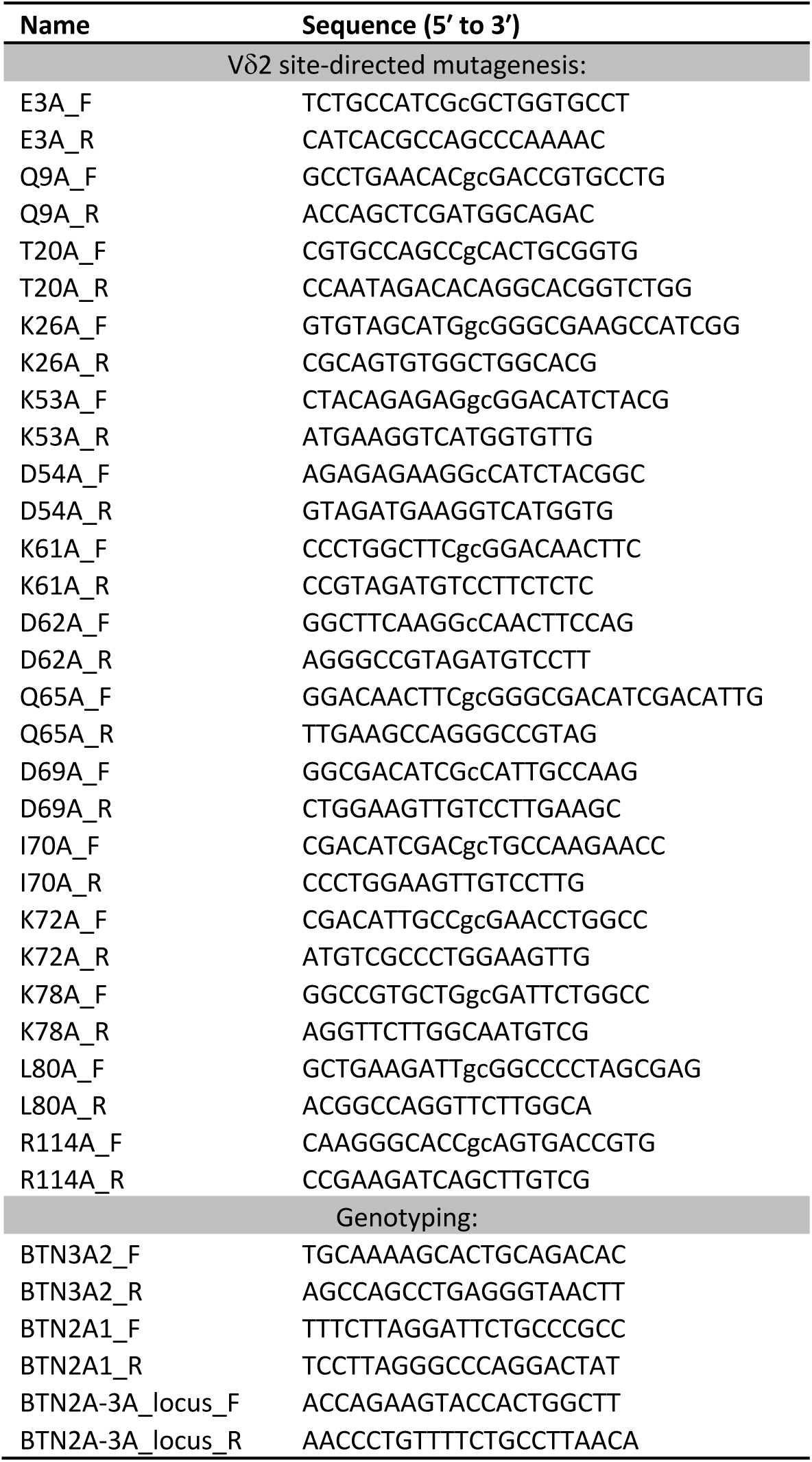
Primers used for PCR and site-directed mutagenesis.

**Extended Data Fig. 1.**
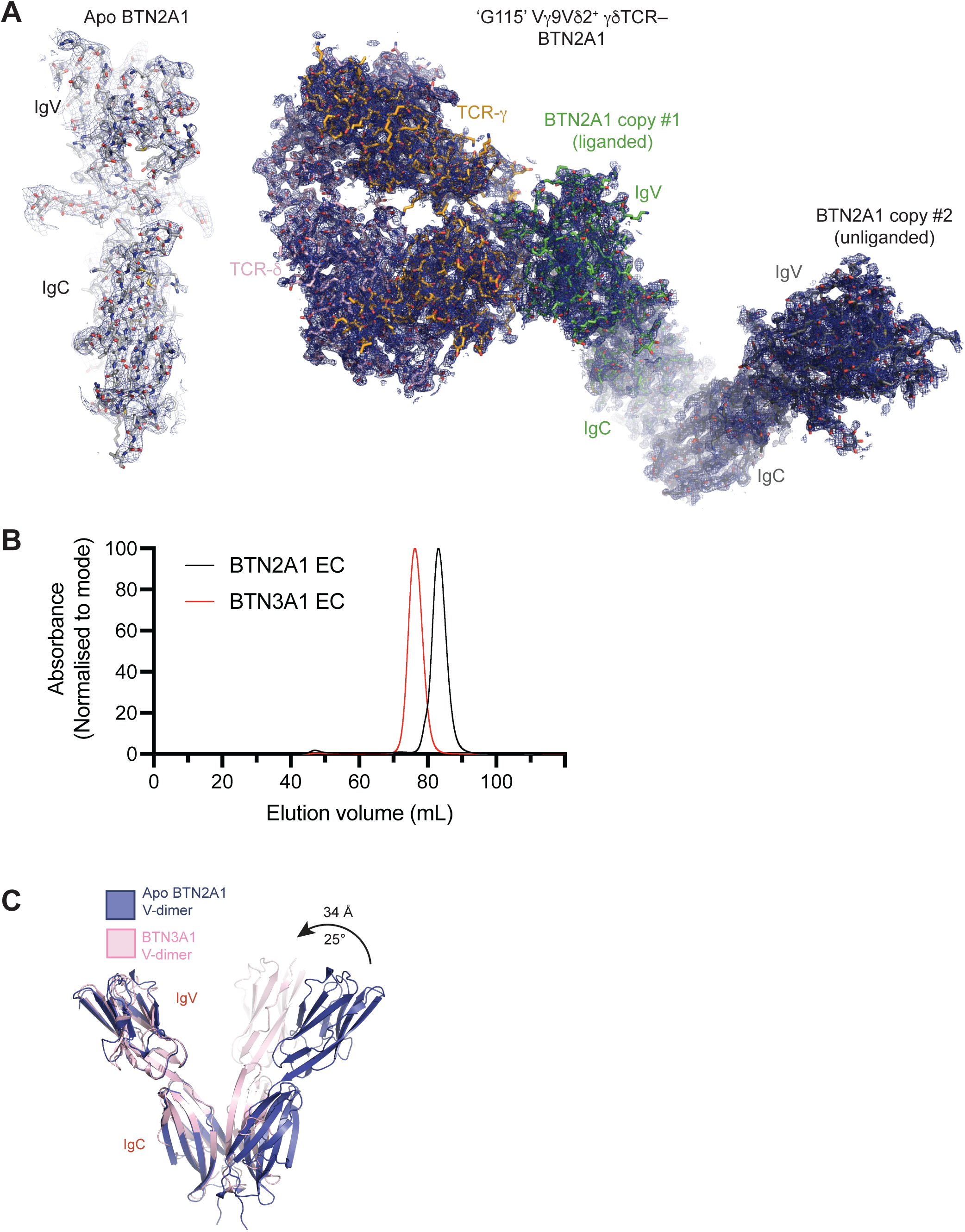
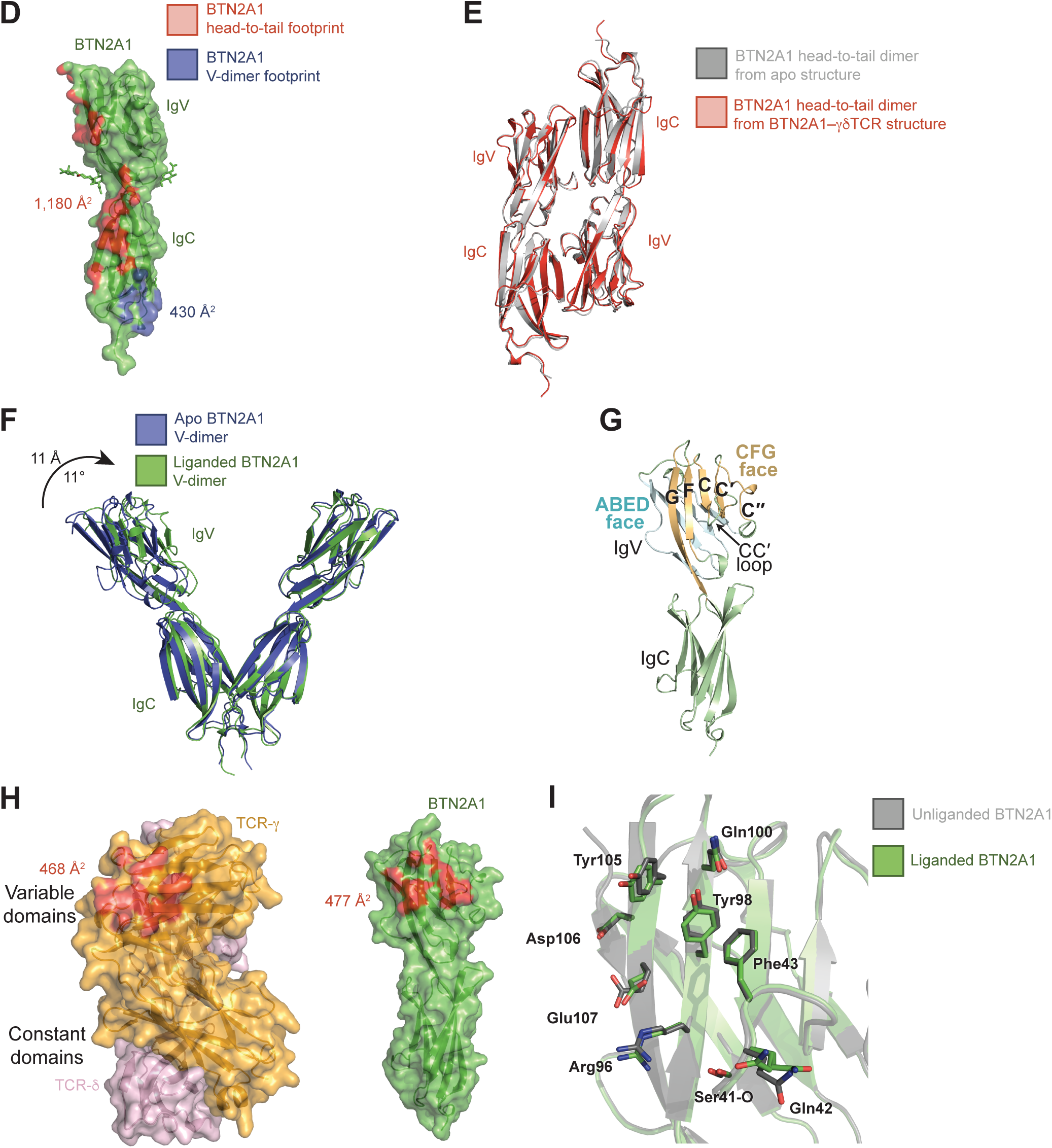
(**A**) Unbiased 2mFo-DFc electron density composite omit map contoured at 1σ, blue mesh, surrounding apo BTN2A1 (averaged over five copies of BTN2A1), and γδTCR-BTN2A1 complex. (**B**) Size exclusion (S200 16/600) gel filtration chromatogram of BTN2A1 (black) and BTN3A1 (red) ectodomains produced in *MGAT1*-deficient Expi293F cells. Larger elution volume indicates smaller protein size. (**C**) Overlay of BTN2A1 V-dimer from apo structure (blue) and BTN3A1 V-dimer (pink) structures (PDB code 4F80). (**D**) Surface representation of BTN2A1 (green) depicting the head-to-tail dimer interface in red and the V-dimer interface in blue. Glycans depicted as sticks. (**E**) Overlay of BTN2A1 head-to-tail dimers derived from the apo BTN2A1 (grey) and γδTCR-BTN2A1 (unliganded copy of BTN2A1, red) crystal structures. (**F**) Overlay of BTN2A1 V-dimer structures derived from the apo (blue) and γδTCR-liganded (green) crystal structures. (**G**) Cartoon of BTN2A1 depicting the CFG face of the IgV domain in gold and the ABED face of the IgV domain in blue (**H**) Surface representation of BTN2A1 (green) and G115 Vγ9Vδ2 TCR (γ-chain, orange; δ-chain, light pink) depicting the interfaces (red) from the γδTCR-BTN2A1 complex. (**I**) Cartoon overlay of apo (grey) and liganded BTN2A1 (green), depicting the conformational changes to residues that are involved in binding to Vγ9Vδ2^+^ TCR (shown as sticks).

**Extended Data Fig. 2.**
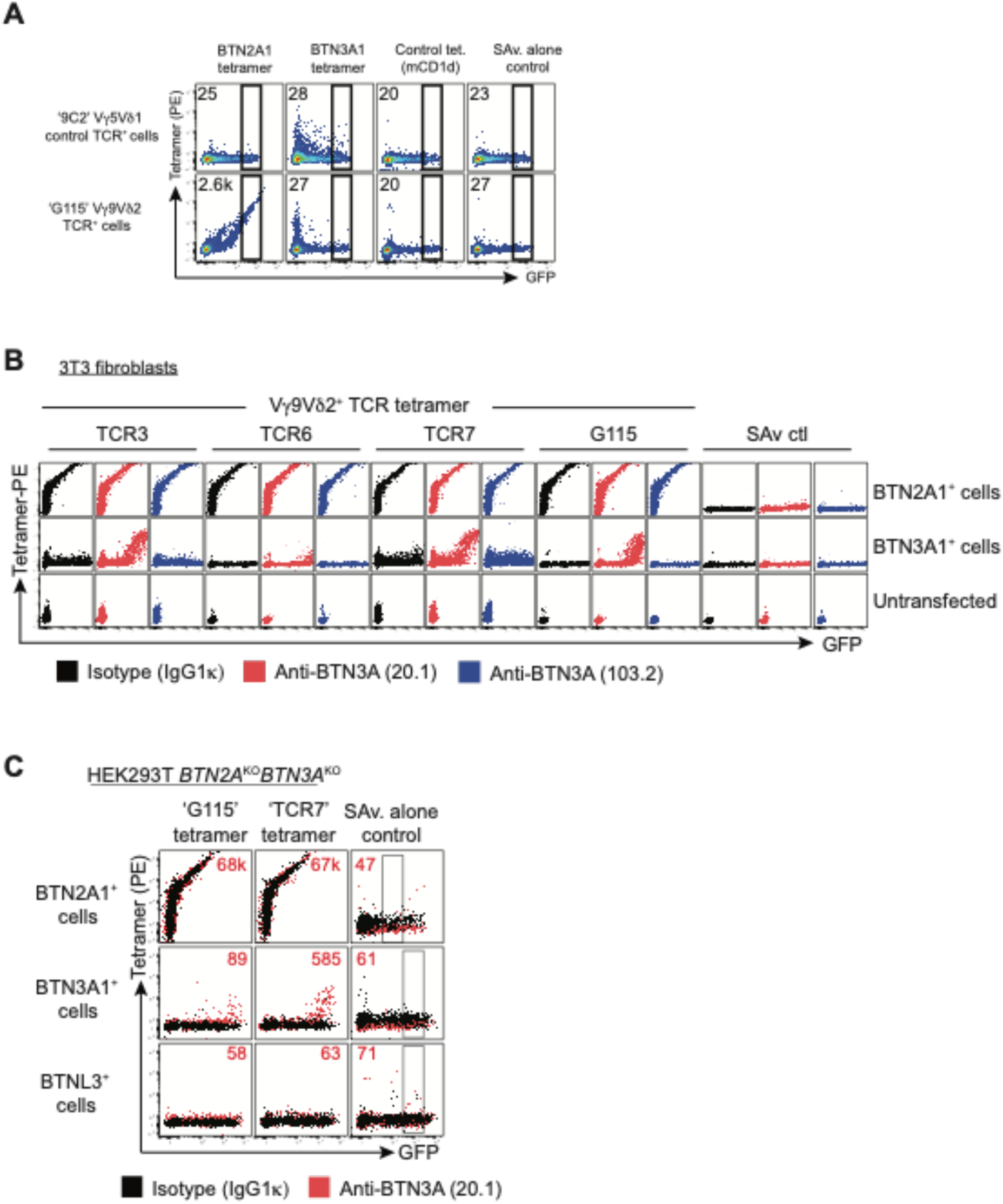
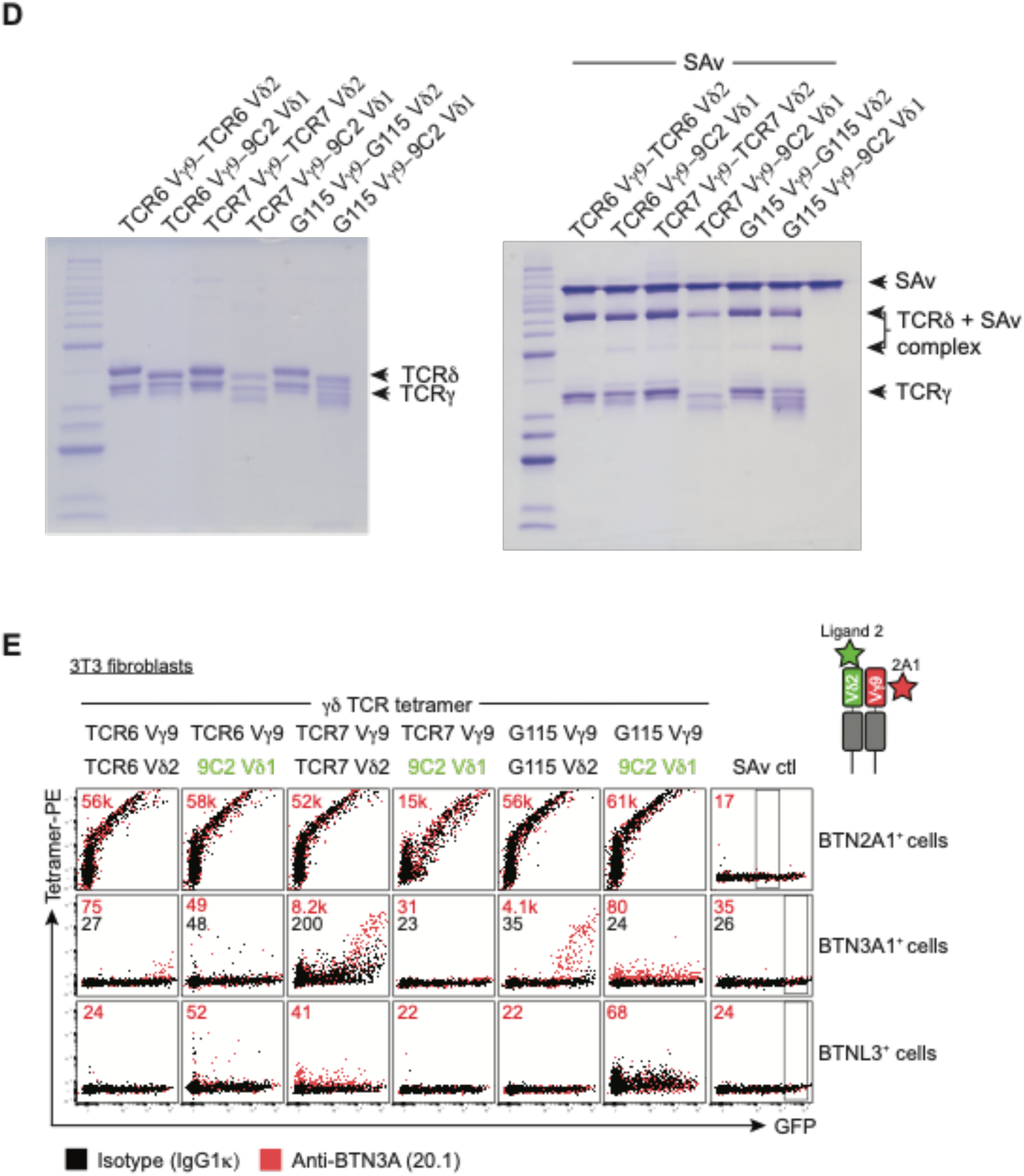
(**A**) BTN2A1 tetramer, BTN3A1 tetramer, control mouse CD1d tetramer, or SAv-PE staining of human HEK293T cells transfected with plasmids co-encoding GFP and either G115 Vγ9Vδ2^+^ or control 9C2 Vγ5Vδ1^+^ γδTCRs. Plots gated on GFP^+^ cells. Data from one of 10 independent experiments. Inset – median fluorescence intensity (MFI) of PE parameter. (**B**) Vγ9Vδ2^+^ TCR tetramer-PE (clones TCR3, TCR6, TCR7 and G115) or streptavidin (SAv.) control staining of mouse NIH-3T3 cells transfected with BTN2A1, BTN3A1 or no DNA following pre-incubation with anti-BTN3A mAb clones 20.1 (red) or 103.2 (blue), or isotype control (IgG1,κ, black). (**C**) γδTCR tetramer or SAv control staining of HEK293T *BTN2A.BTN3A^KO^*cells transfected with plasmids co-encoding GFP and either BTN2A1, BTN3A1 or control BTNL3, which were pre-incubated with anti-BTN3A mAb 20.1 or isotype control (mouse IgG1,κ) antibody. Plots gated on GFP^+^ cells. Inset – MFI of PE parameter of mAb 20.1-treated cells within the representative GFP^+^ gate. Representative of one of two independent experiments. (**D**) SDS-PAGE separation of wild-type and Vδ1–chimeric γδ TCRs alone (**LHS**) or mixed with excess SAv (**RHS**). (**E**) Staining of BTN2A1, BTN3A1 or control BTNL3-transfected NIH-3T3 cells with chimeric γδTCR tetramers comprised of the TCR6, TCR7 or G115 pAg-reactive γ-chains, plus either the pAg-reactive Vδ2^+^ or the 9C2 Vδ1^+^ δ-chains ± anti-BTN3A mAb 20.1 (red) or isotype control (IgG1,κ, black). Median fluorescence intensity (MFI) of PE for mAb 20.1-treated cells (red numbers) or isotype control (IgG1,κ)-treated BTN3A1^+^ cells (black numbers) shown within the depicted GFP^+^ gate.

**Extended Data Fig. 3.**
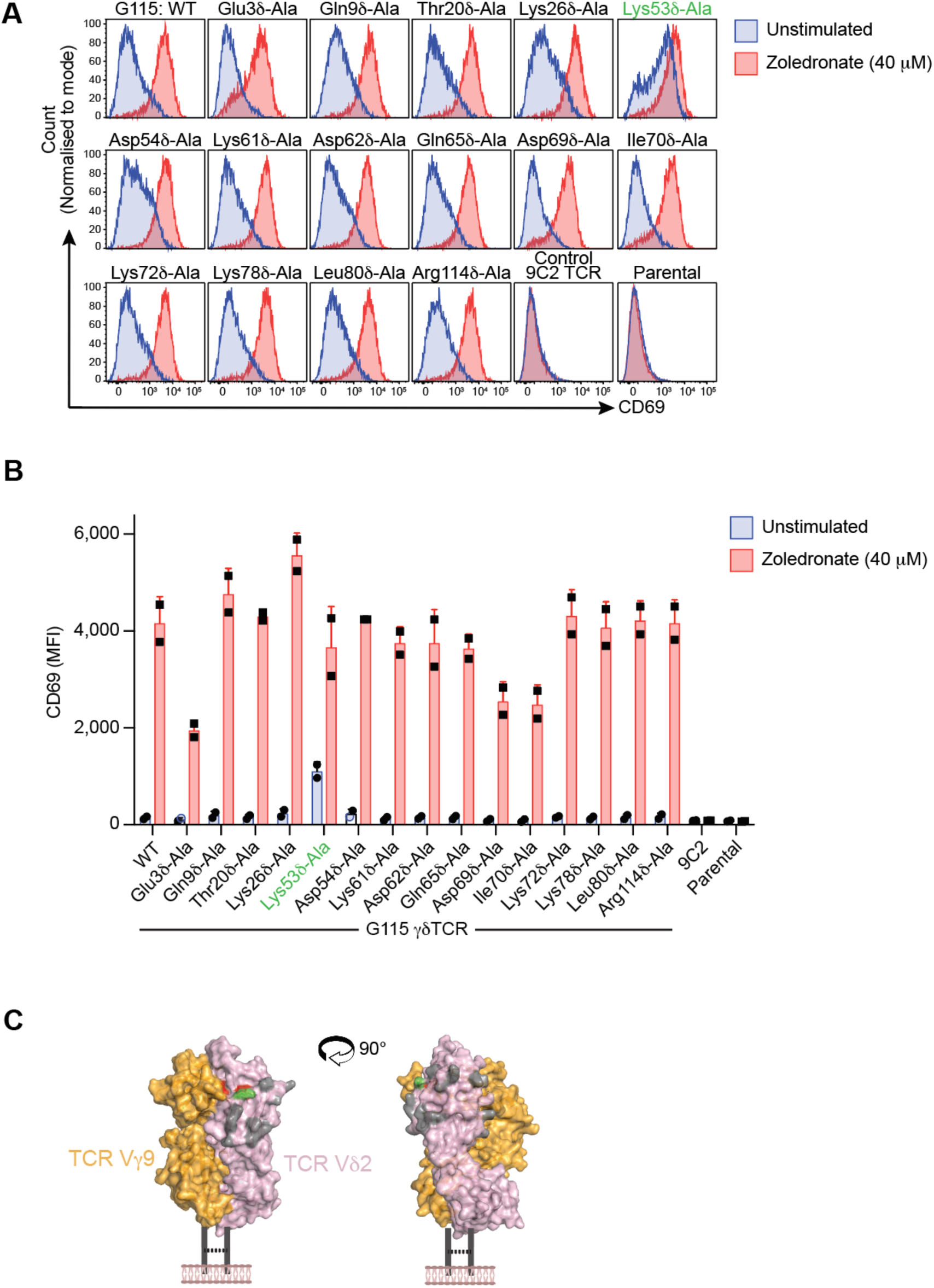
(**A**) and (**B**) CD69-PE expression on G115 WT or mutant Vγ9Vδ2^+^, 9C2 Vγ5Vδ1^+^ γδTCR or parental (TCR^-^) J.RT3-T3.5 Jurkat cells after overnight co-culture with LM-MEL-75 APCs in the presence (red) or absence (blue) of 40 μM zoledronate. Graphs are presented as mean ± SEM. N = 2, where each point represents an independent experiment. (**C**) Surface representation of G115 Vγ9Vδ2^+^ TCR (γ-chain, orange; δ-chain, light pink) depicting the interacting (red) and gatekeeper (green) or uninvolved (gray) residues based on Jurkat activation assays.

**Extended Data Fig. 4.**
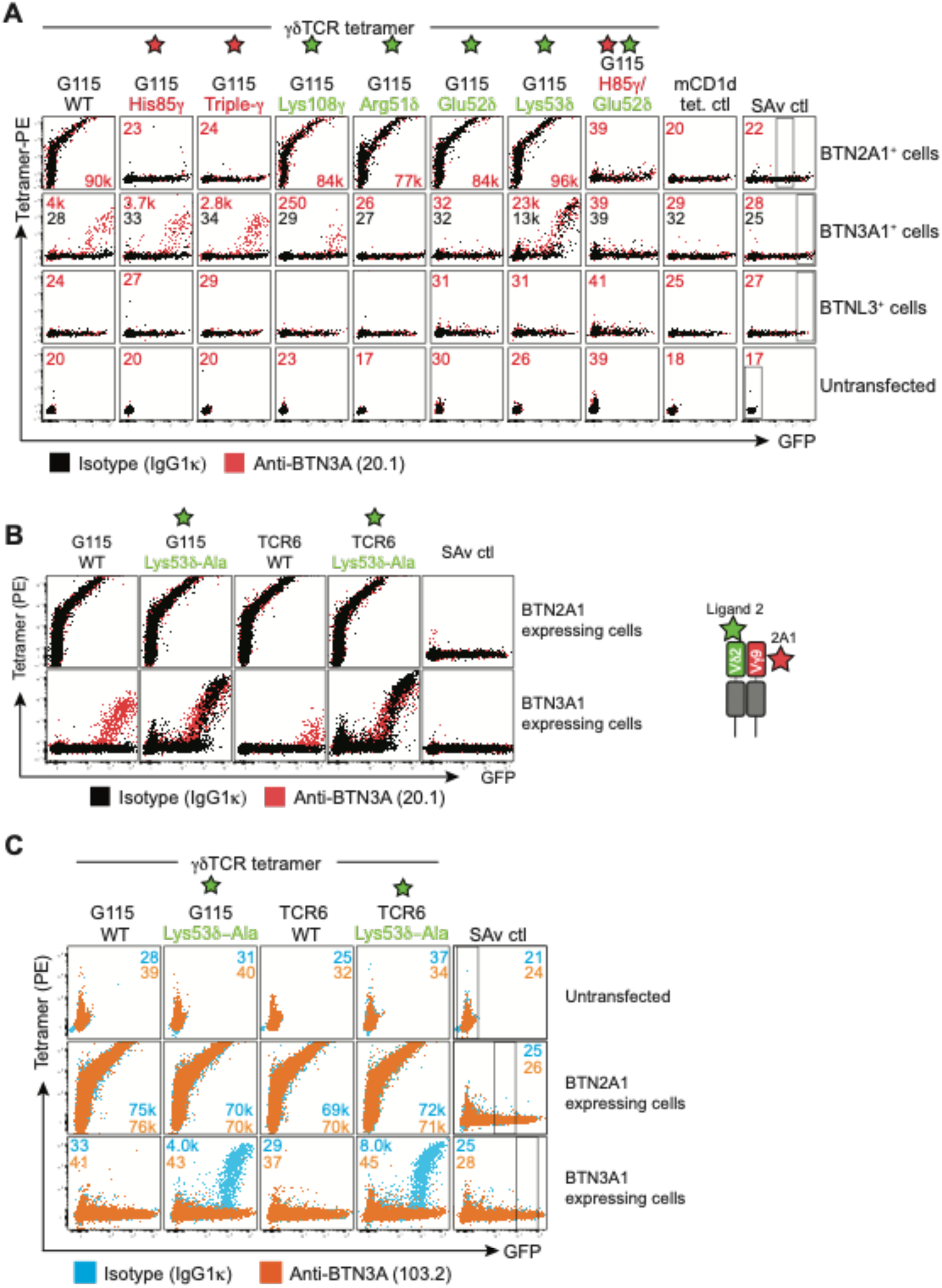
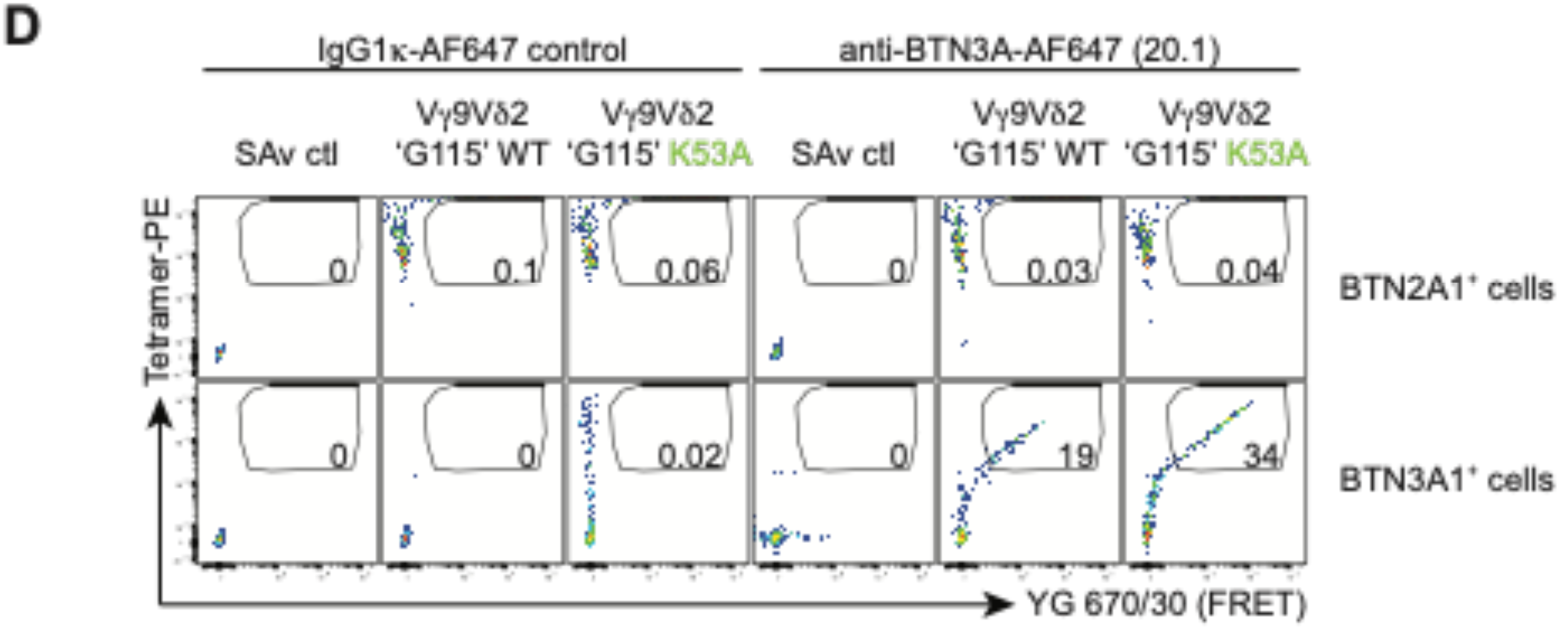
MFI of G115 γδTCR tetramer staining of (**A**) Wild-type or mutant G115 Vγ9Vδ2^+^ TCR tetramer staining, or control mouse CD1d-α-GalCer (mCD1d tet.) or streptavidin alone (SAv) staining of NIH-3T3 cells transfected with BTN2A1, BTN3A1 or BTNL3 ± anti-BTN3A mAb clone 20.1 (red) or isotype control (IgG1,κ, black). Triple-γ mutant comprises Arg20γ-Ala/Glu70γ- Ala/His85γ-Ala mutations. Cartoon inset depicts the locations of BTN2A1-epitope (red star) and the ligand-two epitope (green star). Representative of one of three independent experiments. MFI of PE for mAb 20.1-treated cells (red numbers) or isotype control (IgG1,κ)-treated BTN3A1^+^ cells (black numbers) shown within the depicted GFP^+^ gate. (**B**) GFP^+^ BTN2A1-transfected or GFP^+^ BTN3A1-transfected NIH-3T3 cells were stained with streptavidin (SAv)-PE control, Vγ9Vδ2^+^ ‘G115 WT’, ‘G115 Lys53δ-Ala’, ‘TCR 6 WT’ or ‘TCR 6 Lys53δ-Ala’ TCR tetramers. Representative one of two independent experiments. (**C**) Mouse 3T3 fibroblasts expressing BTN2A1 or BTN3A1 were treated with anti-BTN3A antagonist mAb (103.2; orange) or an isotype control (MOPC-21; blue) and subsequently stained with control SAv-PE or Vγ9Vδ2 TCR tetramer-PE (wild-type or Lys53-Ala mutants of clones G115 and TCR6). Median fluorescence intensity (MFI) of PE for mAb 103.2-treated cells (orange numbers) or isotype control (IgG1,κ)-treated cells (blue numbers) shown within the depicted GFP^+^ gate. Representative one of three independent experiments. (**D**) GFP^+^ BTN2A1-transfected or GFP^+^ BTN3A1-transfected NIH-3T3 cells were stained with isotype control (MOPC21)-AF647 or anti-BTN3A (20.1)-AF647 antibodies followed by control SAv-PE, Vγ9Vδ2 ‘G115 WT’ or ‘G115 Lys53δ-Ala’ TCR tetramer-PE staining. Cells were examined for FRET in the YG 670/30 channel by flow cytometry.

**Extended Data Fig. 5.**
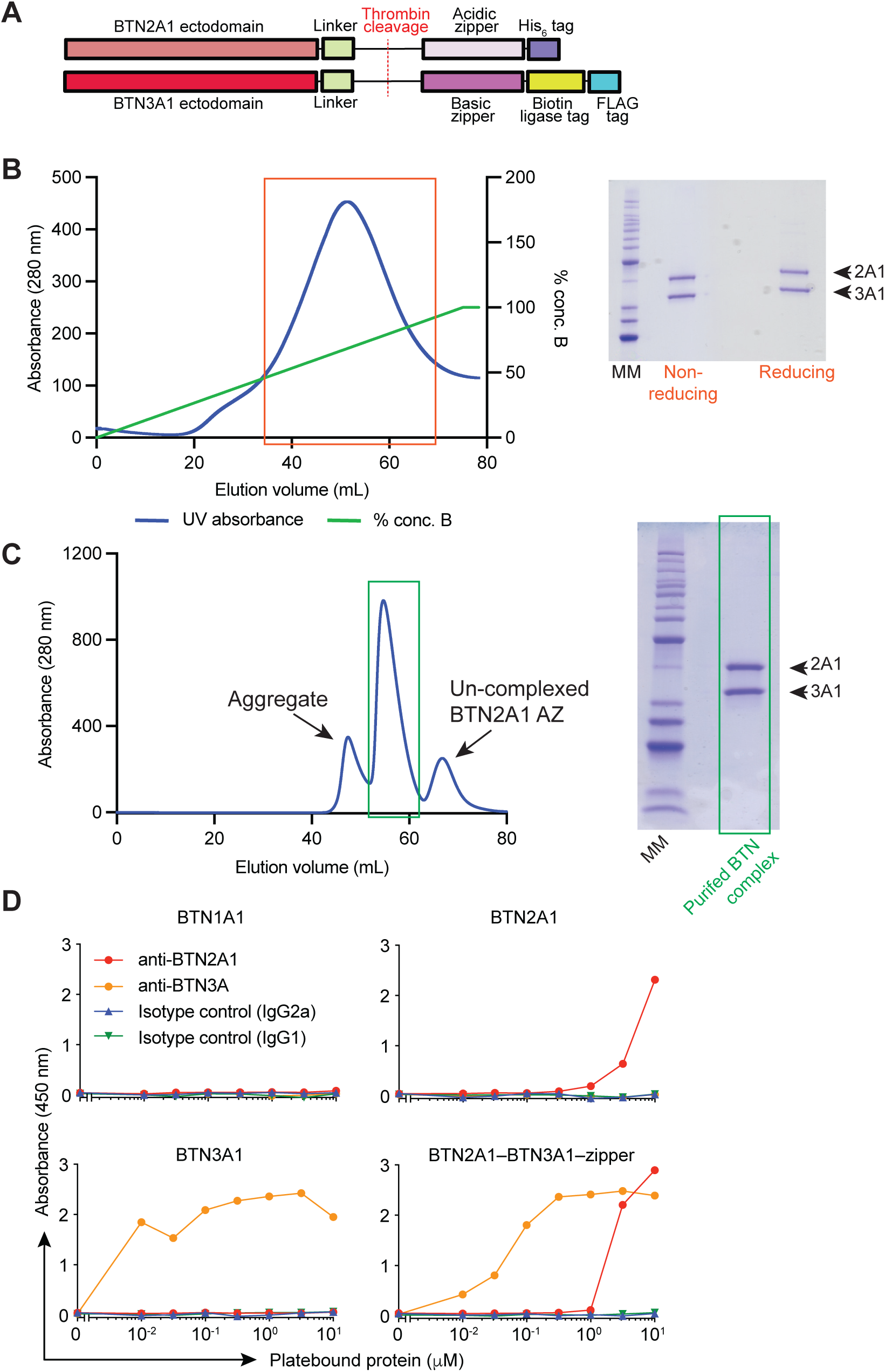
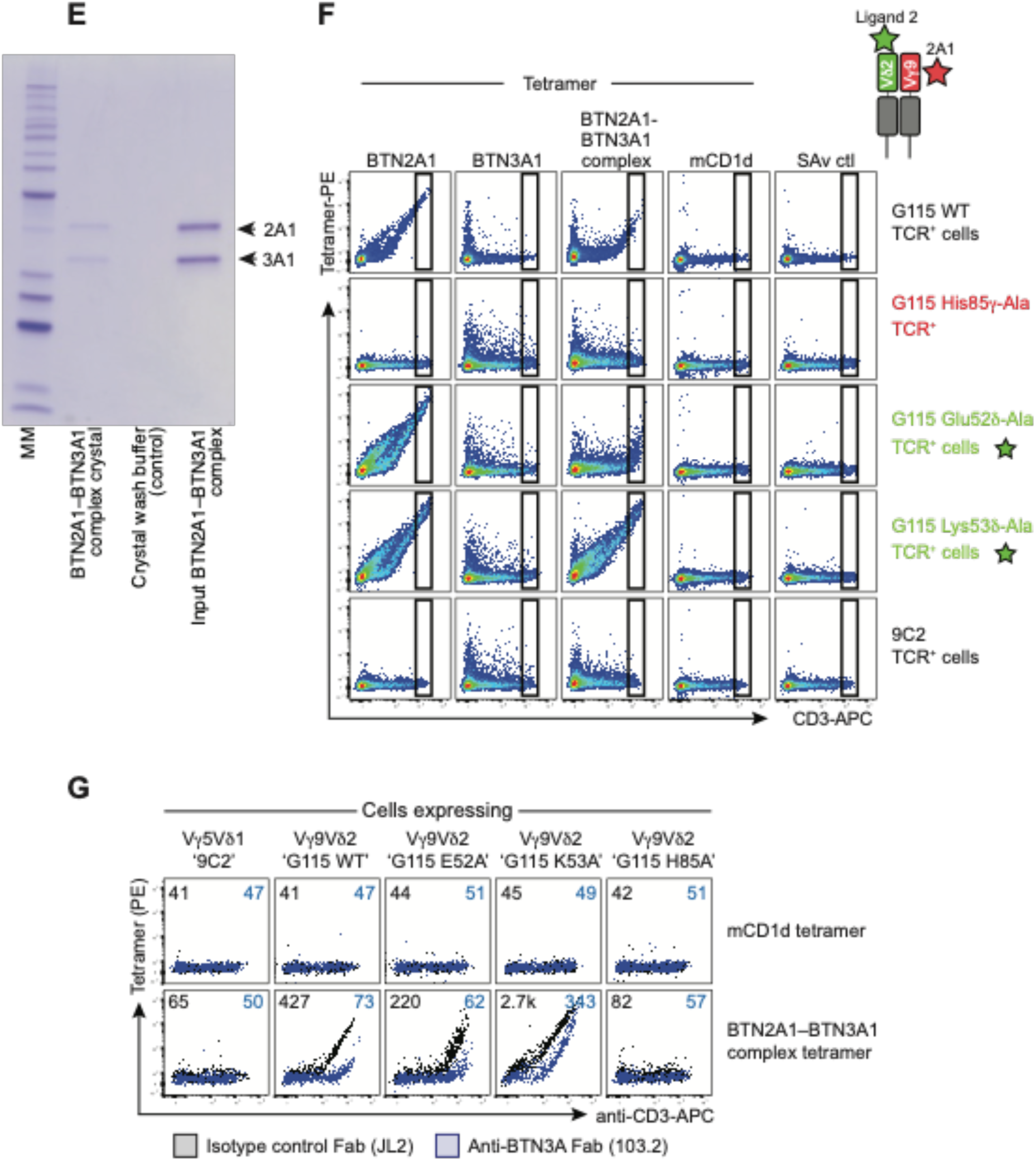
(**A**) Representation of BTN2A1–BTN3A1–zipper complex. (**B-C**) (**LHS**) BTN2A1–BTN3A1 complex was expressed in Expi293F cells and purified by (**B**) affinity (NiNTA) and (**C**) size exclusion (S200) chromatography. (**RHS**) Protein purified in boxes run over SDS-PAGE to confirm identity. MM – molecular weight marker; 2A1 – BTN2A1–acid zipper (AZ)–His_6_; 3A1 – BTN3A1–basic zipper (BZ)–Biotin ligase tag. (**D**) Reactivity of anti-BTN2A1 (clone 259, red), anti-BTN3A (clone 103.2, orange), mouse IgG2a isotype control (clone BM4-2a, blue), or mouse IgG1 isotype control (clone MOPC-21, green) to immobilized BTN1A1, BTN2A1, BTN3A1 or BTN2A1– BTN3A1–zipper ectodomains by ELISA. (**E**) BTN2A1–BTN3A1–zipper complex was crystallized, resolubilized and run on SDS-PAGE, along with crystal wash buffer and input BTN2A1–BTN3A1– zipper complex. MM – molecular weight marker; 2A1 – BTN2A1–AZ–His_6_; 3A1 – BTN3A1–BZ–Biotin ligase tag. (**F**) BTN2A1-, BTN3A1-, BTN2A1–BTN3A1 complex- or control mouse CD1d-ectodomain tetramers, or streptavidin alone (SAv) versus anti-CD3 staining of HEK293T cells co-transfected with CD3 plus G115 Vγ9Vδ2^+^ TCR wild-type, His85γ-Ala, Glu52δ-Ala, Lys53δ-Ala or control 9C2 Vγ5Vδ1^+^ TCR. Cartoon inset depicts the relative locations of BTN2A1-epitope mutants (red star) or ligand-two epitope mutants (green star). Representative of one of three independent experiments. (**G**) Soluble BTN2A1–BTN3A1 ectodomain complex tetramers were pre-incubated with anti-BTN3A Fab (clone 103.2; blue) or an isotype control Fab (JL2; black) prior to staining 293T cells expressing Vγ5Vδ1 ‘9C2’ control TCR or Vγ9Vδ2 ‘G115’ mutant TCRs. Representative one of two independent experiments. Black text – MFI of isotype control (JL2) Fab treated tetramers; blue text – MFI of anti-BTN3A (103.2) Fab treated tetramers.

**Extended Data Fig. 6.**
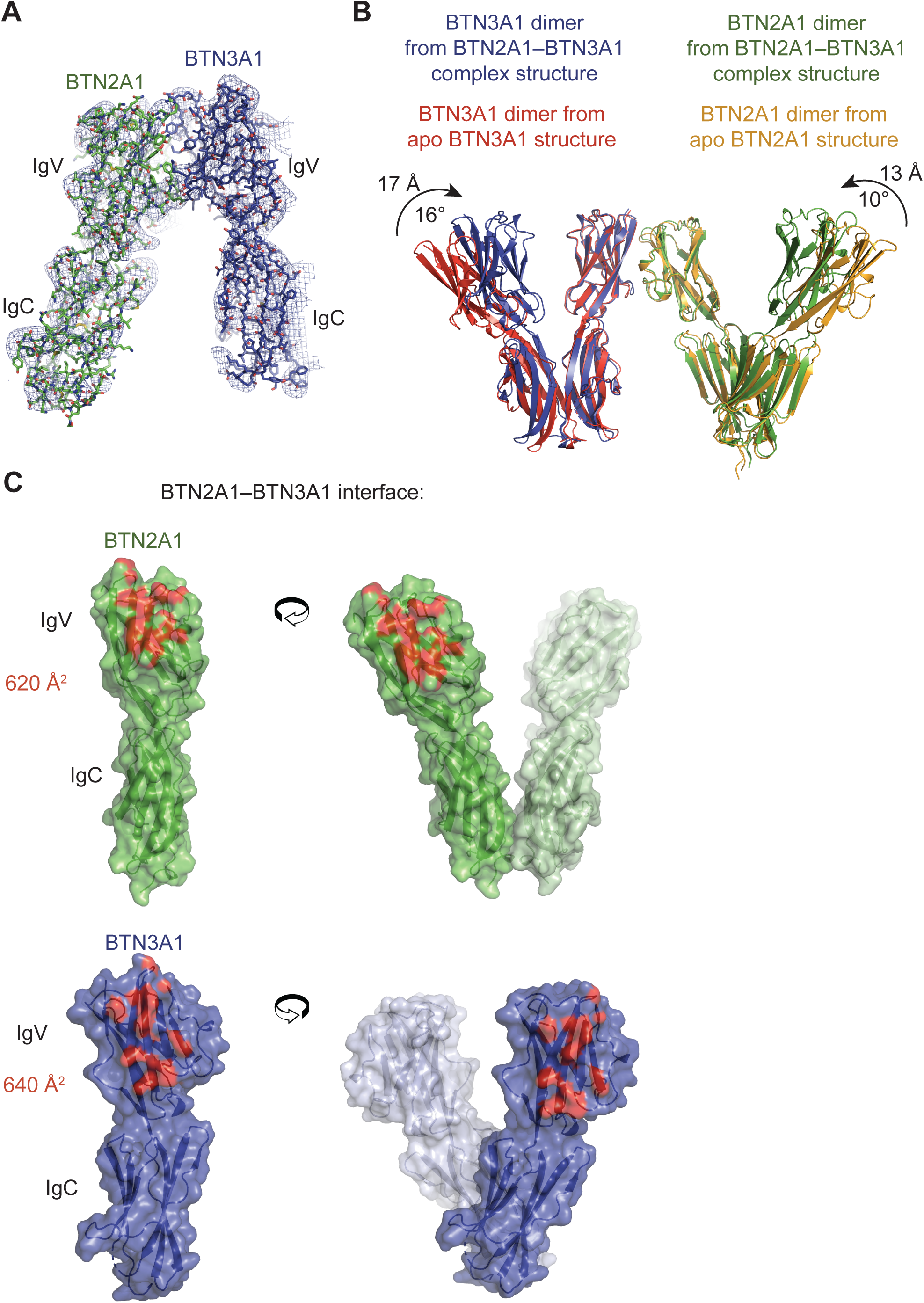
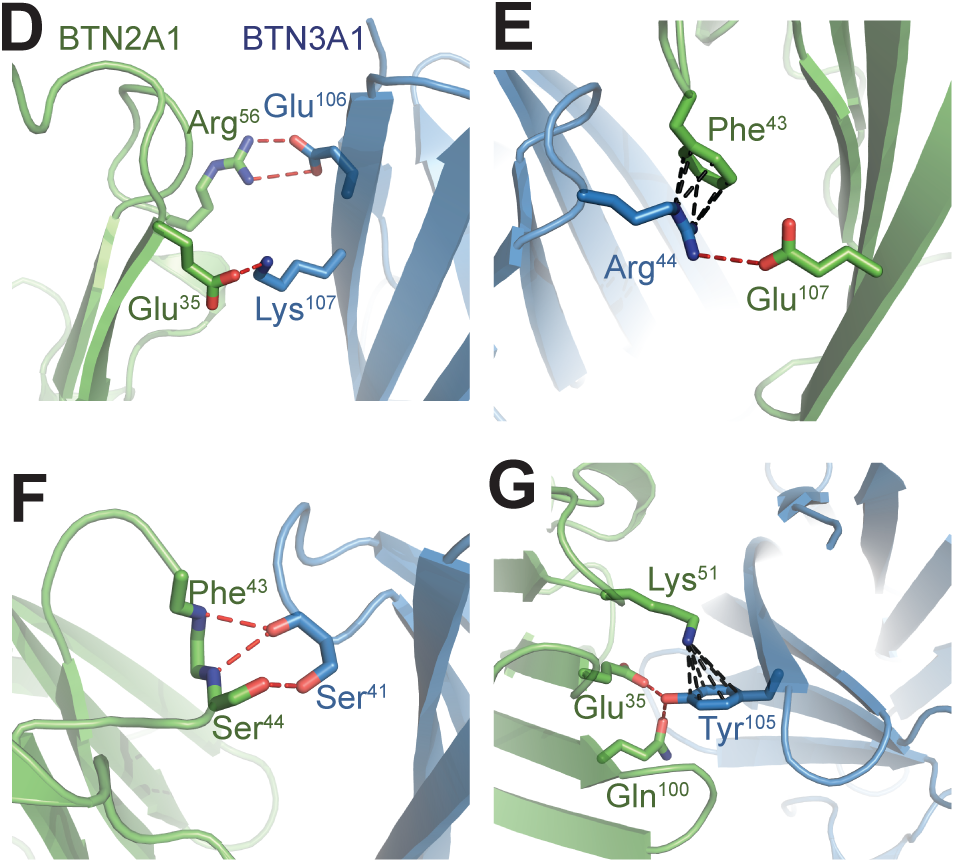
(**A**) 2mFo-DFc electron density composite omit map contoured at 1σ, blue mesh, surrounding BTN2A1 and BTN3A1 molecules. (**B**) Comparison of the apo BTN3A1 homodimer (PDB code 4F80, red cartoon) with BTN3A1 homodimer from the BTN2A1–BTN3A1–zipper complex (PDB code 8DFX, blue cartoon), and a comparison of apo BTN2A1 homodimer (PDB code 8DFY, orange cartoon) with BTN2A1 homodimer from the BTN2A1–BTN3A1-zipper complex (green cartoon). (**C**) Surface representation of BTN2A1 (green) and BTN3A1 (blue) depicting the regions that are contacting each other (red). Molecular contacts between BTN2A1 (green) and BTN3A1 (blue) ecdodomains showing the (**D**) BTN2A1 Arg56 and Glu35, (**E**) Phe43 and Glu107, (**F**) Phe43 N atom and Ser44, and (**G**) Glu35, Lys51 and Gln100 side and/or main chains and their BTN3A1 contacts as sticks. H-bonds and salt-bridges, red; cation-π, black.

**Extended Data Fig. 7.**
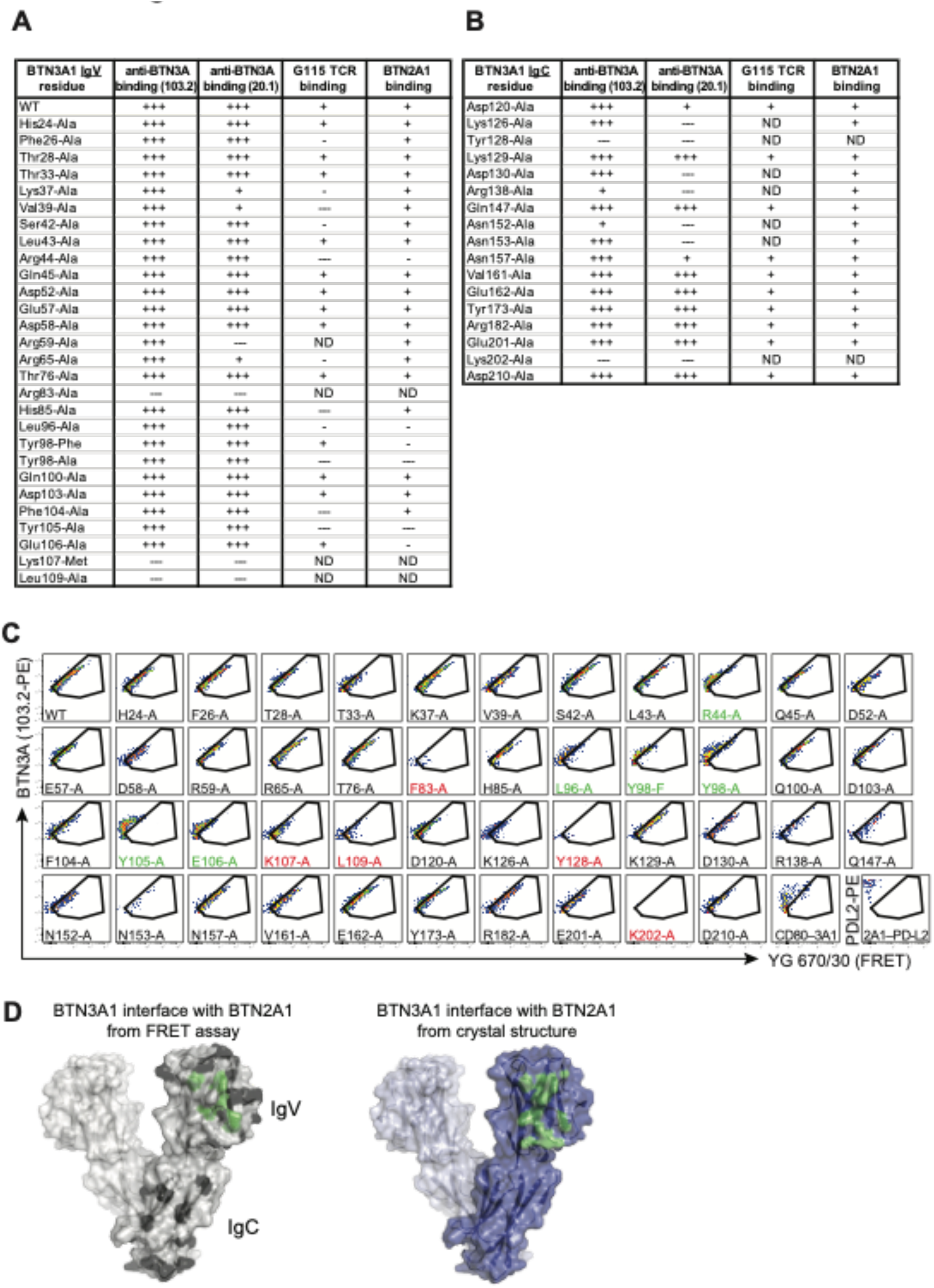
Summary of the effect of single-residue mutations within the (**A**) IgV domain or (**B**) IgC domain of BTN3A1 on anti-BTN3A reactivity (mAb clones 103.2 and 20.1) as well as binding in cis to BTN2A1 as measured by FRET, and binding to G115 γδTCR tetramer. +++ high expression; + intermediate expression; - low expression; --- no expression; ND – no binding data due to lack of BTN3A1 expression or lack of 20.1 mAb binding. (**C**) Förster resonance energy transfer (FRET) between anti-BTN2A1 (clone 259) and anti-BTN3A (clone 103.2) mAb staining on gated BTN2A1^+^BTN3A1^+^ NIH-3T3 cells, 48 h after co-transfection with WT BTN2A1 plus the indicated BTN3A1 mutant, or as irrelevant controls, BTN2A1 plus PD-L2 or BTN3A1 plus CD80. Mutants in red were excluded from analysis due to diminished BTN3A1 staining. Mutants in green are those which reduced FRET levels. Representative one of six independent experiments. (**D**) Surface representation of BTN3A1 V-dimer depicting residue side chains that upon mutation led to an abrogation of BTN3A1 association with BTN2A1 (green), or those which did not impact the interaction with BTN2A1 (black), as determined by the FRET assay (left). The BTN3A1 surface on the right depicts atoms that contacted BTN2A1 based on the crystal structure (green, reproduced from Extended Data Fig. 5B).

**Extended Data Fig. 8.**
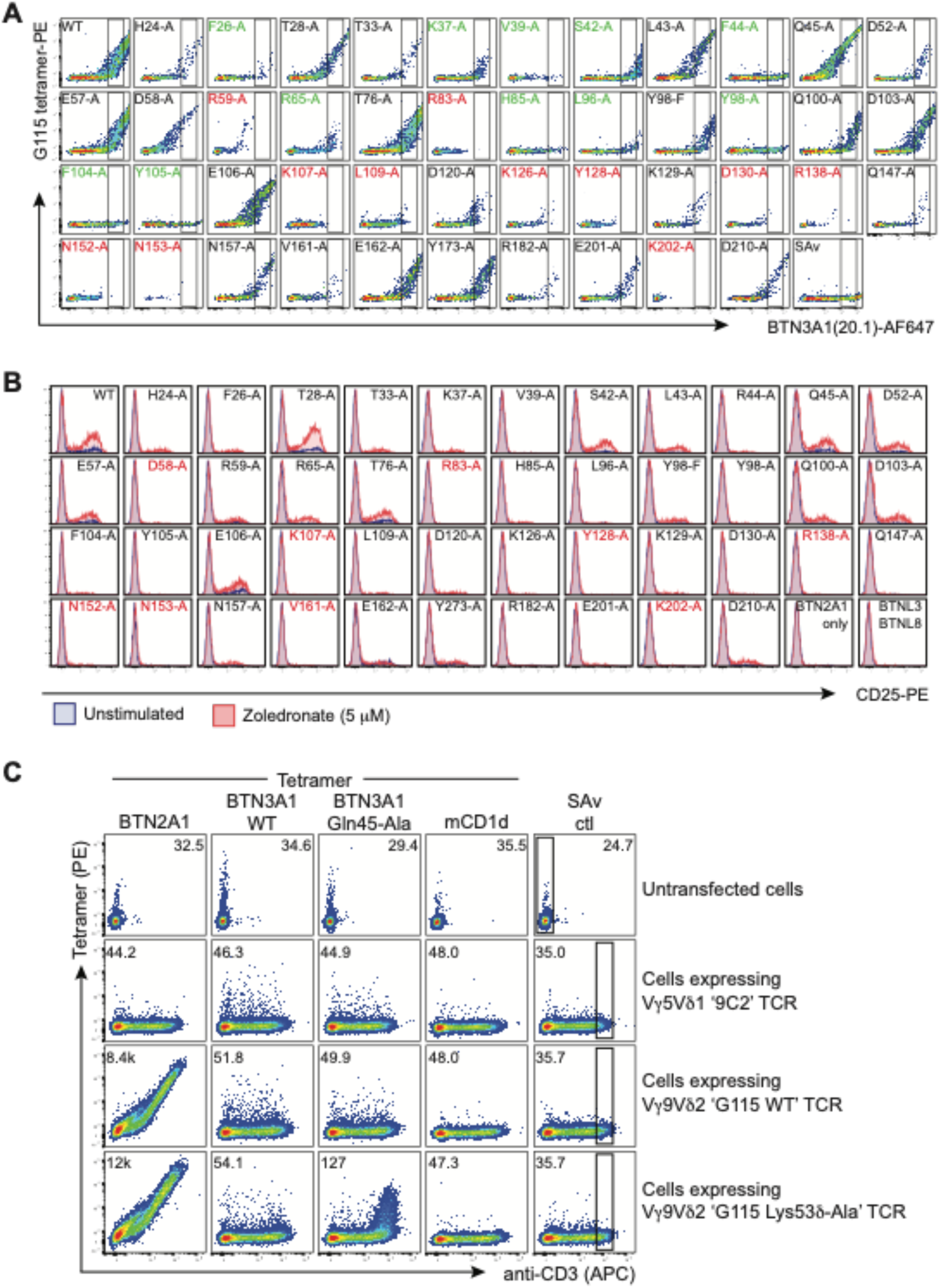
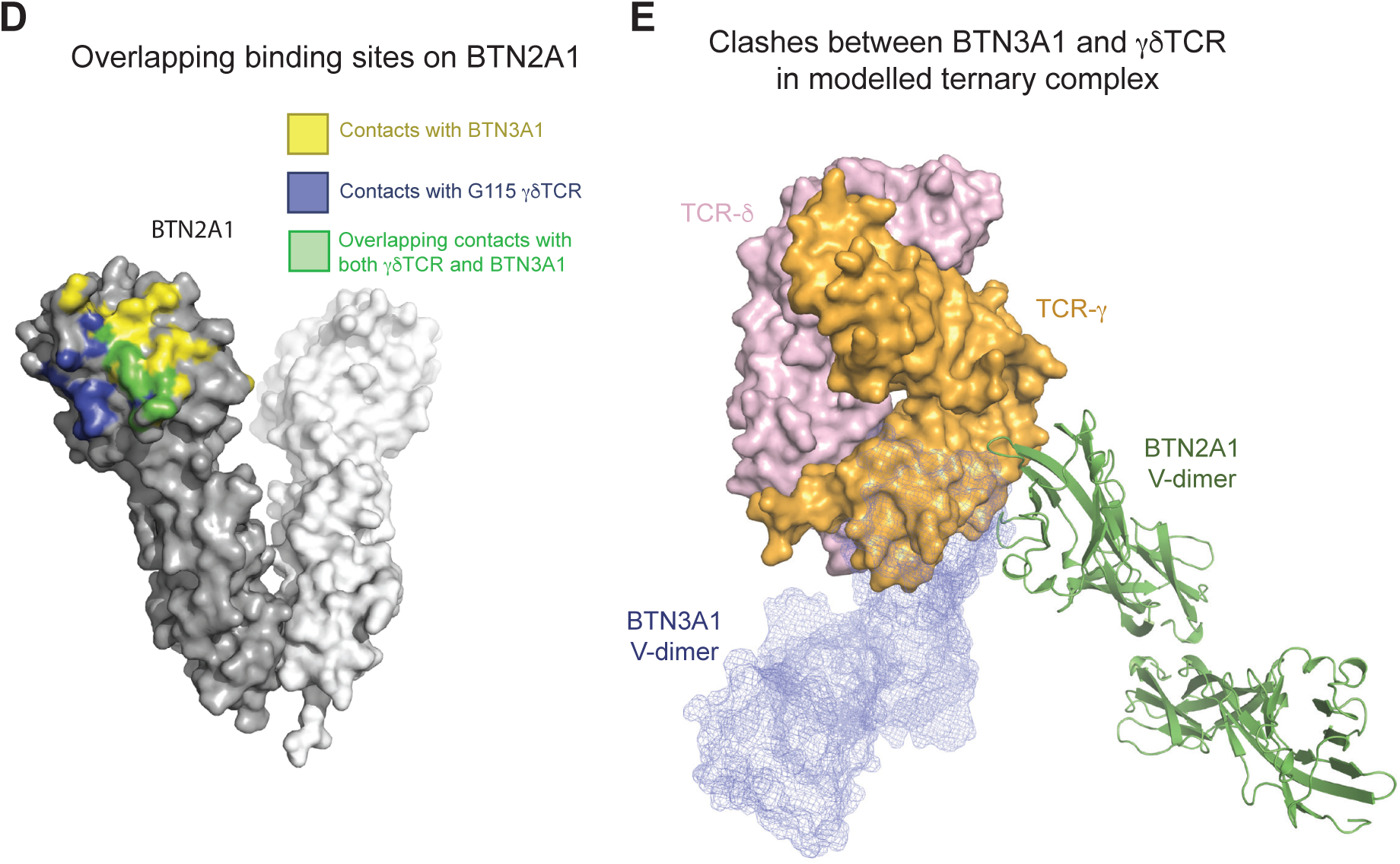
(**A**) G115 tetramer-PE staining of BTN3A1 WT or mutant-transfected NIH-3T3 cells following pre-incubation with anti-BTN3A-AF647 (mAb clone 20.1). Mutants in red were excluded from analysis due to diminished BTN3A1 mAb 20.1 staining. Mutants in green are those which impaired G115 tetramer staining. Representative of one of three independent experiments. (**B**) CD25-PE expression on purified pre-expanded Vδ2^+^ γδ T cells following co-culture with NIH-3T3 cells that were co-transfected with BTN2A1 plus the indicated BTN3A1 mutant, or alternatively control BTNL3 plus BTNL8, ± zoledronate (5 µM) for 24 h. Data are from one of three independent experiments, each with two donors. (**C**) BTN2A1, BTN3A1^WT^, BTN3A1^Gln45-Ala^ or mCD1d tetramer-PE, or control SAv-PE staining of Vγ9Vδ2 TCR-transfected HEK293T *BTN2A^KO^.BTN3A^KO^* cells. MFI of PE for TCR^+^ cells (black numbers) shown within the depicted CD3^+^ gate. Representative one of three independent experiments. (**D**) Surface of BTN2A1 V-dimer depicting residues that contact BTN3A1 based on the BTN2A1–BTN3A1 crystal structure in yellow, residues that contact Vγ9Vδ2^+^ TCR based on the G115 TCR–BTN2A1 crystal structure in blue, and residues that overlap and contact both in red. (**E**) Structure of BTN2A1 (green ribbon)–BTN3A1 (blue mesh) and BTN2A1–G115 Vγ9Vδ2^+^ TCR (surface render) showing clashes between BTN3A1 and TCRγ interacting with BTN2A1.

**Extended Data Fig. 9.**
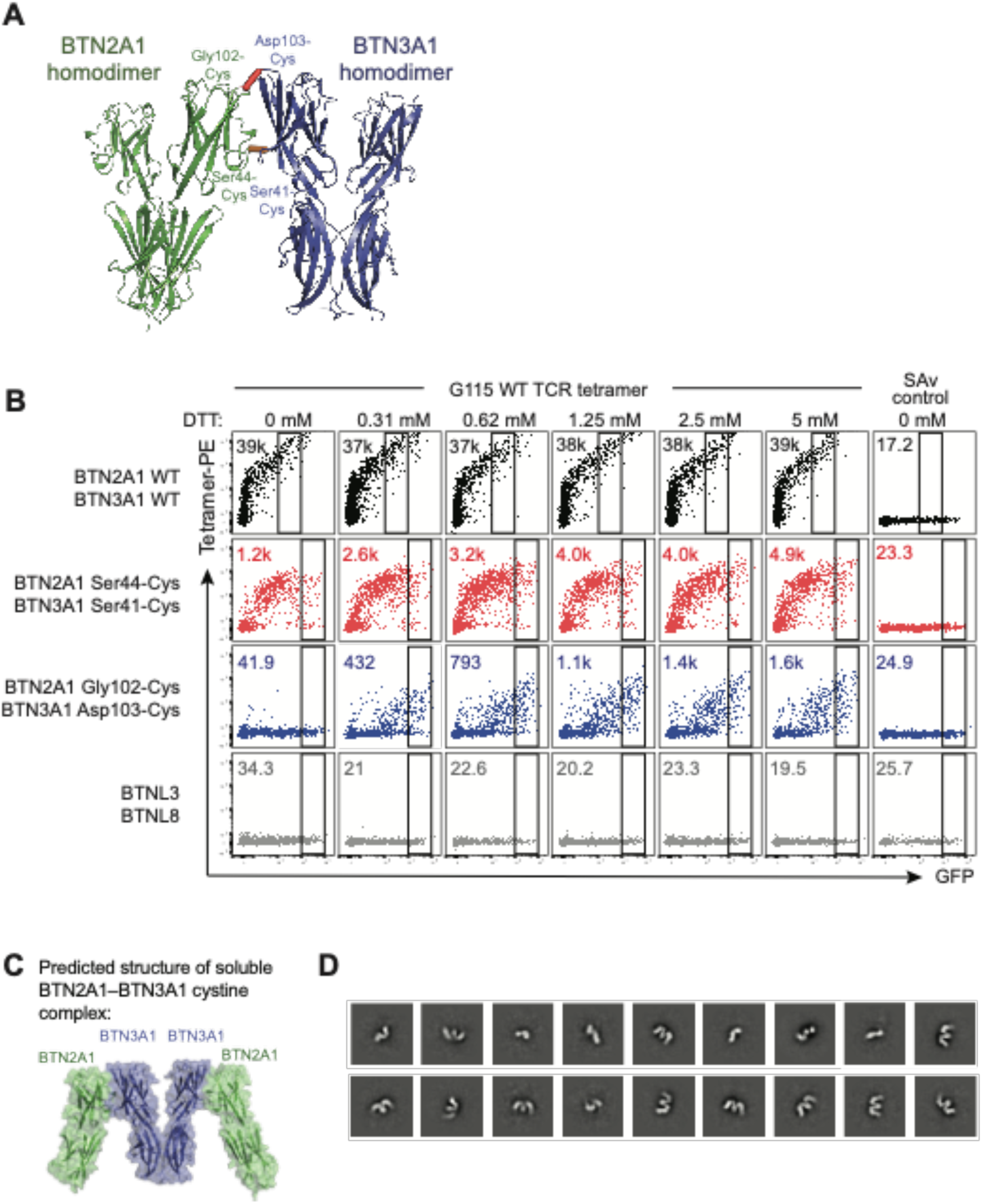
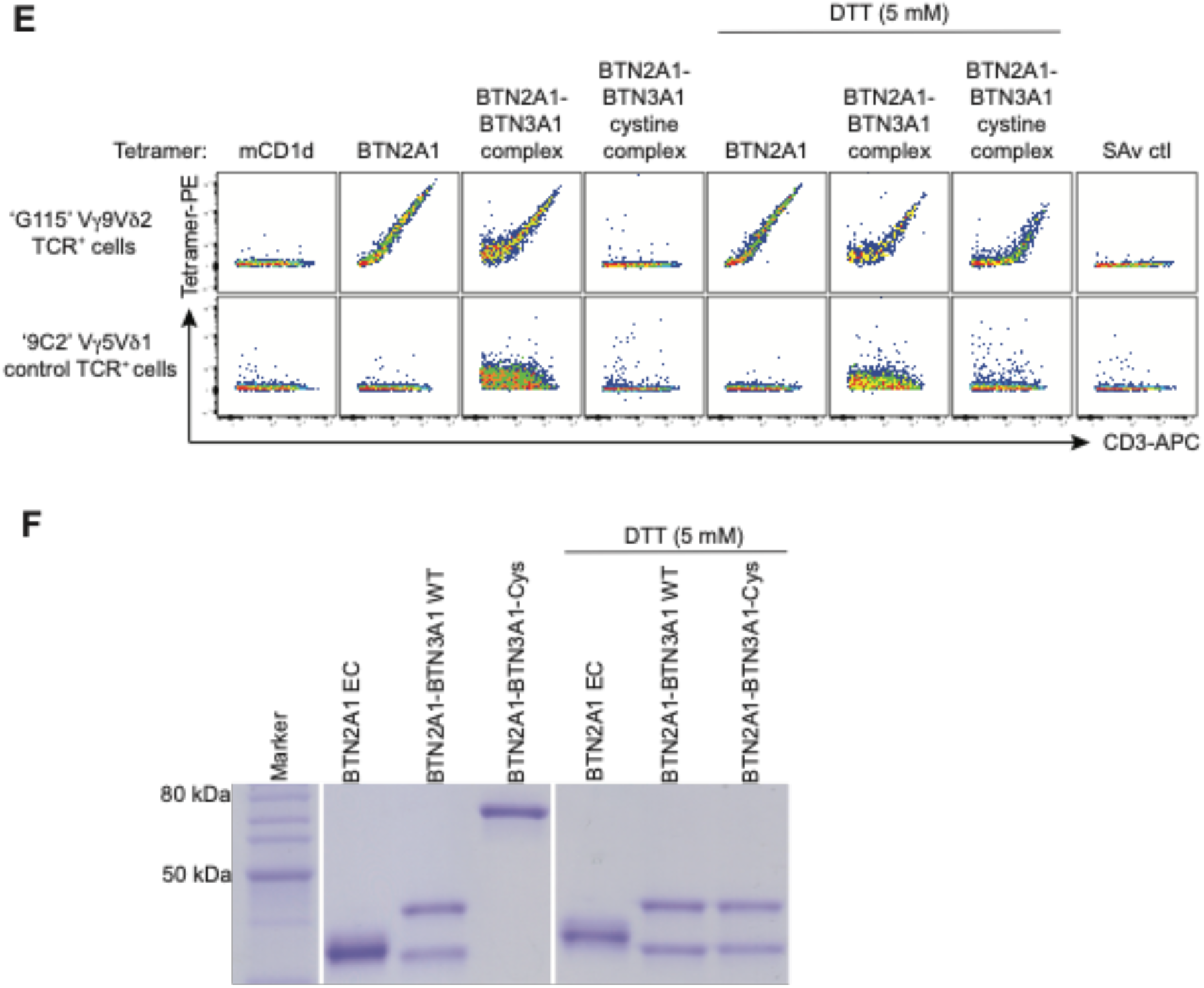
(**A**) Structure of BTN2A1–BTN3A1 depicting the locations of the two cysteine mutant pairs. (**B**) G115 Vγ9Vδ2+ TCR tetramer-PE, or control streptavidin-PE (SAv) staining of mouse NIH-3T3 fibroblasts co-transfected with the indicated BTN2A1 and BTN3A1 cysteine mutant pairs, or control BTNL3 plus BTNL8, following pre-treatment of cells with graded concentrations of dithiothreitol (DTT). Inset – MFI of PE parameter. Representative of one of four independent experiments. (**C**) Predicted structure of the BTN2A1–BTN3A1 complex containing a disulphide bond between BTN2A1 and BTN3A1 molecules, based on the BTN2A1–BTN3A1–zipper ectodomain crystal structure. (**D**) 2D class averages of negatively stained soluble BTN2A1 Gly102-Cys–BTN3A1 Asp103-Cys ectodomain complex. (**E**) BTN2A1 tetramer, BTN2A1–BTN3A1 complex tetramer, BTN2A1 Gly102-Cys– BTN3A1 Asp103-Cys complex tetramer, or control tetramer (mouse CD1d) or SAv.-PE alone staining of HEK293T cells transfected with either G115 Vγ9Vδ2^+^ or control 9C2 Vγ5Vδ1^+^ γδTCRs. Where indicated, BTN molecules were pre-treated with 5 mM DTT prior to being tetramerised with SAv-PE. Representative of one or four independent experiments. (**F**) SDS-PAGE of BTN monomers treated with DTT as in (E). EC, ectodomain.

**Extended Data Fig. 10.**
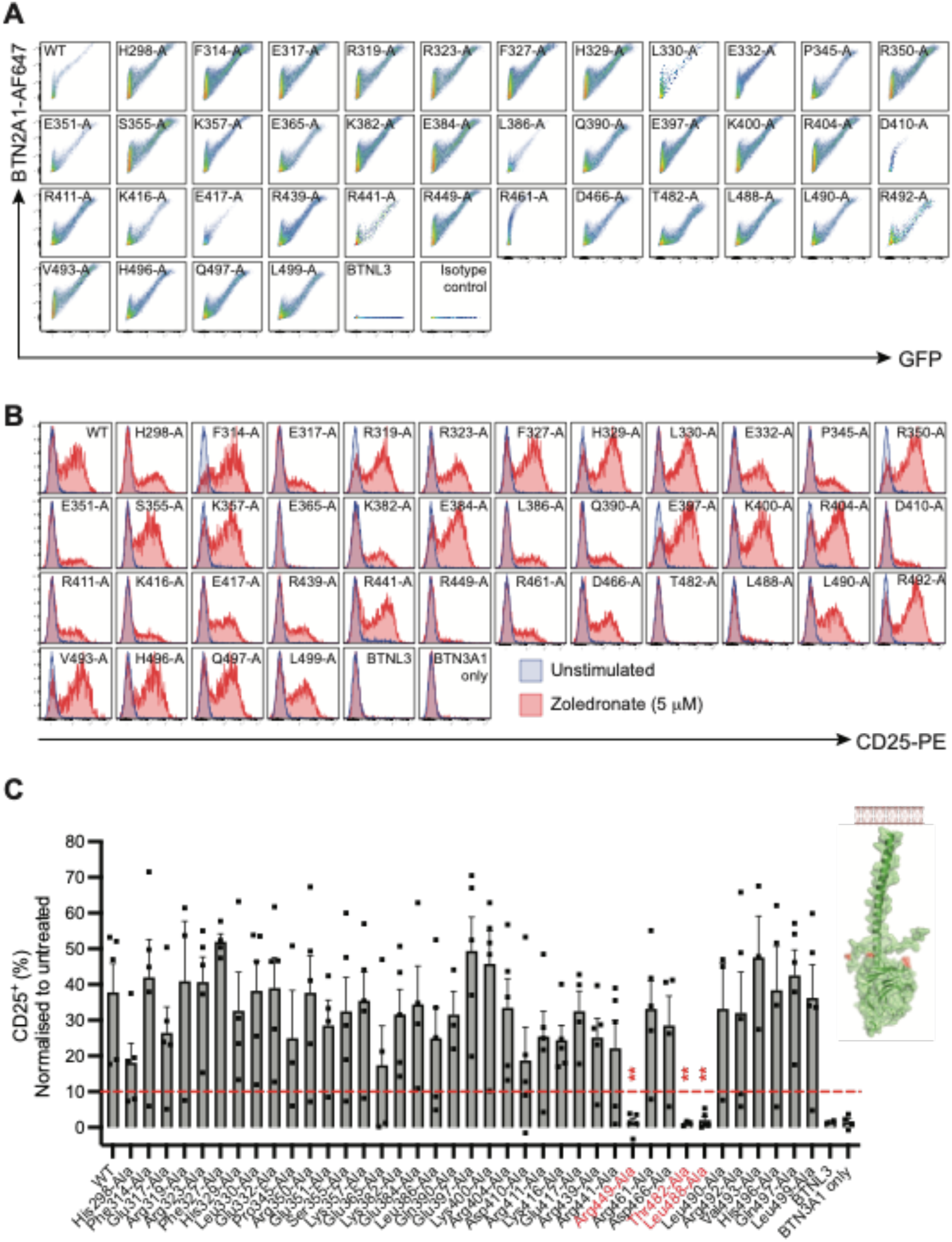
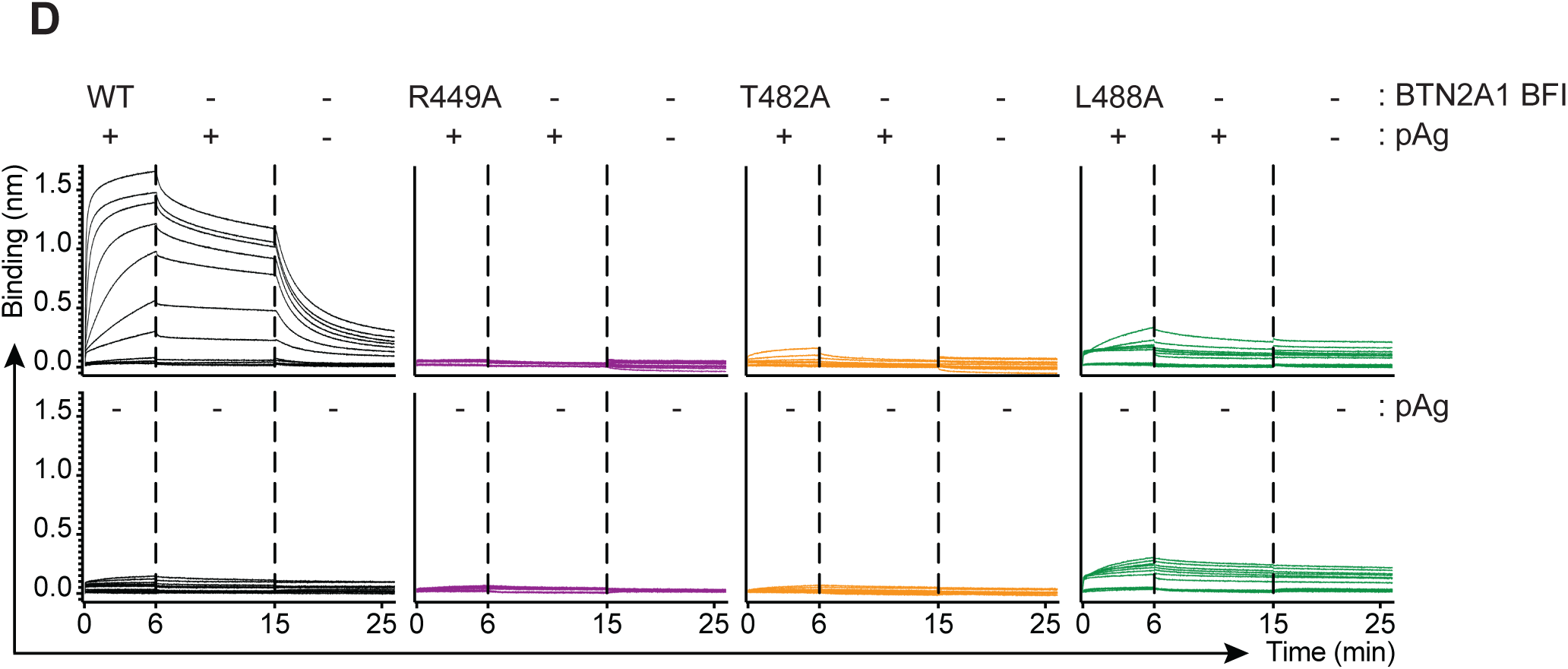
Zoledronate augments the BTN3A1-dependent interaction with Vγ9Vδ2 TCR. (**A**) Surface BTN2A1 expression (clone 259) on HEK293T *BTN2A*^KO^.*BTN3A*^KO^ cells that were transfected with BTN2A1 WT or the indicated BTN2A1 intracellular domain mutants, or control BTNL3. Representative from one of two experiments. (**B**) Representative plots and (**C**) mean ± SEM of CD25-PE expression on purified pre-expanded Vδ2^+^ γδ T cells following co-culture with HEK293T *BTN2A*^KO^.*BTN3A*^KO^ cells that were co-transfected with BTN3A1 plus the indicated BTN2A1 mutant, or alternatively, control BTNL3 alone or BTN3A1 alone, ± zoledronate (5 µM) for 24 h. Replicates with low transfection efficiency (< 10% GFP^+^) were excluded from analysis. Data are from three independent experiments each with n=2 different donors. ** P < 0.01 by two-way ANOVA with Šidák multiple comparison correction. Insert, molecular model of the BTN2A1 B30.2 intracellular domain generated by AlphaFold v2 with functionally important residues shown in red. (**D**) The B30.2 domain of wild-type (WT) BTN3A1 was used as ligand to probe the interaction with WT or mutant BTN2A1 full length intracellular domain (BFI), in the absence or presence of pAg (20 µM HMBPP) by biolayer interferometry (BLI).

**Extended Data Fig. 11.**
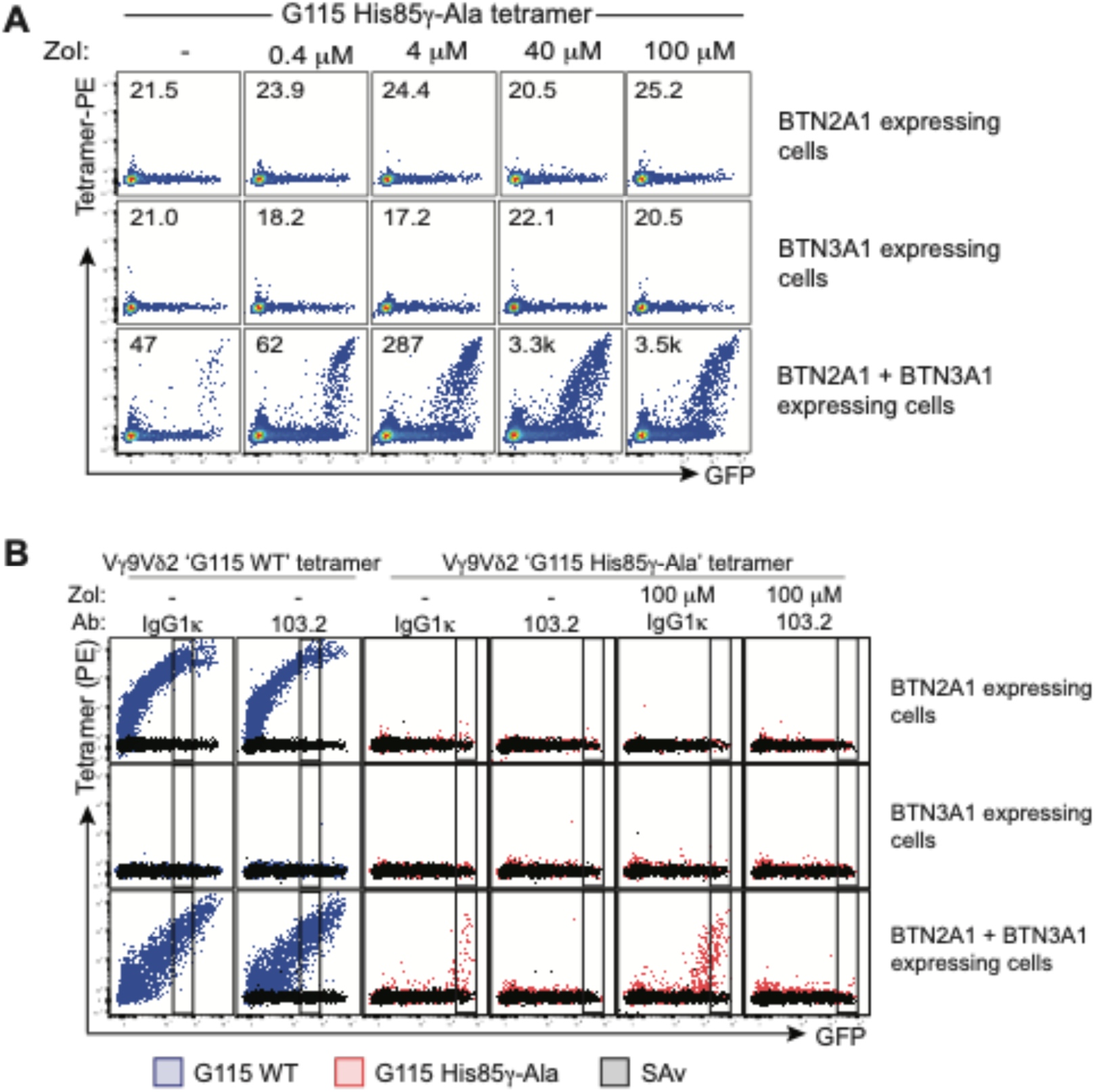
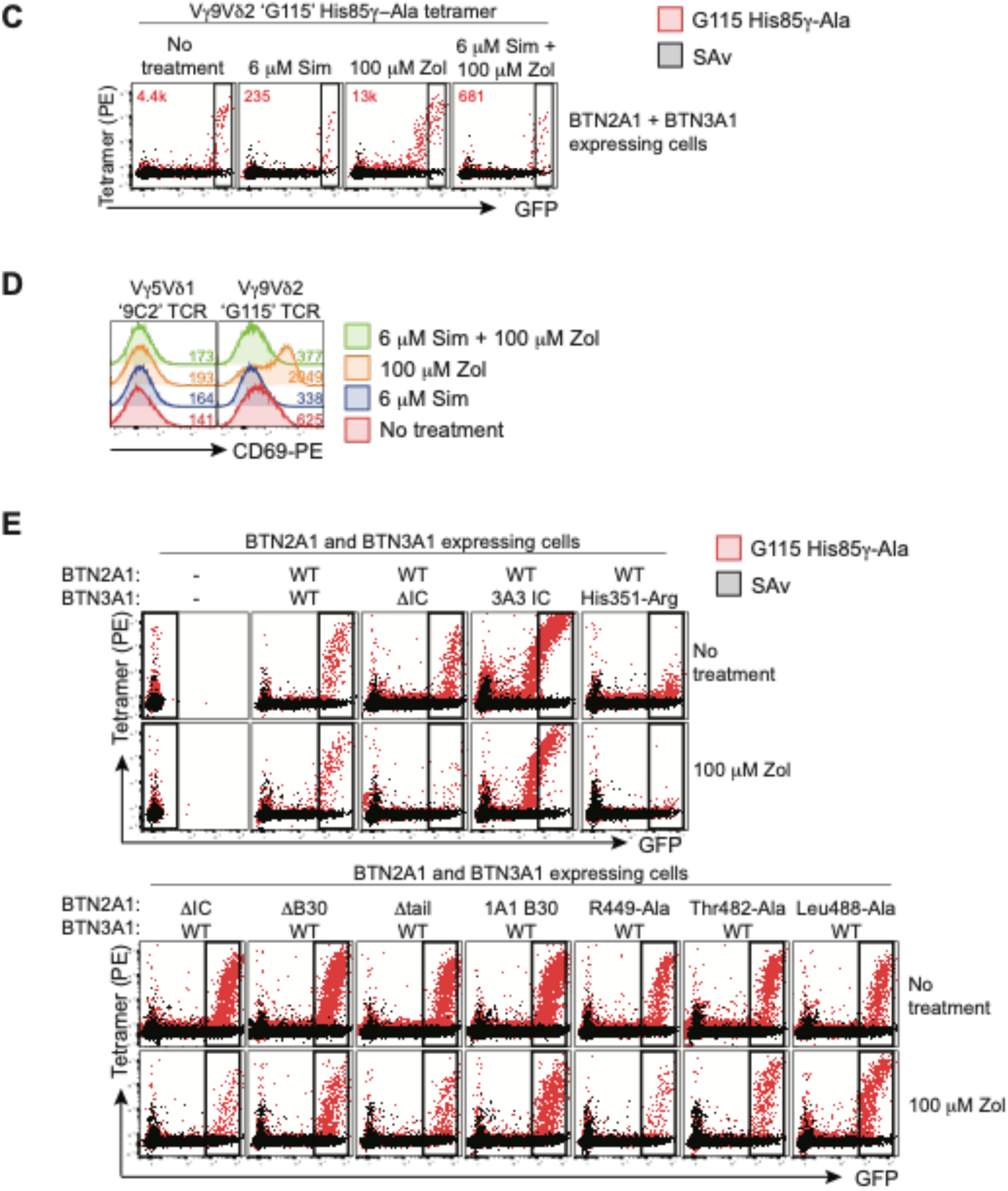
(**A**) Mouse NIH-3T3 cells transfected with BTN2A1, BTN3A1 or both BTN2A1 and BTN3A1 were treated with various concentrations of Zol and stained with Vγ9Vδ2^+^ ‘G115 His85γ- Ala’ TCR tetramer, after first gating on GFP^+^ cells. Representative of one of four (BTN3A1 expressing cells and BTN2A1 plus BTN3A1 co-expressing cells) or two (BTN2A1 expressing cells) independent experiments. (**B**) NIH-3T3 fibroblasts transfected with BTN2A1, BTN3A1 or both BTN2A1 and BTN3A1, were untreated or treated with 100 µM Zoledronate (Zol), and/or isotype (IgG1k) or anti-BTN3A (clone 103.2). Cells were then probed by Vγ9Vδ2^+^ ‘G115 His85γ-Ala’ TCR tetramer or control streptavidin-PE (SAv) staining. (**C**) 3T3 cells were transfected with BTN2A1 and BTN3A1 and treated with 100 µM Zol or 6 µM Simvastatin (Sim). (**D**) CD69-PE expression on G115 Vγ9Vδ2^+^, 9C2 Vγ5Vδ1^+^ γδTCR J.RT3-T3.5 Jurkat cells after co-culture with LM-MEL-62 APCs treated with 100 µM Zol or 6 µM Sim for 24hr. (**E)** 3T3 cells were transfected with wild-type BTN2A1 plus BTN3A1 molecules engineered with impaired pAg-reactivity (upper); or wild-type BTN3A1 plus BTN2A1 molecules engineered to lose pAg-reactivity (lower).

**Extended Data Fig. 12.**
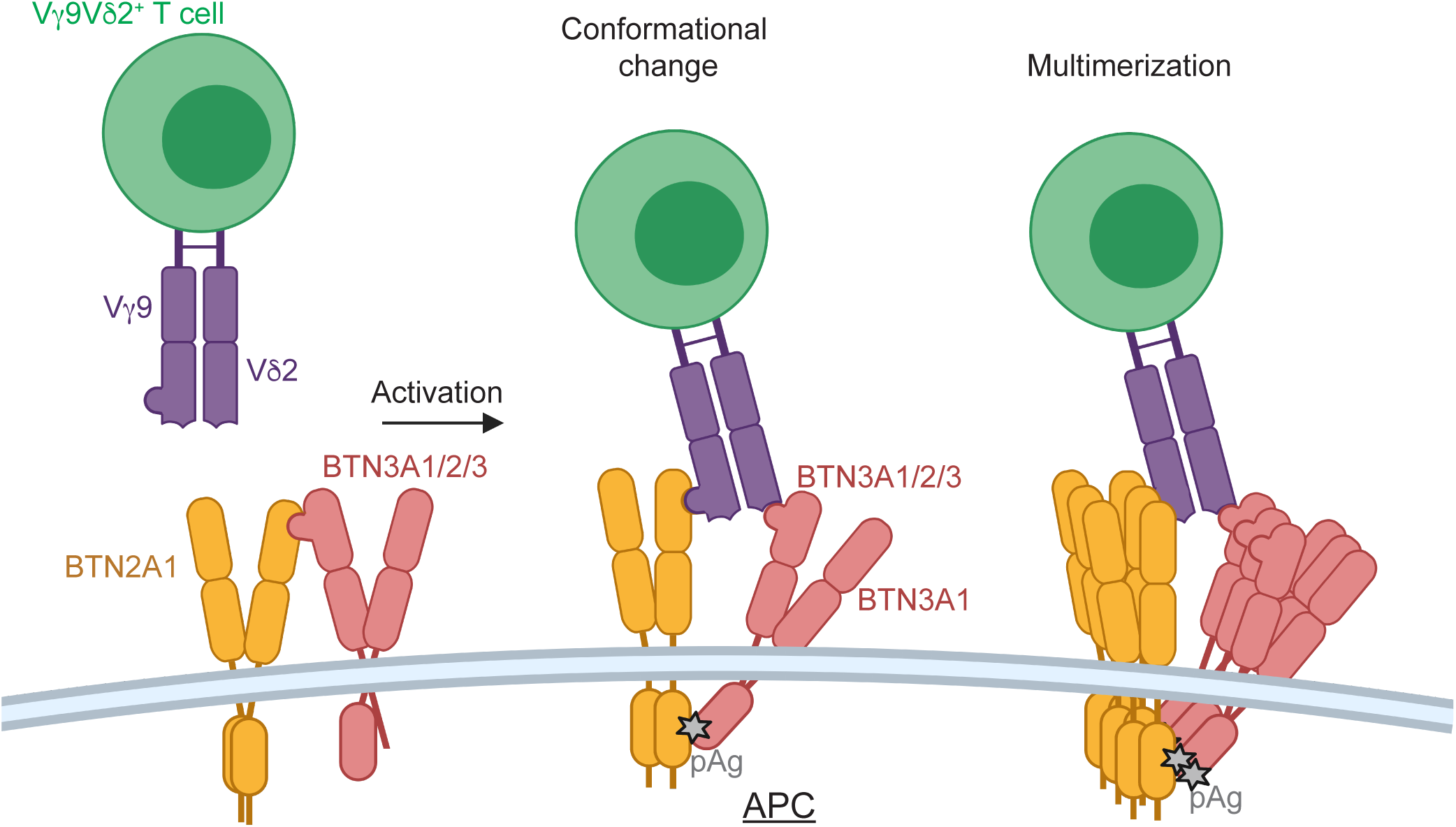
Proposed model(s) of Vγ9Vδ2^+^ TCR interacting with both components of the BTN2A1–BTN3A complex following a conformational change or multimerization of the BTN complex on APCs with activation. Created with BioRender.com.

**Extended Data Fig. 13.**
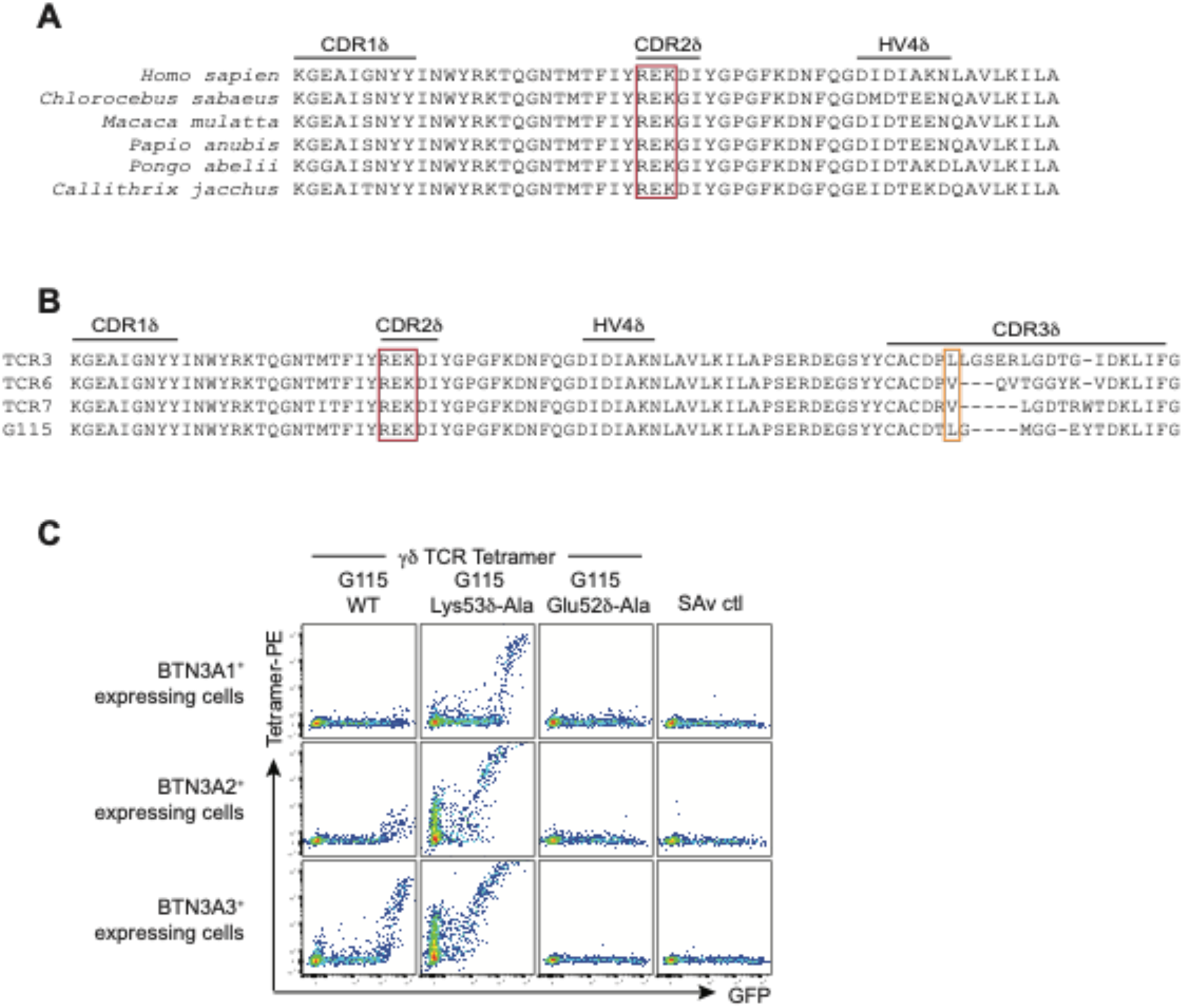
(**A**) Residues in the CDR2δ [Arg51δ, Glu52δ and Lys53δ; red box] are conserved between human and primate orthologues of TRDV2. TRDV2 alignment from ensembl.org. (**B**) Arg51δ, Glu52δ and Lys53δ are conserved in the CDR2δ between pAg-reactive Vγ9Vδ2 TCRs. Red box, conserved CDR2δ residues we have identified as being important for the BTN3A1 interaction. Orange box, semi-conserved CDR3δ residue 97 implicated in pAg-responsiveness. (**C**) Mouse 3T3 fibroblasts expressing BTN3A1, BTN3A2 or BTN3A3 were probed for binding by Vγ9Vδ2 TCR PE-tetramers ‘G115 WT’, ‘G115 Lys53δ-Ala’ or ‘G115 Glu52δ-Ala’, or SAv-PE control. Representative one of two independent experiments.

**Extended Data Fig. 14.**
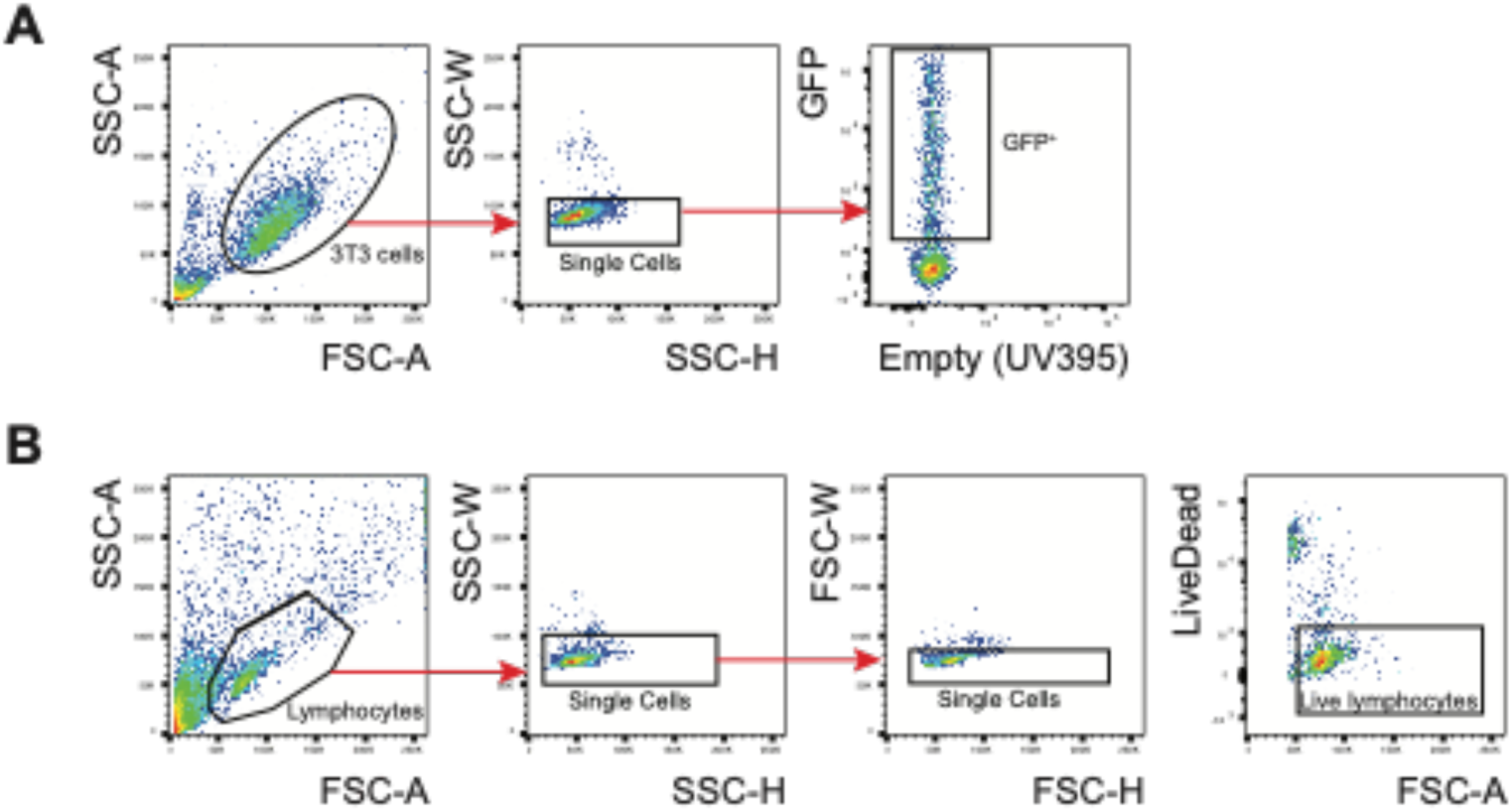
Flow cytometry gating strategy to identify (**A**) transfected NIH-3T3 or HEK293T cells, or (**B**) lymphocytes in co-culture assays.

